# Depsipeptide nucleic acids: prebiotic formation, oligomerization, and self-assembly of a new candidate proto-nucleic acid

**DOI:** 10.1101/2020.09.01.278838

**Authors:** David M. Fialho, Suneesh C. Karunakaran, Katherine W. Greeson, Isaac Martínez, Gary B. Schuster, Ramanarayanan Krishnamurthy, Nicholas V. Hud

**Affiliations:** School of Chemistry and Biochemistry and Parker H. Petit Institute for Bioengineering and Bioscience, Georgia Institute of Technology, Atlanta, Georgia 30332, USA; NSF-NASA Center for Chemical Evolution, Atlanta, Georgia 30332, USA; Department of Chemistry, The Scripps Research Institute, La Jolla, California 92037, USA

## Abstract

The mechanism by which genetic polymers spontaneously formed on the early Earth is currently unknown. The RNA World hypothesis implies that RNA oligomers were produced prebiotically, but the demonstration of this process has proven challenging. Alternatively, RNA may be the product of evolution and some, or all, of its chemical components may have been preceded by functionally analogous moieties that were more readily accessible under plausible early-Earth conditions. We report a new class of nucleic acid analog, depsipeptide nucleic acid, which displays several properties that make it an attractive candidate for the first informational polymer to arise on the Earth. The monomers of depsipeptide nucleic acids can form under plausibly prebiotic conditions. These monomers oligomerize spontaneously when dried from aqueous solutions to form nucleobase-functionalized depsipeptides. Once formed, these depsipeptide nucleic acid oligomers are capable of complementary self-assembly, and are resistant to hydrolysis in the assembled state. These results suggest that the initial formation of primitive, self-assembling, informational polymers may have been relatively facile.

## Introduction

The prebiotic emergence of the first genetic polymers is a poorly understood phenomenon.^1^ The influential RNA World hypothesis suggests that pre-formed nucleosides or nucleotides, through some hitherto unclear mechanism, spontaneously polymerized to form RNA polymers.^2^ However, despite repeated attempts, demonstrating the formation of oligomeric RNA from mononucleotides in a plausibly prebiotic manner is still a formidable challenge.^3-4^ This, and other issues associated with the initial formation of the nucleosides, suggest that RNA is a product of evolution, and was preceded by a proto-RNA polymer that was similar in function but more easily accessed in a prebiotic manner.^5-9^

A number of proto-RNA nucleobases have been suggested that have greater propensities for self-assembly^10-13^ and are more reactive with ribose^10, 12, 14-17^ and other sugars^13, 18^ than adenine, guanine, uracil, or cytosine (the canonical RNA nucleobases). Similarly, it may be that chemical features and properties of the backbone of proto-RNA differed significantly from RNA. The backbone of RNA consists of two principal components: the ribose sugar and the phosphate linker. A number of studies have been performed on the formation and base-pairing properties of phosphodiester-linked nucleic acids with alternative sugars.^19-20^ The prebiotic feasibility of the phosphodiester linkage has long been disputed,^21^ although some recent studies offer solutions to some obstacles to prebiotic phosphorylation.^22-24^ A number of alternatives to the phosphodiester linkage have been suggested, including acetals,^25^ amides,^26-27^ phosphite esters,^28^ and thiophosphate esters.^29^

The carboxylic ester linkage, which arguably has more in common with the phosphodiester linkage than most other proposed alternatives, has been investigated for nucleic acid analogs, but has only been formed under non-prebiotic conditions with a carbodiimide condensing agent.^30-31^ In the context of proto-polypeptide formation, it has been found that oligoesters are formed readily from α-hydroxy acids under relatively mild drying conditions and without the use of a condensing agent.^32^ Furthermore, when α-hydroxy acids and α-amino acids are combined in wet-dry cycling reactions, depsipeptide oligomers (containing both ester and amide moieties) are readily formed.^33-34^ It may be that proto-nucleic acids were also formed with ester and/or amide linkages, which were, through a process of chemical and/or biological evolution, eventually replaced by the more optimal, but prebiotically less accessible, phosphodiester linkage.^6-8^

In this study, we describe a new class of plausibly prebiotic polymer, depsipeptide nucleic acid, based on nucleobase-functionalized α-hydroxy acid-α-amino acid heterodimers as the proto-nucleic acid monomers. These monomers can be formed under plausibly prebiotic conditions and oligomerize by esterification, without the use of a chemical condensing agent, to form depsipeptide nucleic acids. Furthermore, depsipeptide nucleic acid oligomers self-assemble and show backbone-dependent resistance to hydrolysis. This new class of oligomers has the potential to function as an informational base-pairing system, and the specific compounds used in this study serve as models for prebiotic polymer systems that may be ancestral to more advanced genetic systems.

### Prebiotic Synthesis of Depsipeptide Nucleic Acid Monomers

The prebiotic synthesis of nucleobase-functionalized compounds capable of oligomerization is described in the literature and has been discussed in a prebiotic context by Cleaves: nucleophilic nucleobases react readily with acrolein (a plausibly prebiotic compound^35^) to give, among other products, γ-nucleobase-functionalized aliphatic aldehydes.^36^ These aldehydes can act as substrates for cyanohydrin or α-amino nitrile formation, which are hydrolyzed to give γ-nucleobase-α-hydroxy acids or γ-nucleobase-α-amino acids respectively.^7^ This synthetic path was also proposed by Miller for the formation of γ-functionalized amino acids such as methionine.^37^ A prebiotic precedent for the formation of γ-nucleobase-functionalized aldehydes was also recently demonstrated by Rodriguez et al., who found that performing Miller-Urey-type electrolysis reactions in a reducing atmosphere above aqueous solutions containing nucleobases produced, as the major products, 1,4-addition products of nucleobases (canonical or noncanonical) with Michael acceptors, including acrolein.^38^

Accordingly, we attempted one-pot, aqueous syntheses of γ-nucleobase-functionalized cyanohydrins from the reactions of one canonical and two noncanonical plausibly prebiotic heterocycles, cyanuric acid (Cy), melamine (Mel), and adenine (Ad), with acrolein and with potassium ferrocyanide as the cyanide source. These bases were chosen because they are formed in the same model prebiotic reaction,^39^ and because cyanuric acid, with an acceptor-donor-acceptor hydrogen-bonding pattern, can form aqueous hexad-based supramolecular assemblies by Watson-Crick-like pairing with derivatives of melamine (donor-acceptor-donor),^11^ as well as supramolecular polymers with homo-adenine (donor-acceptor) RNA or PNA oligomers.^40-42^ Ferrocyanide was chosen as the cyanide source because iron(II) cyanide complexes have been noted as compounds that were likely present on the early Earth (due to the inferred abundance of iron(II) and cyanide,^43^ or due to their direct detection in meteorites^44^) which can act as a slowly-releasing cyanide reservoir. In this regard, ferrocyanide has been successfully used in aqueous Strecker reactions as a “green” alternative to methods that employ more toxic cyanide sources.^45^

The aqueous reactions of cyanuric acid and adenine with acrolein and potassium ferrocyanide produced the corresponding nucleobase-functionalized cyanohydrins (Supplementary Figs 5 and 7), and acidic hydrolysis of these crude reaction mixtures gave a cyanuric acid-functionalized hydroxy acid, Cy^HA^ (Fig. 1a, Supplementary Fig. 6), and an adenine-functionalized hydroxy acid, Ad^HA^ (Fig. 1b, Supplementary Fig. 8), respectively. Melamine, when combined with potassium ferrocyanide, precipitates to form an insoluble salt; therefore, only a small amount of a melamine-acrolein adduct is formed (Supplementary Fig. 9). However, when acetone cyanohydrin is used as a cyanide source, melamine remains soluble, and a large amount of melamine-acrolein adduct is formed, but only a small amount of melamine-functionalized cyanohydrin is formed (Supplementary Fig. 11), which is likely due to sequestration of the aldehyde substrate by cyclization of the melamine-acrolein adduct (Supplementary Fig. 14).

**Figure 1.**
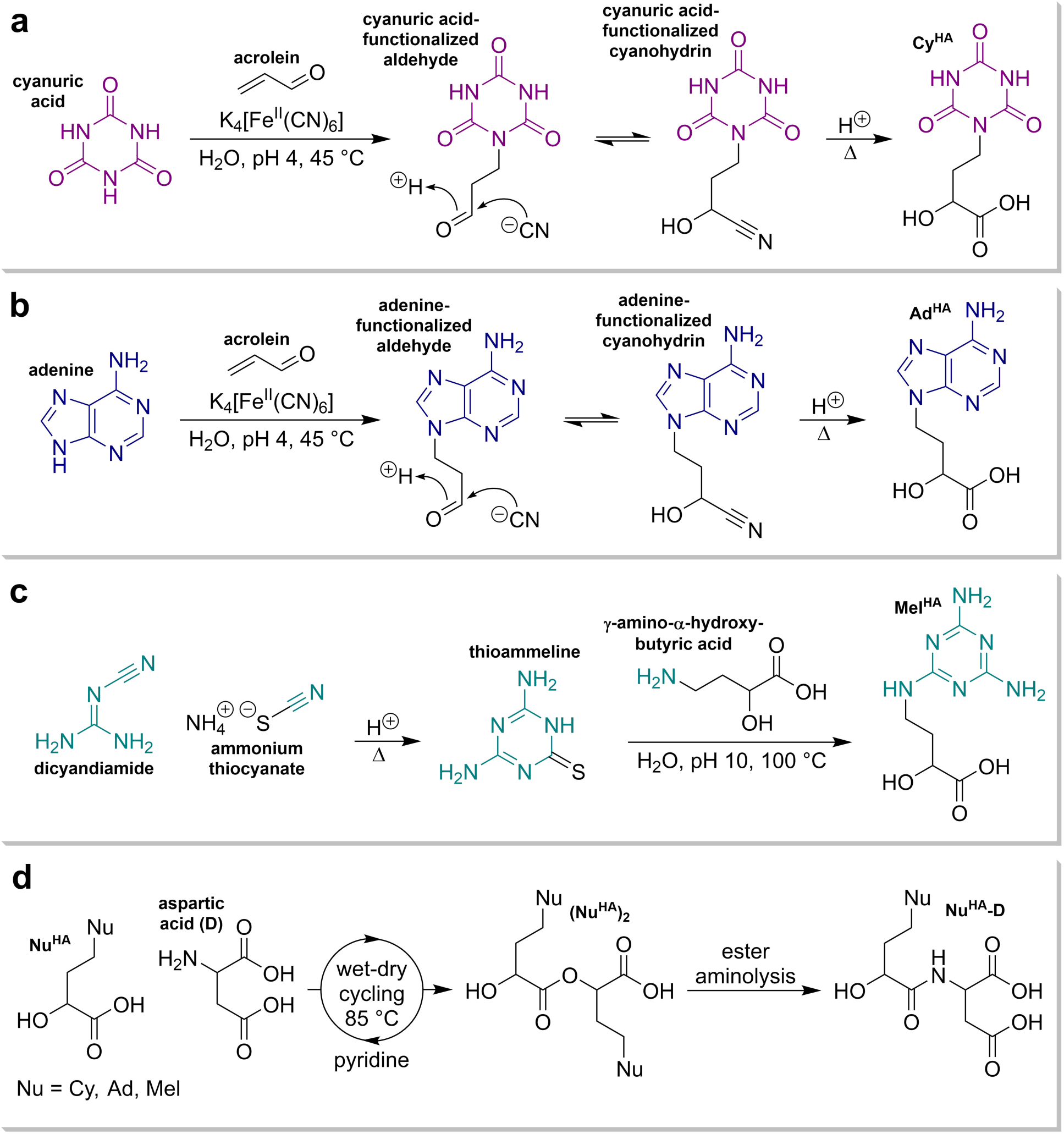
Prebiotic syntheses of γ-nucleobase-functionalized α-hydroxy acids and nucleobase-functionalized α-hydroxy acid-α-amino acid heterodimers. **a**, The reaction of cyanuric acid with acrolein in the presence of potassium ferrocyanide gives the γ-cyanuric acid-functionalized aldehyde, which reacts with cyanide to form the corresponding cyanohydrin. Upon acidic hydrolysis, Cy^HA^ is formed. **b**, The reaction of adenine with acrolein in the presence of potassium ferrocyanide gives the γ-adenine-functionalized aldehyde, which reacts with cyanide to form the corresponding cyanohydrin. Upon acidic hydrolysis, Ad^HA^ is formed. **c**, The reaction of dicyandiamide with thiocyanate gives thioammeline, which reacts further with α-hydroxy-γ-aminobutyric acid by nucleophilic aromatic substitution to form Mel^HA^. **d**, When subjected to wet-dry cycles at 85 °C from an aqueous solution containing pyridine, nucleobase-functionalized hydroxy acids condense with aspartic acid (via an ester homodimer intermediate) to form nucleobase-functionalized heterodimers.

Although a melamine-functionalized cyanohydrin was not formed in large amounts, the prebiotic formation of aminoalkyl nucleobases can also occur through nucleophilic aromatic substitution of sulfur-substituted heterocycles.^17, 46^ Thioammeline, which readily forms from the reaction of dicyandiamide (the dimer of cyanamide) with thiocyanate,^47^ reacts with amines to form melamine derivatives.^48^ The reaction of thioammeline with α-hydroxy-γ-aminobutyric acid (itself formed prebiotically from acrolein by the Cleaves-Miller path to γ-functionalized amino acids and hydroxy acids^37^) gives the melamine-functionalized hydroxy acid, Mel^HA^ (Fig. 1c, Supplementary Fig. 15).

We next investigated the prebiotic co-oligomerization of nucleobase-functionalized hydroxy acids (Nu^HA^; Nu = Cy, Ad, Mel) with aspartic acid (abbreviated as D), an amino acid which is formed concomitantly with cyanuric acid, adenine, and melamine.^39^ It was previously found that the co-oligomerization of amino acids and hydroxy acids by wet-dry cycling produces oligopeptides with N-terminal hydroxy acid residues.^49^ We reasoned that wet-dry cycling of Nu^HA^ species with aspartic acid would produce short oligomers of the form Nu^HA^-D_n_, and aspartic acid was chosen for this model system so that the depsipeptide oligomers would contain ionizable residues to maintain solubility in water. The formation of Nu^HA^-D_n_ oligomers proceeds by an initial acid-catalyzed esterification of Nu^HA^ to form (Nu^HA^)_2_ species, which are then aminolyzed by aspartic acid to form Nu^HA^-D (Fig. 1d). This heterodimer can be further elongated by esterification of the C-terminus (at either the α- or β-carboxylic acid of aspartic acid) and subsequent ester aminolysis by another molecule of aspartic acid.

Subjecting any of the Nu^HA^ compounds (prepared by conventional organic synthesis for these experiments; see Section II of the Supplementary Information) and one equivalent of aspartic acid to wet-dry cycling at 85 °C for 7 cycles (i.e., one instance of rehydration per day, followed by drying at 85 °C for 24 hours) gives only an insignificant amount of product (Supplementary Figs 16-20). We attribute the poor yields of these reactions to the physical state of the dried residues, which appear as white powders. In this solid, microcrystalline state, diffusion cannot occur, and reactive moieties therefore do not encounter each other. We reasoned that the addition of a volatile, mildly basic co-solvent might maintain hydroxy acid solubility as the sample dried (without changing the pH significantly) to eventually give a glass-like state in which diffusion could occur, thereby enabling the reaction. Because aromatic compounds, including heteroaromatics, are abundant in carbonaceous meteorites^50-51^ and are commonly produced in model prebiotic reactions,^52-53^ we selected pyridine as a model heterocyclic compound for these wet-dry cycling reactions. Including 8 equivalents of pyridine in the reaction of Cy^HA^ with aspartic acid changed the physical state of the dried pellet to a glass-like semi-solid. Comparison of the UV-LC/MS chromatograms of these reactions showed that the addition of pyridine significantly increased the formation of Cy^HA^-D (Supplementary Fig. 16). The physical state of the reactions of Ad^HA^ and Mel^HA^ with aspartic acid changed slightly by the addition of pyridine, but the formation of Ad^HA^-D and Mel^HA^-D remained low (Supplementary Figs 17 and 18). However, when wet-dry cycling reactions were performed with mixtures of Cy^HA^ and one of its pairing partners (Ad^HA^ or Mel^HA^) with aspartic acid and 8 equivalents of pyridine, the formation of Ad^HA^-D and Mel^HA^-D improved to amounts comparable with those found for Cy^HA^-D (Supplementary Figs 19 and 20).

### Prebiotic Oligomerization of Model Proto-Nucleic Acid Monomers

To further investigate the potential for oligomerization of this proto-nucleic acid system, we synthesized and isolated (*S,S*)-Cy^HA^-D (Fig. 2a), (*S,S*)-Ad^HA^-D (Fig. 2b), and (*S,S*)-Mel^HA^-D (Fig. 2c) by conventional organic synthesis. These compounds, which serve as model proto-nucleic acid monomers, are expected to oligomerize by esterification to form depsipeptides when subjected to dry-down conditions at elevated temperature and mildly acidic pH (Fig. 3a). This esterification reaction can proceed either directly at the α-carboxylic acid of the aspartic acid moiety or through intermediate lactonization to form morpholine-2,5-diones, to produce a backbone with a 6-atom repeat. This backbone may assume conformations that facilitate self-assembly by presenting nucleobases and solubilizing carboxylates on alternating sides of the depsipeptide chain, and is geometrically similar to previously studied oligopeptide nucleic acid systems that were found to support base pairing.^27, 54-55^ Competing β-esterification produces a 7-atom segment in the backbone of the nascent informational oligomer. This segment may also access conformations that permit self-assembly. It is possible for α-esters and β-esters to be present in the same oligomer in either a linear or a branched manner.

**Figure 2.**
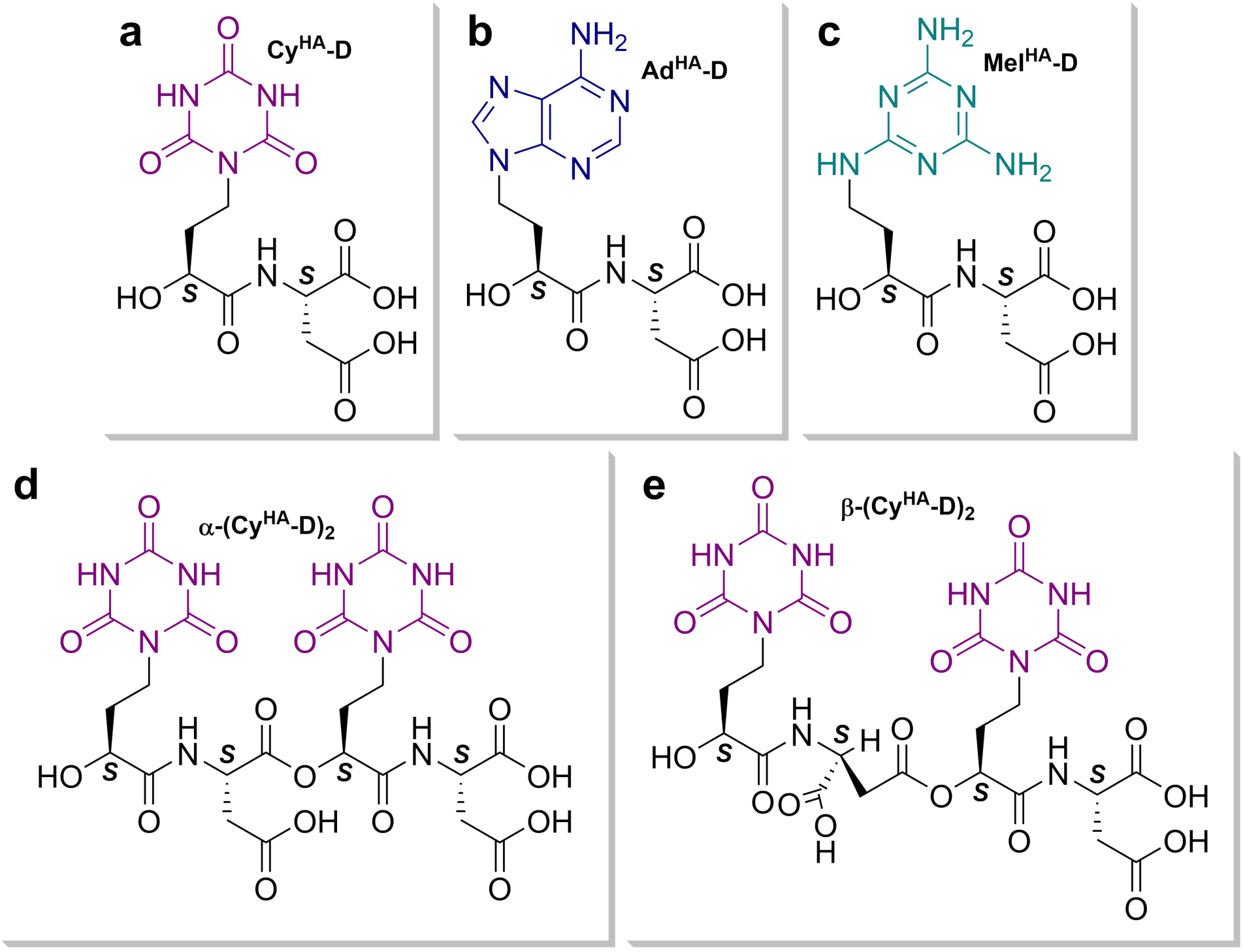
Structures of the three model proto-nucleic acid monomers and two model dimers used in this study. **a**, (*S,S*)-Cy^HA^-D, a heterodimer of Cy^HA^ and aspartic acid. **b**, (*S,S*)-Ad^HA^-D, a heterodimer of Ad^HA^ and aspartic acid. **c**, (*S,S*)-Mel^HA^-D, a heterodimer of Mel^HA^ and aspartic acid. **d**, α-(*S,S,S,S*)-(Cy^HA^-D)_2_, a dimer of Cy^HA^-D in which the two units are linked by esterification of the α-carboxylic acid of the first aspartic acid residue. **e**, β-(*S,S,S,S*)-(Cy^HA^-D)_2_, a dimer of Cy^HA^-D in which the two units are linked by esterification of the β-carboxylic acid of the first aspartic acid residue. Not that in the nomenclature of the current study, the model proto-nucleic acid monomers are themselves hydroxy acid-amino acid heterodimers, and the model dimers are heterotetramers.

**Figure 3.**
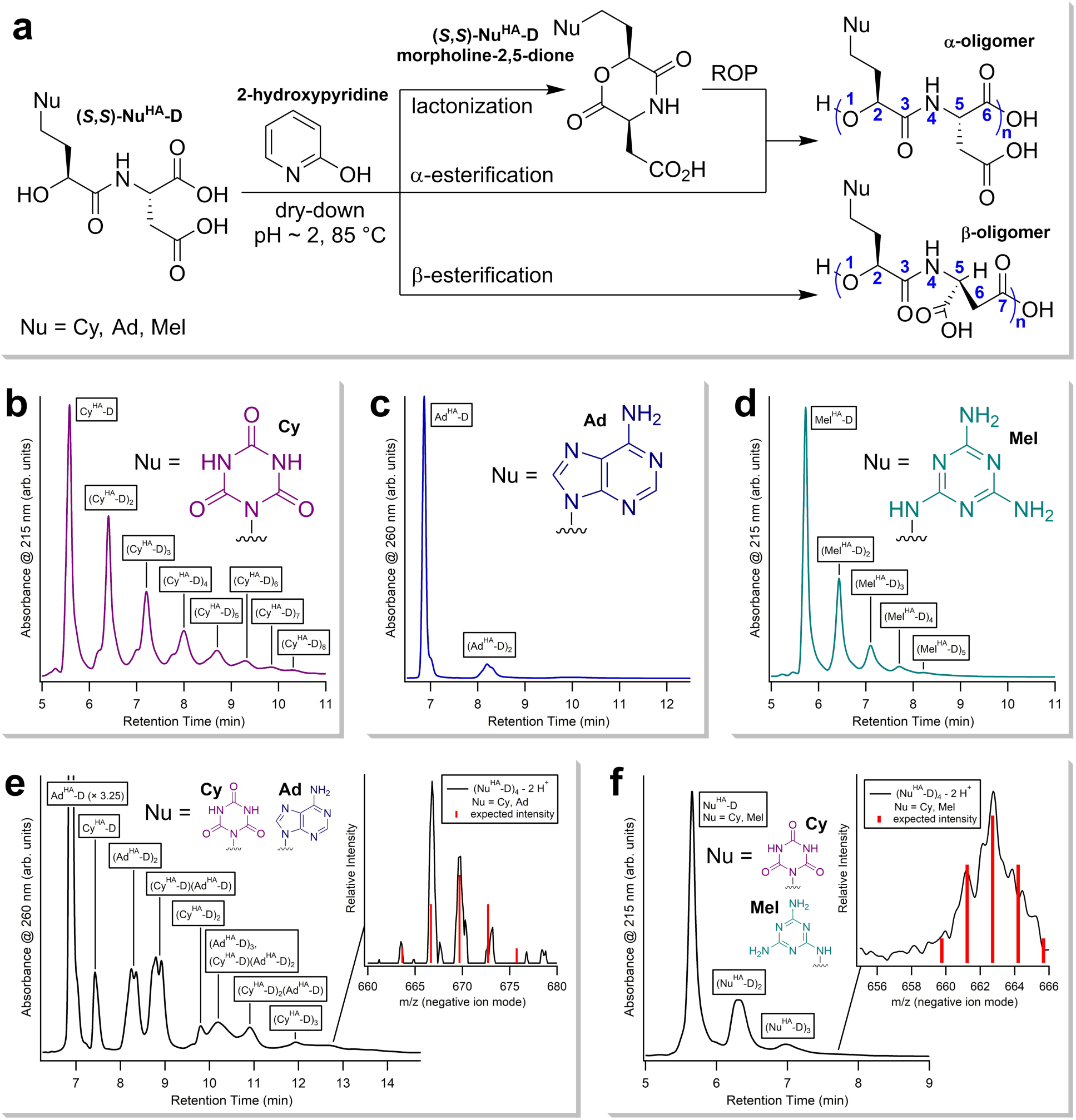
UV-LC/MS analysis of the oligomerization reactions of Cy^HA^-D, Ad^HA^-D, and Mel^HA^-D at 85 °C with 1 equivalent of 2-hydroxypyridine after 8 weeks. **a**, Prebiotic oligomerization of model monomers. When dried from an acidic aqueous solution containing a heterocyclic additive, oligomerization by esterification occurs. Two backbone linkages are possible: the α-linkage, formed by ring-opening polymerization of the morpholine-2,5-dione form of the monomer, or by direct α-esterification, and the β-linkage, formed by direct β-esterification. **b**, UV chromatogram of Cy^HA^-D oligomerization. **c**, UV chromatogram of Ad^HA^-D oligomerization. **d**, UV chromatogram of Mel^HA^-D oligomerization. **e**, UV chromatogram of mixed oligomerization of 1:1 Cy^HA^-D:Ad^HA^-D scaled by a factor of 3.25 relative to the height of the Ad^HA^-D peak. Oligomers of mixed sequence up to 4 units in length are detectable (inset) with abundances slightly offset from those predicted by a binomial distribution with equal probabilities of incorporation of each monomer. **f**, UV chromatogram of mixed oligomerization of 1:1 Cy^HA^-D:Mel^HA^-D. Oligomers of mixed sequence up to 4 units in length are detectable (inset) with abundances similar to those predicted by a binomial distribution with equal probabilities of incorporation of each monomer. In all cases, species are labeled according to the masses identified.

As with the coupling of the nucleobase-functionalized hydroxy acids to aspartic acid, the addition of a small heterocyclic compound increases the efficiency of oligomerization of the model proto-RNA monomers. For these dry-down oligomerization reactions, 2-hydroxypyridine, 2-mercaptopyridine, pyridine, and pyrimidine were assessed as additives. Pyridine derivatives, including 2-hydroxypyridine, have been identified in meteorites.^51, 56^ Pyrimidine may be produced prebiotically from acrylonitrile.^57^ 2-Hydroxypyridine (and, by analogy, 2-mercaptopyridine) was chosen for its propensity to act as a bifunctional catalyst in ester aminolysis reactions,^58^ which may also enhance the extent of the esterification reaction (Supplementary Fig. 21). Unlike pyridine, 2-hydroxypyridine and 2-mercaptopyridine and non-volatile and non-basic. Therefore, pyrimidine was tested because it allows a reasonable comparison to be made with pyridine with regard to basicity: both compounds are liquids with boiling points close to 120 °C; however, pyrimidine is less basic than pyridine. Initial screenings were performed by drying the aqueous solutions of Nu^HA^-D compounds with heterocyclic additives and maintaining them in a dry state at 85 °C for one week. All four of the heterocycles tested showed some enhancement of oligomerization. For the volatile compounds pyridine and pyrimidine, the extent of oligomerization increased with higher loadings (Supplementary Figs 22 and 23). For the non-volatile compounds 2-hydroxypyridine and 2-mercaptopyridine, an optimum loading of one equivalent was determined. Reactions containing 2-mercaptopyridine showed the significant formation of degradation products resulting from amide hydrolysis and subsequent ester aminolysis (Supplementary Fig. 24). One equivalent of 2-hydroxypyridine was found to be effective at enhancing the extent of oligomerization without degradation (Supplementary Fig. 25). It was further found that longer reaction times gave greater extents of oligomerization, with the greatest conversion of monomer observed with one equivalent of 2-hydroxypyridine after 8 weeks (Supplementary Fig. 26). The UV-LC/MS analyses of these oligomerization reactions are shown in Figure 3b-f.

Of the three model proto-nucleic acid monomers studied, Cy^HA^-D oligomerized the most efficiently, with oligomers detected up to 8 units in length. Ad^HA^-D oligomerizes poorly; only dimers were detected. Mel^HA^-D oligomerizes with intermediate efficiency, with oligomers detected up to 5 units in length. In the most prebiotically relevant scenario, these monomers would be present in the same mixture so that oligomers of mixed sequence could be formed. When Cy^HA^-D was co-oligomerized with either of its potential pairing partners, Ad^HA^-D or Mel^HA^-D, mixed-sequence oligomers up to 4 units in length were detected. Assuming comparable ionization efficiencies, the ion abundances of the tetramers from these reactions approximate a binomial distribution in which incorporation of either monomer is equally likely. For the co-oligomerization of Cy^HA^-D with Mel^HA^-D, this binomial approximation is very accurate (Fig. 3f, inset). For the co-oligomerization of Cy^HA^-D with Ad^HA^-D, the distribution is skewed slightly to favor oligomers enriched in Cy^HA^-D (Fig. 3e, inset).

The results of these oligomerization reactions are similar to those found in the wet-dry cycling reactions of Cy^HA^, Ad^HA^, and Mel^HA^ with aspartic acid in that the cyanuric acid-functionalized monomer reacts more efficiently than the adenine- or melamine-functionalized monomers. This is likely a consequence of a diminished rate of esterification in the deposited state due to the microcrystallinity of compounds containing carboxylic acid residues and adenine or melamine moieties.

### Self-Assembly of Depsipeptide Nucleic Acids

With the confirmation of the ability of these monomers to oligomerize by esterification in a plausibly prebiotic manner, we next turned to molecular self-assembly studies to determine whether the monomers or oligomers form pairing systems in water. Derivatives of cyanuric acid and melamine have been shown to form hexad-based supramolecular assemblies in water.^11^ Each hexad contains three cyanuric acid units and three melamine units, with each nucleobase interacting with two complementary nucleobases to create a highly regular structure with three-fold rotational symmetry. These planar, hydrophobic hexads stack in water to form micron-length supramolecular fibers when one or both of the nucleobase substituents provides a solubilizing electrostatic charge (Fig. 4a).^11-12^ Evidence of similar structures has been presented for cyanuric acid and homo-adenine oligomers, in which a hexad motif has also been proposed, but with a less regular structure due to the reduced structure symmetry of adenine.^40-42^ Although this hexad mode of self-assembly is distinct from the familiar duplex mode of extant nucleic acids, information transfer by complementary base pairing is still possible, and the considerable stability of supramolecular structures associated with the hexad motif may have provided an evolutionarily advantage that, historically, allowed hexad-based self-assembly to precede duplex-based self-assembly.

**Figure 4.**
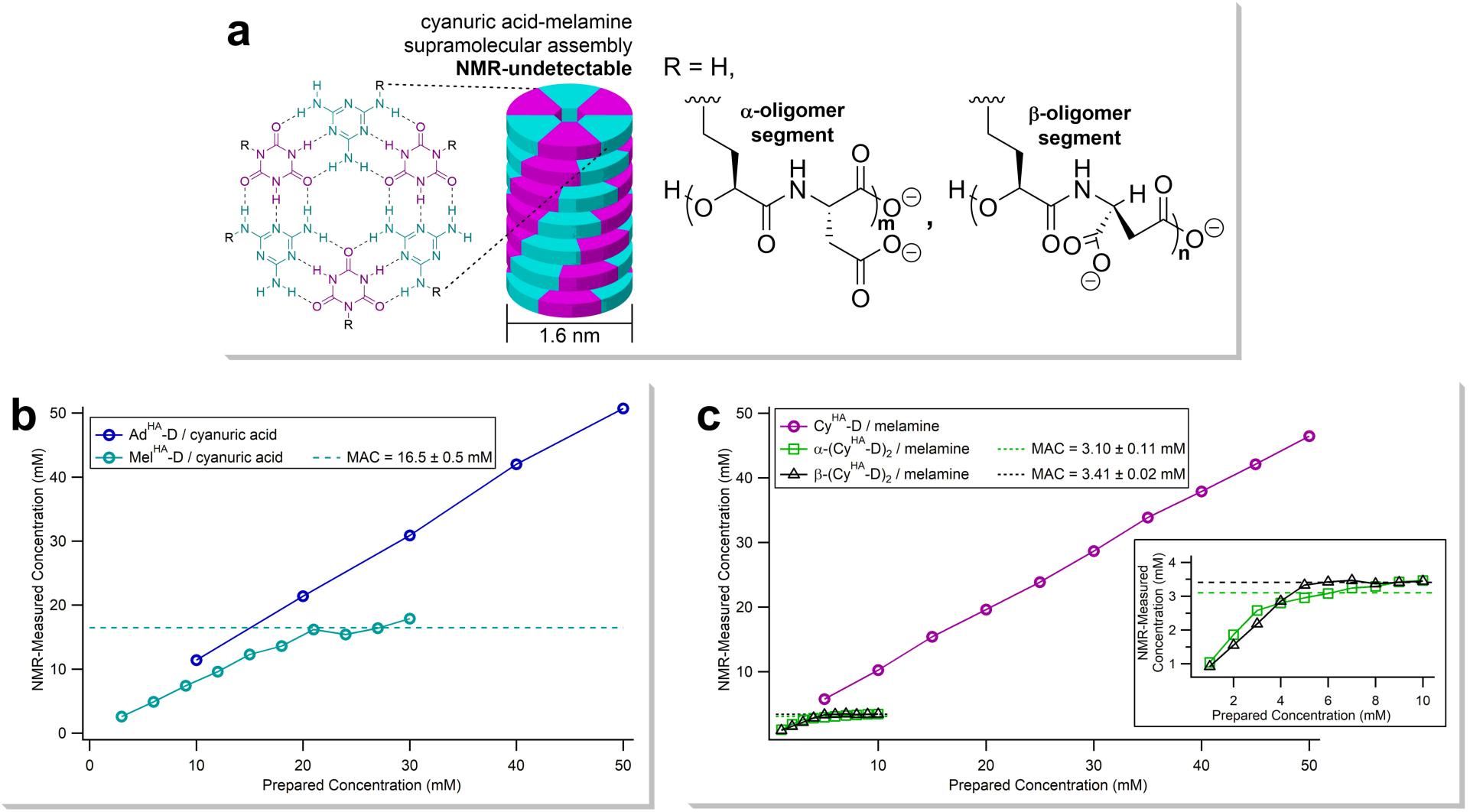
^1^H NMR analysis of the self-assembly of cyanuric-melamine systems and cyanuric acid-adenine systems. **a**, When cyanuric acid derivatives are incubated in water with melamine derivatives (or adenine derivatives) under certain conditions of temperature, pH, and ionic strength, self-assembly can occur to form supramolecular fibers. This supramolecular assembly forms after a critical concentration, the minimum assembly concentration (MAC), has been reached. Unassembled molecules remain detectable by NMR, but assembled molecules do not, allowing the MAC to be directly measured. All measurements were taken at 25 °C. **b**, Self-assembly of Ad^HA^-D or Mel^HA^-D with one equivalent of cyanuric acid in D_2_O, pD 7.5, with MgCl_2_ 25 mM. Ad^HA^-D does not assemble below 50 mM; Mel^HA^-D assembles with a MAC of 16.5 mM. **c**, Self-assembly of Cy^HA^-D, α-(Cy^HA^-D)_2_, or β-(Cy^HA^-D)_2_ with melamine in D_2_O, pD 6.5. Cy^HA^-D was incubated with one equivalent of melamine, while α-(Cy^HA^-D)_2_ and β-(Cy^HA^-D)_2_, with two cyanuric acid moieties for each, were incubated with two equivalents of melamine. Cy^HA^-D does not assemble below 50 mM; α-(Cy^HA^-D)_2_ and β-(Cy^HA^-D)_2_ assemble with MAC values of 3.10 mM and 3.41 mM, respectively (inset).

**Figure 5.**
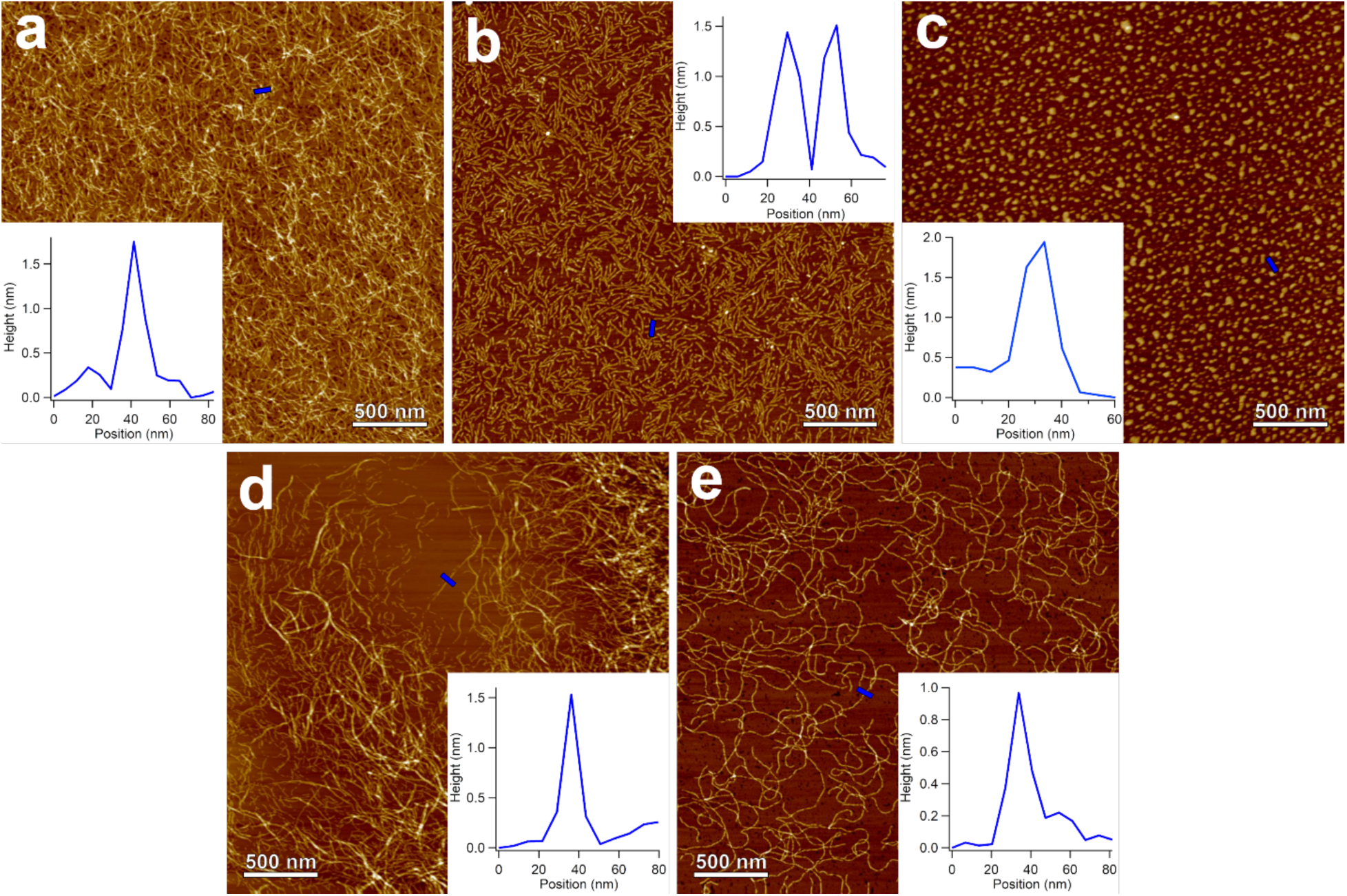
Analysis of the supramolecular assembly of the oligomers of Cy^HA^-D and Mel^HA^-D by atomic force microscopy. **a**, Topographical image of oligo-(Cy^HA^-D) paired with melamine, pD 6.5. In this panel, and others, insets show fiber height measurements taken across blue bar. **b**, Topographical image of oligo-(Mel^HA^-D) oligomers paired with cyanuric acid, pD 6.5. **c**, Topographical image of co-oligomers of Cy^HA^-D and Mel^HA^-D with no externally added underivatized pairing heterocycle, pD 7.5, MgCl_2_ 25 mM. **d**, Topographical image of α-(Cy^HA^-D)_2_ paired with melamine, pD 6.5. **E**, Topographical image of β-(Cy^HA^-D)_2_ paired with melamine, pD 6.5.

**Figure 6.**
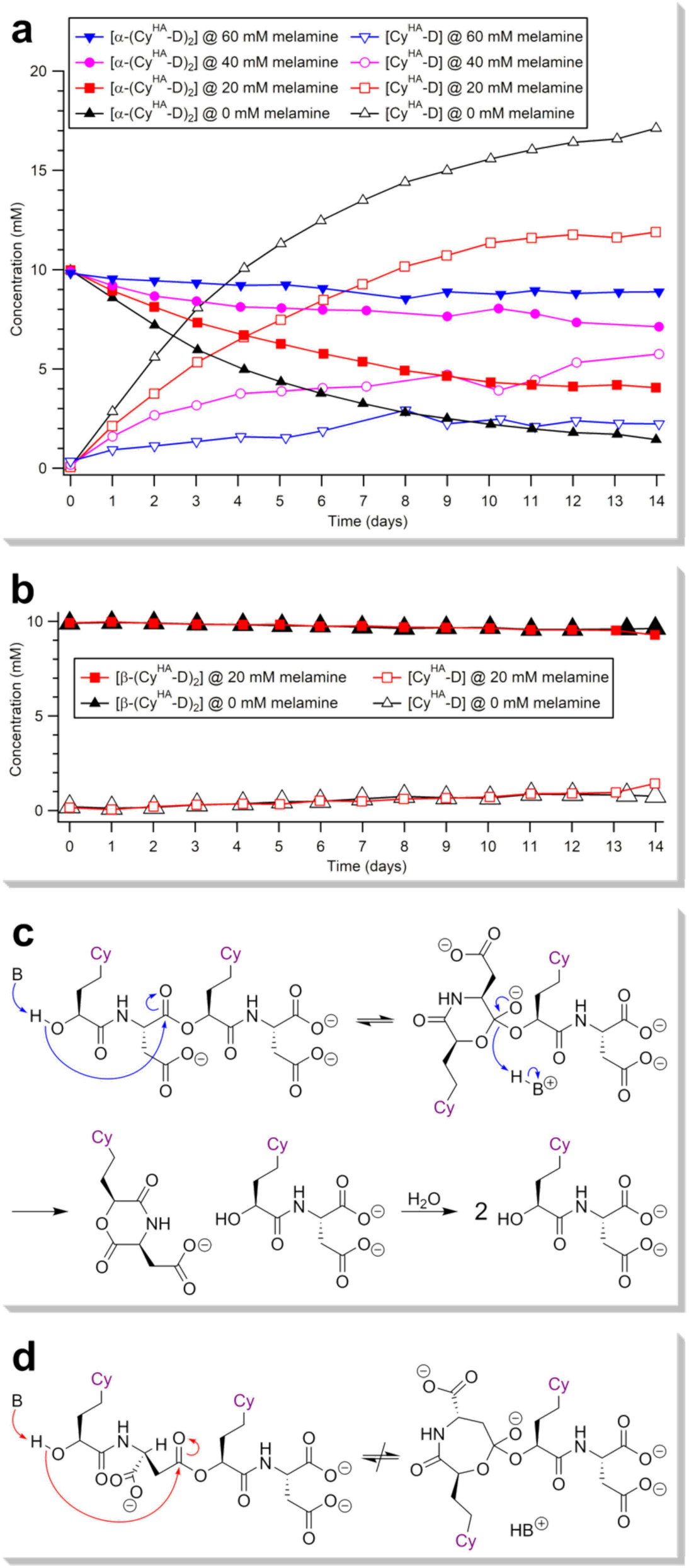
Hydrolysis of (Cy^HA^-D)_2_ at pD 6.5 and its inhibition by supramolecular assembly with melamine. **a**, Hydrolysis of α-(Cy^HA^-D)_2_ with varying concentrations of melamine. **b**, Hydrolysis of β-(Cy^HA^-D)_2_ with or without melamine at 20 mM. **c**, The proposed mechanism of hydrolysis of α-(Cy^HA^-D)_2_. Base-catalyzed intramolecular attack on the ester moiety produces a 6-membered cyclic intermediate which ejects one equivalent of Cy^HA^-D to form the morpholine-2,5-dione of Cy^HA^-D. This morpholine-2,5-dione rapidly hydrolyzes to give the second equivalent of Cy^HA^-D. **d**, An analogous mechanism of hydrolysis for β-(Cy^HA^-D)_2_ is unlikely to occur, as the formation of a 7-membered cyclic intermediate is required. All reported concentrations were measured by UV-LC/MS monitoring at 215 nm and were normalized according to the formula [(Cy^HA^-D)_2_] + 1/2[Cy^HA^-D] = 10 mM.

With charged backbone moieties, stacked-hexad supramolecular fibers remain soluble; however, their tumbling rates in solution are so slow that their ^1^H NMR signals broaden to baseline, leaving only the remaining unassembled molecules detectable. This phenomenon can be exploited to determine a minimum assembly concentration (MAC) that is a measure of the propensity of a given system to self-assemble: low MAC values indicate a strong propensity for self-assembly. Using this technique,^59-60^ MAC values were determined in D_2_O at 25 °C, pD 7.5, with MgCl_2_ 25 mM. These conditions were chosen based on previously determined trends in supramolecular assembly with pH (or pD)^11, 60^ and on the dependence of supramolecular assembly on divalent metal cations.^40^ Under these conditions, Mel^HA^-D forms long, soluble assemblies with cyanuric acid with a MAC of 16.5 mM (Fig. 4b). However, Ad^HA^-D did not assemble with cyanuric acid (Fig. 4b). Cy^HA^-D assembles with melamine under these conditions; however, these supramolecular aggregates precipitate from solution (Supplementary Fig. 27). No assembly was detected for Cy^HA^-D with Mel^HA^-D or Ad^HA^-D (Supplementary Figs 29 and 31).

Although the monomers did not consistently show propensity for self-assembly, this does not preclude the oligomers from undergoing self-assembly. To study this, the dimers of Cy^HA^-D (the monomer which oligomerized to the greatest extent) were isolated. As expected, two main species were formed: the α-ester, α-(Cy^HA^-D)_2_ (Fig. 2d), and the β-ester, β-(Cy^HA^-D)_2_ (Fig. 2e). The identities of these compounds were assigned based on the ^1^H NMR chemical shifts of the α- and β-protons of the aspartic acid residues, which shift downfield when the adjacent carboxylic acid residues are esterified (Supplementary Fig. 3). Compounds α-(Cy^HA^-D)_2_ and β-(Cy^HA^-D)_2_ were isolated in a relative yield of 2:1 (respectively), suggesting that the mechanism of oligomerization by lactonization and nucleophilic ring-opening was operative, but not exclusively (Fig. 3a).

For either of these dimers to self-assemble in the stacked-hexad motif, the cyanuric acid residues must point in the same direction (into the assembly), while the solubilizing carboxylate residues must face opposite the cyanuric acid residues (into the solvent). These criteria are met in the fully extended backbone conformation of α-(Cy^HA^-D)_2_, where all dihedral angles are 180°. However, for β-(Cy^HA^-D)_2_, these criteria can only be met with a distorted backbone conformation containing at least three bonds with dihedral angles deviating from 180°, potentially introducing steric strain. Nevertheless, α-(Cy^HA^-D)_2_ and β- (Cy^HA^-D)_2_ both form assemblies with melamine: at pD 6.5, with no divalent metal ion present, MAC values of about 3 mM were determined for both dimers (Fig. 4c, inset). These slightly modified conditions were employed to preemptively attenuate ester hydrolysis, which may be accelerated by slightly basic pH and by divalent metal cations.^61^ The product of this potential hydrolysis, Cy^HA^-D, did not assemble (or precipitate) with melamine under these conditions (Fig. 4c). These positive results for the dimers of Cy^HA^-D suggest that both 6-atom and 7-atom depsipeptide backbone segments can adopt conformations that permit assembly. No assemblies of either α-(Cy^HA^-D)_2_ or β-(Cy^HA^-D)_2_ with adenine were detected at concentrations up to 20 mM under these conditions (Supplementary Fig. 33).

The assemblies were next investigated by atomic force microscopy (AFM). Samples of oligomerized Cy^HA^-D, Mel^HA^-D, and Ad^HA^-D (dry state, 1 eq. 2-hydroxypyridine, 85 °C, two weeks) were incubated with their complement (either melamine or adenine for Cy^HA^-D oligomers, or cyanuric acid for oligomers of Mel^HA^-D and Ad^HA^-D) in D_2_O at pD 6.5; conditions for which no assembly was observed by ^1^H NMR spectroscopy for monomeric Cy^HA^-D, Ad^HA^-D, or Mel^HA^-D (Fig. 4c, Supplementary Figs 28-31). The cyanuric acid-melamine system shows a stronger propensity for self-assembly than the cyanuric acid-adenine system. While no fibers were detected for oligo-(Ad^HA^-D) with cyanuric acid (Supplementary Fig. 42), long fibers are formed from the assembly of oligo-(Cy^HA^-D) with melamine (Fig. 5a) or from the assembly of oligo-(Mel^HA^-D) with cyanuric acid (Fig. 5b). The heights of these assemblies (Fig. 5a,b, insets) match the predicted value for the stacked-hexad motif of 1.6 nm (Fig. 4a).

Co-oligomers formed from Cy^HA^-D and Mel^HA^-D (dry state, 1 eq. 2-hydroxypyridine, 85 °C, two weeks), assembled in the absence of an underivatized base at pD 7.5 with MgCl_2_ 25 mM, mostly forming nonlinear aggregates (Fig. 5c). Assuming that this assembly still occurs by the stacked-hexad motif, these aggregates contain six charged backbones whose electrostatic repulsion may inhibit the formation of long fibers. The heights of the interspersed short linear assemblies are slightly greater than 1.6 nm (Fig. 5c, inset), possibly also due to the presence of six backbones in one assembly.

Long fibers could also be observed for both α-(Cy^HA^-D)_2_ and β-(Cy^HA^-D)_2_ with melamine at pD 6.5 (with no divalent metal cation). The 1.6 nm height (Fig. 5d, inset) of the linear assemblies formed from α-(Cy^HA^-D)_2_ with melamine (Fig. 5d) is consistent with a stacked-hexad motif; however, assemblies formed from β-(Cy^HA^-D)_2_ with melamine (Fig. 5e) showed a height of 1.0 nm (Fig. 5e, inset), which could indicate a distorted or altogether different mode of self-assembly.

### Resistance of Depsipeptide Nucleic Acids to Hydrolysis

Orgel suggested that the hydrolytic instability of the ester bond diminishes its viability as a backbone linkage for proto-nucleic acids.^8^ We performed hydrolysis studies using the dimers α-(Cy^HA^-D)_2_ and β-(Cy^HA^-D)_2_ to test whether these esters would be too unstable to sustain an early informational polymer system. The dimers were incubated (10 mM, 25 °C in D_2_O, pD 6.5) both in the presence and absence of melamine, a gel-forming pairing partner. Over a period of 14 days, the concentrations of α-(Cy^HA^-D)_2_ or β-(Cy^HA^-D)_2_ and monomeric Cy^HA^-D were measured by UV-LC/MS (Fig. 6). It was found that the rate of hydrolysis of α-(Cy^HA^-D)_2_ diminished as the concentration of melamine increased (Fig. 6a), suggesting that α-(Cy^HA^-D)_2_ resists hydrolysis in the assembled state. Interestingly, the rate of hydrolysis of β-(Cy^HA^-D)_2_ is much slower than that of α-(Cy^HA^-D)_2_, and the effect of self-assembly with melamine is insignificant (Fig. 6b) on the timescale of this experiment.

The rate expressions for the hydrolysis of α-(Cy^HA^-D)_2_ or β-(Cy^HA^-D)_2_ in the presence of melamine are complex due to the existence of two equilibrating soluble phases (assembled and unassembled). However, a first-order approximation provides expressions for the half-lives that permit reliable comparison. In the absence of melamine, α-(Cy^HA^-D)_2_ hydrolyzes with a half-life of approximately 4.5 days. With 60 mM melamine present, this half-life increases more than 25-fold to about 100 days. On the other hand, β-(Cy^HA^-D)_2_, in the presence or absence of melamine, hydrolyzes with a half-life of approximately 200 days. (See Supplementary Table 1 for all rate constants.) The faster rate of hydrolysis of α-(Cy^HA^-D)_2_ compared to β-(Cy^HA^-D)_2_ can be attributed to a “backbiting” mechanism of hydrolysis (Fig. 6c).^62^ The terminal hydroxyl group of α-(Cy^HA^-D)_2_, acting as an intramolecular nucleophile, attacks the ester moiety to form a 6-membered ring intermediate, which ultimately produces the morpholine-2,5-dione of Cy^HA^-D by ejecting one equivalent of monomeric Cy^HA^-D. The morpholine-2,5-dione is then hydrolyzed into a second equivalent of Cy^HA^-D. This mechanism of hydrolysis is expected to be much less effective for β-(Cy^HA^-D)_2_, as intramolecular attack of the ester by the terminal hydroxyl group must proceed through a less favorable 7-membered ring intermediate (Fig. 6d).

## Discussion

The results presented here demonstrate that depsipeptide nucleic acids exhibit multiple properties required of a candidate proto-nucleic acid. The nucleobase-functionalized hydroxy acids are formed in a plausibly prebiotic manner. For adenine and cyanuric acid, the corresponding γ-nucleobase-functionalized cyanohydrins are formed in one pot. For melamine, a functionalized hydroxy acid is formed from prebiotic precursors by nucleophilic aromatic substitution of a sulfur-substituted heterocycle. Additionally, either of these reactions may be generalizable for the formation of other nucleobase-functionalized hydroxy acids and amino acids. Cy^HA^-D, Ad^HA^-D, and Mel^HA^-D, which serve as model monomers, are formed prebiotically by wet-dry cycling reactions of the Nu^HA^ species with aspartic acid in the presence of pyridine. These monomers oligomerize by esterification when dried from an aqueous solution with 2-hydroxypyridine at 85 °C at a mildly acidic pH. During this hot, dry phase, esterification is thermodynamically favorable and proceeds without the need for chemical activation. Furthermore, Cy^HA^-D and Mel^HA^-D, and their oligomers, self-assemble in water. Importantly, the species formed from the co-oligomerization of Cy^HA^-D and Mel^HA^-D also self-assemble, as is required for information transfer in a pre-RNA system. Finally, the hydrolysis rates of ester-linked dimers is strongly dependent on backbone geometry and greatly attenuated by association, which provides a mechanism for evolution by selection for enhanced hydrolytic stability.

It is important to note that prebiotic polymers formed by irreversible oligomerization may not have been amenable to chemical evolution. Reversible oligomerization provides an opportunity for error correction, which is an essential requirement, as the supply of monomers would otherwise be rapidly exhausted by “failed” sequences, preventing further evolution of the system.^63^ The oligomers described here are stabilized against hydrolysis by self-assembly but can also selectively be hydrolyzed to reform their component monomers. The proposed accelerated degradation by the backbiting mechanism of α-(Cy^HA^-D)_2_ compared with β-(Cy^HA^-D)_2_ has an important consequence for longer oligomers. If an α-ester linkage is present at the O-terminus of the depsipeptide (analogous to the N-terminus of a peptide), backbiting will occur until a β-linkage is encountered in the oligomer. Once a β-linkage is present at the O-terminus, the backbiting degradation mechanism is no longer operative, and any linkages (whether α or β) in the middle of the oligomer will be relatively stable to hydrolysis.

Over the past half-century, a great deal of experimental effort has been devoted to attempts to demonstrate a feasible prebiotic synthesis of canonical RNA nucleotides, the initial oligomerization of these nucleotides, and the non-enzymatic, template-directed replication of RNA oligomers. Despite this, no scenario yet exists that displays all of these phenomena.^64^ Our results suggest that alternative ancestral proto-nucleic acid monomers and oligomers, which can be formed and oligomerized under prebiotic conditions, hold the potential to address some of these difficulties. We hope this work will help inspire further exploration of this possibility.

## Conclusion

We have presented here a new candidate proto-nucleic acid, depsipeptide nucleic acid, which is accessible via plausible prebiotic synthesis from prebiotically available starting materials, and oligomerizes without the need for condensing agents. The depsipeptide nucleic acid displays propensity for self-assembly, and selective resistance to hydrolytic degradation upon self-assembly. These properties, combined with the reversible linker chemistry, create a system that is suitable for facilitating chemical evolution. The self-assembly of depsipeptide nucleic acid oligomers is enabled by the noncanonical nucleobases cyanuric acid and melamine, which may have preceded the extant canonical nucleobases in the evolution of RNA.^65^ The propensity of depsipeptide nucleic acids for oligomerization in the dry state and selective hydrolysis in the rehydrated state are inherent properties of esters. Like the nucleobase component of nucleic acids, the ester linkage, which is readily formed under prebiotic conditions, may have preceded the prebiotically challenging phosphodiester as the backbone linker of nucleic acids.

## Acknowledgments

The authors thank Martin C, Tyler Roche, and Prof. Charles Liotta for helpful discussions, and Gary Newnam and Leslie Gelbaum for technical assistance. This work was supported by the NSF and the NASA Astrobiology Program under the NSF Center for Chemical Evolution (CHE-1504217).

## Author contributions

N.V.H. and D.M.F. conceived the study. D.M.F., S.C.K., N.V.H., R.K., and G.B.S. designed the experiments. D.M.F., S.C.K., K.W.G., and I.M. performed the experiments. D.M.F. and S.C.K. analyzed the data. All authors discussed the results. D.M.F. and N.V.H. wrote the paper with input from R.K., G.B.S., S.C.K., K.W.G., and I.M. D.M.F. wrote the Supplementary Information.

