## Supplementary Information for "Depsipeptide nucleic acids: prebiotic formation, oligomerization, and self-assembly of a new candidate proto-nucleic acid"

### I. Materials and Methods

All reagents and solvents were purchased from Sigma-Aldrich/Millipore Sigma, VWR, or Fisher Scientific, and were used without further purification.

NMR analyses were performed on a Bruker Avance IIIHD 700 MHz NMR spectrometer or a Bruker Avance IIIHD 800 MHz NMR spectrometer. $^{1}$H NMR spectra were collected over 64 scans. $^{13}$C NMR spectra were collected over 1024 scans. 1D ROE spectra were collected over 256 scans. All NMR experiments were performed at 25 °C unless stated otherwise.

UV-LC/MS analyses were performed with an Agilent LC-MS system with a single-quadrupole mass spectrometer. The following chromatographic method was used:

Column: Waters XBridge Amide, 3.5 μm, 2.1 x 150 mm
Column Temperature: 25 °C
Flow Rate: 0.5 mL/min
Injection Volume: 1 μL
Mobile Phase A: Aqueous $NH_4HCO_2$ buffer, 10 mM, pH 9
Mobile Phase B: MeCN
Gradient:

| Time (min) | %A | %B |
|---|---|---|
| 0 | 10 | 90 |
| 4 | 25 | 75 |
| 11 | 35 | 65 |
| 11.2 | 40 | 60 |
| 12.8 | 40 | 60 |
| 13 | 10 | 90 |
| 18 | 10 | 90 |

Atomic force microscopy (AFM) images were obtained with a Nanoscope IIIa (Digital Instruments) in tapping mode using silicon tips (Vistaprobes, 48 N $m^{-1}$) on freshly cleaved mica. Samples were deposited onto mica in 3 μL aliquots, incubated in a humid atmosphere for 10 minutes, and then blown dry with a stream of nitrogen.

### II. Preparative Syntheses of Model Proto-Nucleic Acid Monomers

*Preparative Synthesis of* rac-*Cy^HA*

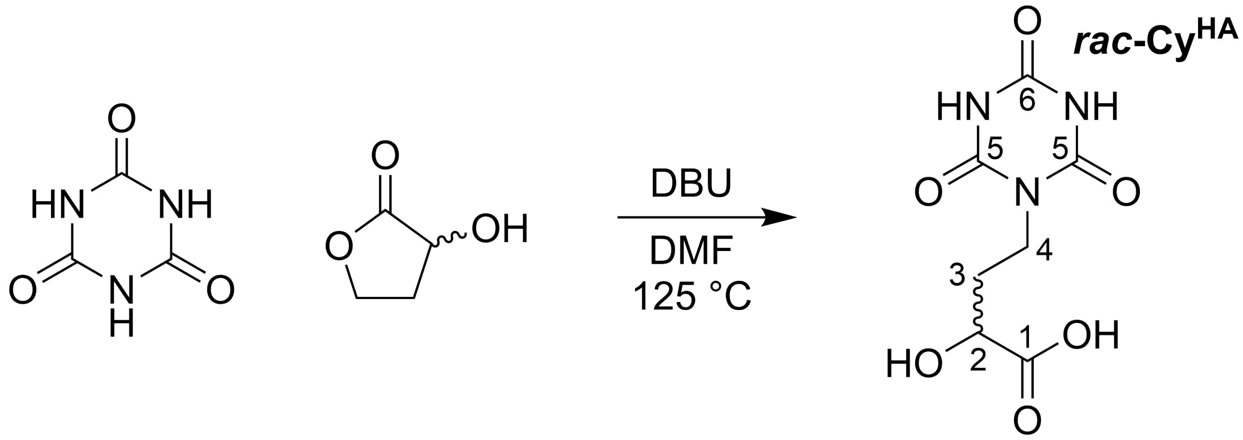

To 100 mL of DMF, 6.45 g of cyanuric acid (50 mmol) and 7.48 mL of DBU (1,8-diazabicyclo[5.4.0]undec-7-ene, 50 mmol) were added. The mixture was heated to 125 °C with rapid stirring. Once the cyanuric acid had fully dissolved, 1.95 mL of *rac*-α-hydroxy-γ-butyrolactone (25 mmol) were added dropwise. The reaction was stirred at 125 °C for 24 hours. The solution was then poured onto a watch glass and allowed to evaporate over several days, eventually producing a thick, brown oil above crystals of unreacted cyanuric acid. This material was suspended in 50 mL of water, and the suspension was centrifuged. The pellet was discarded and the supernatant was separated, in 5 mL portions, using a Teledyne Isco CombiFlash Rf+ purification system (monitoring by UV absorbance at 210 nm) with a RediSep C18Aq 150 g Gold column with a 100% water eluent. The fractions containing product, as detected by LC/MS, were pooled and reduced to about 10 mL with a rotary evaporator. To remove residual DBU, this solution was eluted over 25 g of Dowex 50WX8 cation exchange resin, hydrogen form, with 0.1 % aqueous formic acid solution. 100 mL were collected and the solvent was removed with a rotary evaporator to give white flakes of *rac*-$Cy^{HA}$ in its free acid form (1.73 g, 30%). $^1H$ NMR (700 MHz, $D_2O$): δ 4.33 (dd, 8.1 Hz, 4.1 Hz, 1H, $\mathbf{H^2}$), 3.96 (m, 2H, $\mathbf{H^4}$), 2.15 (m, 1H, $\mathbf{H^3}$), 2.01 (m, 1H, $\mathbf{H^{3'}}$). $^{13}C$ NMR (176 MHz, $D_2O$): δ 177.2 (**C1**), 150.9 (**C5**), 150.0 (**C6**), 68.1 (**C2**), 37.9 (**C4**), 31.0 (**C3**). HRMS (m/z): $[M+H]^+$ $C_7H_{10}O_6N_3$, theoretical mass: 232.0564, actual mass: 232.0568.

*Preparative Synthesis of* rac-*Cy^HA Isopropyl Ester*

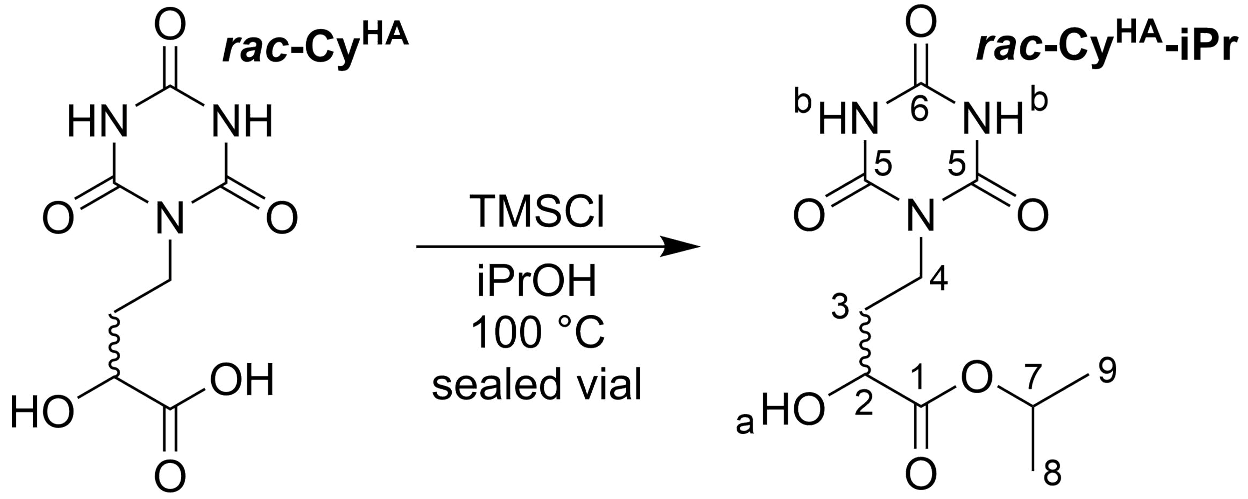

In a glass vial equipped with a stir bar, *rac*-Cy$^{HA}$ (116 mg, 0.5 mmol) was suspended in 1 mL isopropanol, and trimethylsilyl chloride (12.7 µL, 0.1 mmol) was added. The vial was sealed and the mixture was stirred rapidly at 100 °C for six hours, producing a solution. (WARNING: Heating solvents in sealed vessels above their boiling points creates a risk of explosion due to high pressure.)* The solvent was then evaporated first by blowing a stream of nitrogen onto the solution, and then by removing any residual solvent under vacuum, giving *rac*-Cy$^{HA}$ isopropyl ester as a white solid in quantitative yield (as determined by $^{1}$H NMR). The solid was left in the glass vial and taken to the next synthetic step without any further purification. $^{1}$H NMR (700 MHz, *d6*-DMSO): δ 11.37 (s, 2H, **H$^{b}$**), 5.43 (d, 5.5 Hz, 1H, **H$^{a}$**), 4.91 (m, 6.2 Hz, 1H, **H$^{7}$**), 4.04 (m, 1H, **H$^{2}$**), 3.74 (m, 2H, **H$^{4}$**), 1.89 (m, 1H, **H$^{3}$**), 1.76 (m, 1H, **H$^{3'}$**). $^{13}$C NMR (176 MHz, *d6*-DMSO): δ 173.5 (**C1**), 150.7 (**C5**), 149.1 (**C6**), 68.7 (**C2** or **C7**), 68.1 (**C2** or **C7**), 38.0 (**C4**), 32.1 (**C3**), 22.0 (**C8** or **C9**), 21.9 (**C8** or **C9**). HRMS (m/z): [M+H]$^{+}$ $C_{10}H_{16}O_6N_3$, theoretical mass: 274.1034, actual mass: 274.1039.

*This reaction is best performed in specialized high-pressure-tolerant glassware.

*Preparative Synthesis of (S,S)-Cy^HA-D*

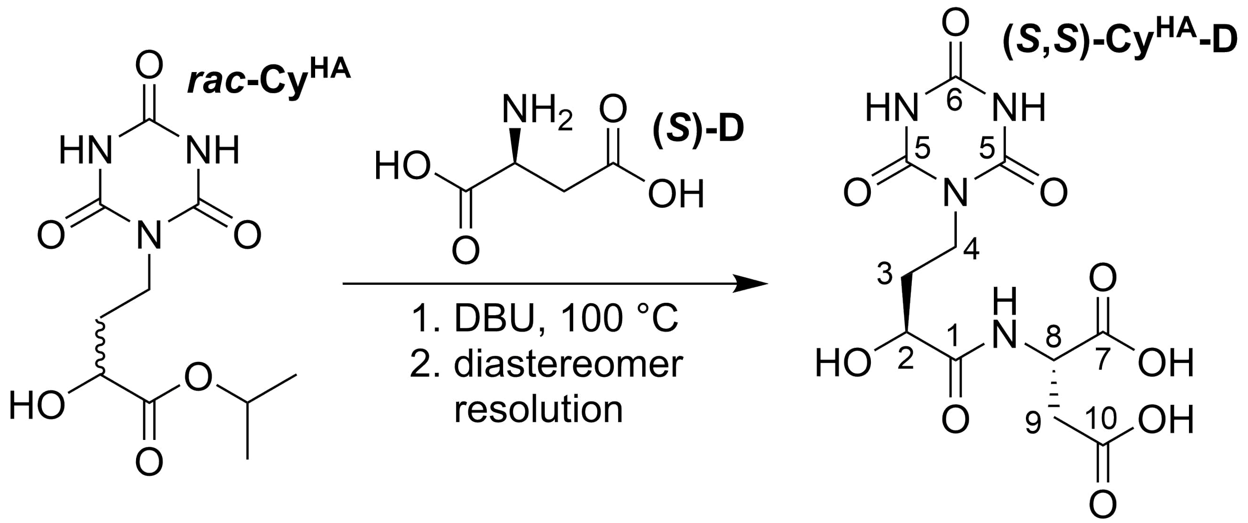

To a glass vial containing a recently prepared sample of *rac*-Cy$^{HA}$ isopropyl ester (137 mg, 0.5 mmol), vacuum-dried L-aspartic acid (i.e., (*S*)-D, 83.2 mg, 625 µmol) and dry DBU (0.3 mL, 2 mmol) were added. The glass vial, equipped with a stir bar, was sealed and stirred at 100 °C for 48 hours, forming a thick, brown solution. Upon returning to room temperature, the solution formed a thick gel. This gel was dissolved in 450 µL water with 50 µL formic acid, and the pH of the resulting solution was adjusted to 5 by the further addition of formic acid. The solution was purified in three chromatographic stages. In the first stage, the solution was loaded onto a column containing 75 mL of QAE Sephadex A-25 resin suspended in ammonium formate buffer, 50 mM, pH 5, and separated using a Teledyne Isco CombiFlash Rf+ purification system (monitoring by UV absorbance at 210 nm) with a gradient of ammonium formate pH 5, 50 mM, to ammonium formate, pH 5, 500 mM. The fractions containing product were pooled and reduced to about 5 mL with a rotary evaporator. In the second stage of purification, this solution was eluted over 10 g of Dowex 50WX8 cation exchange resin, hydrogen form, with 0.1 % aqueous formic acid solution. 100 mL were collected and the solvent was reduced to about 5 mL with a rotary evaporator. In the third stage of purification, this solution was separated using a Teledyne Isco CombiFlash Rf+ purification system (monitoring by UV absorbance at 210 nm) with a RediSep C18Aq 150 g Gold column with a 100% water eluent. This method of purification gave a slight degree of separation between the two diastereomers of Cy$^{HA}$-D. The fractions containing product (as identified by LC/MS) were then lyophilized and redissolved in $D_2O$ for $^1$H NMR analysis. From this analysis, the fractions could be separated into three groups: the first containing pure diastereomer 1 (i.e., the first diastereomer to elute), the second containing a mixture of the two diastereomers, and the third containing pure diastereomer 2 (i.e., the second diastereomer to elute). The NMR samples were pooled according to group and lyophilized. The second group (containing a mixture of the two diastereomers) was redissolved in 1 mL water and again separated using a Teledyne Isco CombiFlash Rf+ purification system with a

RediSep C18Aq 150 g Gold column with a 100% water eluent. This process was continued until the diastereomers were fully separated. 29.2 mg of diastereomer 1 (16.9%), which was found to be (*R*,*S*)-Cy$^{HA}$-D, were obtained, and 25.6 mg of diastereomer 2 (14.8%), which was found to be (*S*,*S*)-Cy$^{HA}$-D, were obtained. (See below for stereochemical determination.)

(*S*,*S*)-Cy$^{HA}$-D: $^1$H NMR (800 MHz, $D_2O$, 5 °C): δ 4.58 (dd, 6.7 Hz, 5.1 Hz, 1H, **H$^8$**), 4.05 (dd, 8.4 Hz, 3.6 Hz, 1H, **H$^2$**), 3.75 (m, 1H, **H$^4$**), 3.68 (m, 1H, **H$^{4'}$**), 2.77 (m, 2H, **H$^9$**), 1.89 (m, 1H, **H$^3$**), 1.69 (m, 1H, **H$^{3'}$**). $^{13}$C NMR (201 MHz, $D_2O$, 5 °C): 175.6 (**C1**, **C7**, or **C10**), 174.3 (**C1**, **C7**, or **C10**), 173.8 (**C1**, **C7**, or **C10**), 150.7 (**C5**), 149.9 (**C6**), 68.9 (**C2**), 48.5 (**C8**), 37.5 (**C4**), 35.2 (**C9**), 31.0 (**C3**). HRMS (m/z): $[M+H]^+$ $C_{11}H_{15}O_9N_4$, theoretical mass: 347.0834, actual mass: 347.0840.

(*R*,*S*)-Cy$^{HA}$-D: $^1$H NMR (800 MHz, $D_2O$, 5 °C): δ 4.57 (dd, 6.8 Hz, 5.1 Hz, 1H, **H$^8$**), 4.03 (dd, 8.4 Hz, 3.6 Hz, 1H, **H$^2$**), 3.75 (m, 1H, **H$^4$**), 3.70 (m, 1H, **H$^{4'}$**), 2.77 (m, 2H, **H$^9$**), 1.88 (m, 1H, **H$^3$**), 1.72 (m, 1H, **H$^{3'}$**). $^{13}$C NMR (201 MHz, $D_2O$, 5 °C): 175.6 (**C1**, **C7**, or **C10**), 174.4 (**C1**, **C7**, or **C10**), 173.7 (**C1**, **C7**, or **C10**), 150.7 (**C5**), 149.9 (**C6**), 68.9 (**C2**), 48.6 (**C8**), 37.5 (**C4**), 35.2 (**C9**), 31.1 (**C3**). HRMS (m/z): $[M+H]^+$ $C_{11}H_{15}O_9N_4$, theoretical mass: 347.0834, actual mass: 347.0833.

*Stereochemical Determination of Cy$^{HA}$-D*

*1. Synthesis of Morpholine-2,5-Dione of Cy$^{HA}$-D (Cyclic Cy$^{HA}$-D)*

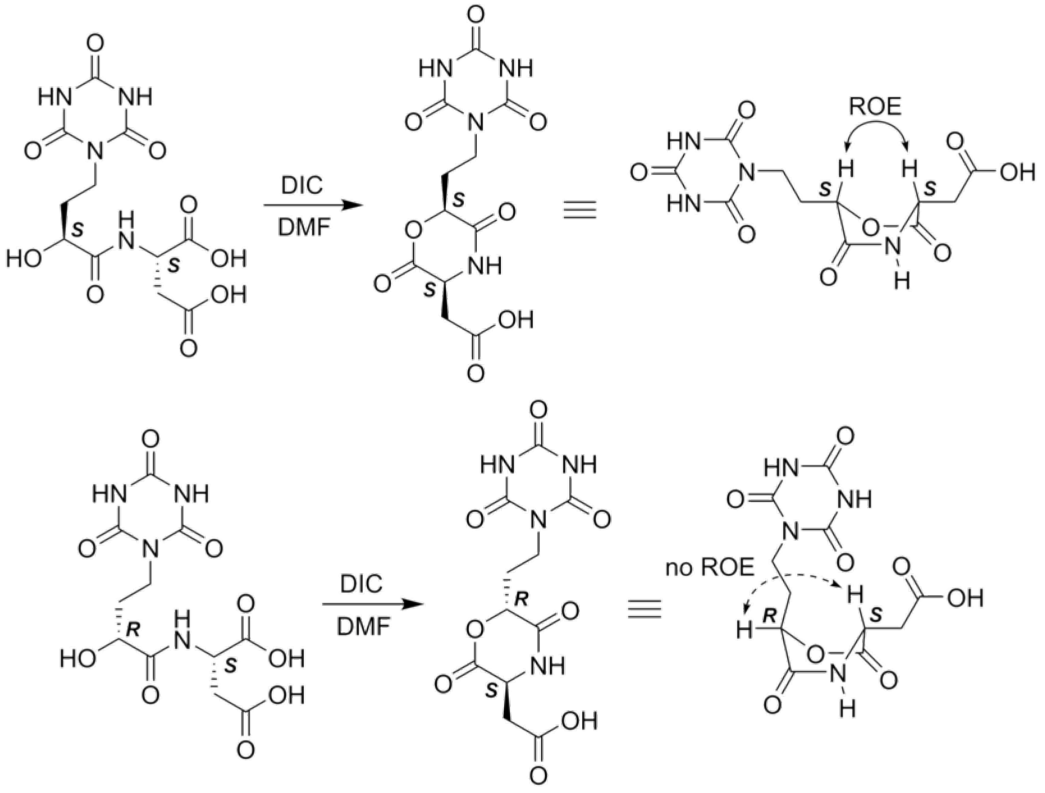

In a glass vial, 6.9 mg of a pure diastereomer of Cy$^{HA}$-D (20 μmol) was dissolved in 200 μL DMF. With stirring at room temperature, 3.4 μL of *N*,*N′*-diisopropylcarbodiimide (22 μmol) were added, and the solution was stirred for a further 3 hours. The solvent was removed by placing the open vial under vacuum for three days. The resulting residue was washed four times with 400 μL DCM, each time vortexing the crude mixture for one minute followed by centrifugation and decantation. The resulting crude solid was dried under vacuum and dissolved in 600 μL *d6*-DMSO for NMR analysis.

*2. NMR Analysis of Cyclic Cy$^{HA}$-D*

NMR analysis of the sample was performed on a Bruker 800 MHz instrument. 1D ROE analysis of the α-protons of cyclic Cy$^{HA}$-D, which adopts a boat-like structure, determined the relative orientation of the two stereocenters (see above). A through-space correlation will be observed between the α-protons of cyclic (*S*,*S*)-Cy$^{HA}$-D, but no such correlation will be present for cyclic (*R*,*S*)-Cy$^{HA}$-D. It was found that diastereomer 2 was the desired (*S*,*S*)-diastereomer.

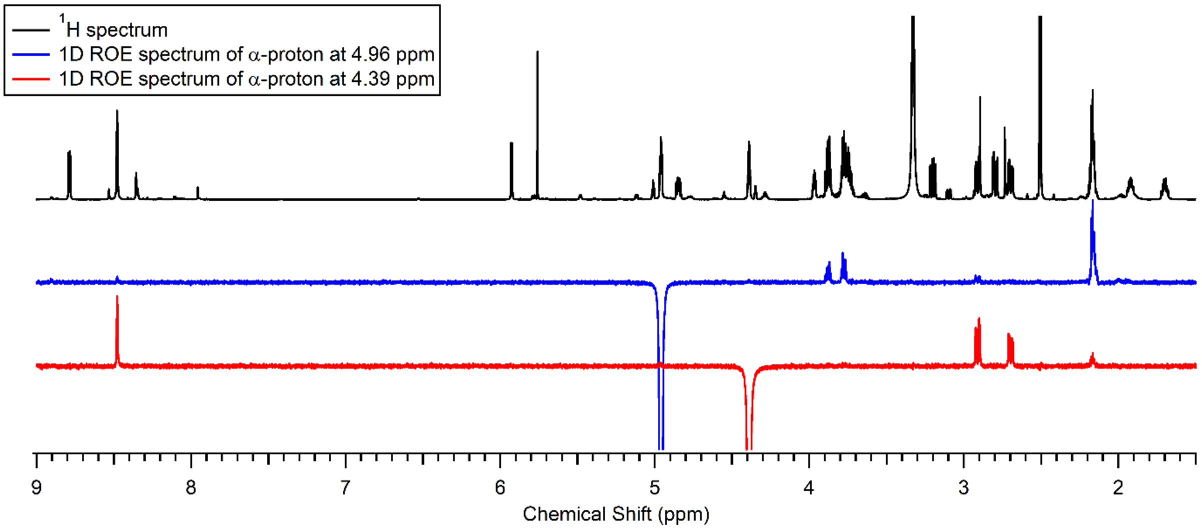

**Supplementary Figure 1.** NMR analysis of cyclic Cy$^{HA}$-D diastereomer 1 ((*R*,*S*)-diastereomer). Note that no through-space correlation exists between the two α-protons, indicating that the two α-carbons are of opposite stereochemical configuration.

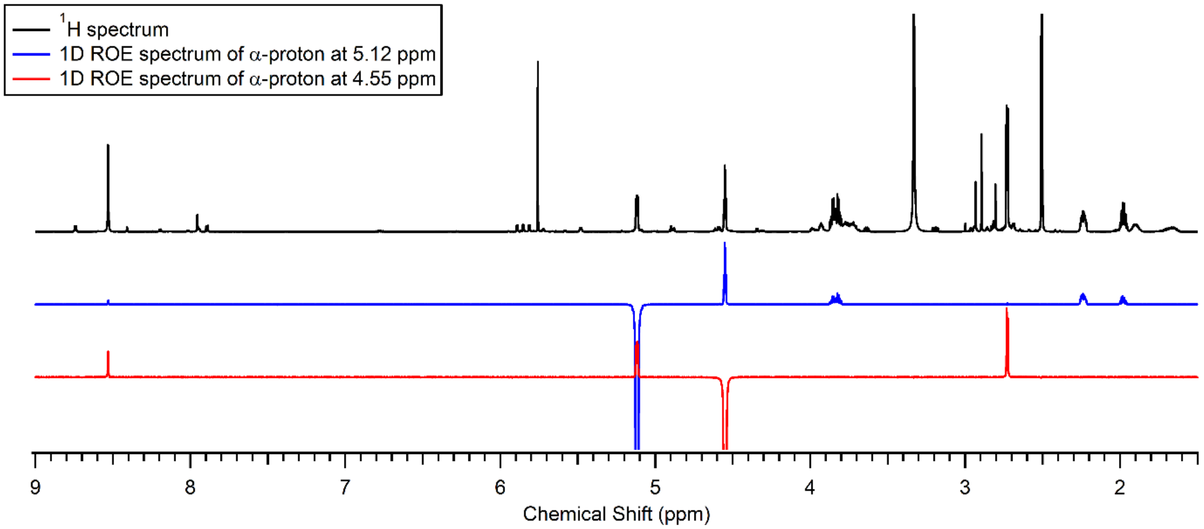

**Supplementary Figure 2.** NMR analysis of cyclic Cy$^{HA}$-D diastereomer 2 ((*S*,*S*)-diastereomer). Note that a through-space correlation exists between the two α-protons, indicating that the two α-carbons are of the same stereochemical configuration.

*Preparative/Prebiotic Synthesis of $(Cy^{HA}\text{-}D)_2$*

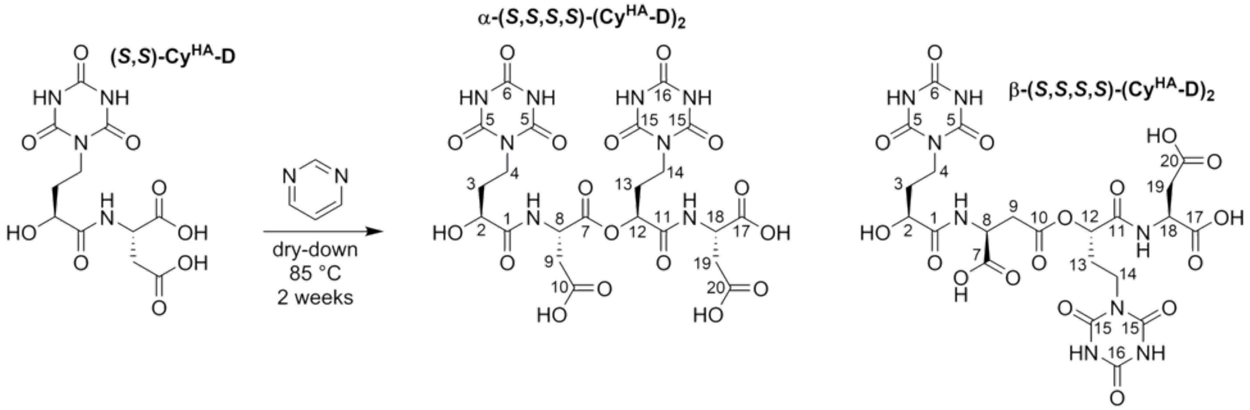

51.9 mg (*S*,*S*)-$Cy^{HA}$-D (150 µmol) were dissolved in 1.2 mL water. 47 µL of pyrimidine (48 mg, 600 µmol) were added to the solution. The solution of $Cy^{HA}$-D and pyrimidine was applied evenly across the full surface of six Fisherfinest Premium Superslip cover glasses (each 2.4 cm x 4 cm, 200 µL per cover glass) that were previously desilanized by heating in a bath of 5 M NaOH at approximately 90 °C for two hours and washed with deionized water. These cover glasses were placed uncovered in an oven at 85 °C for two weeks. After this period, the material deposited on the cover glasses was recovered by redissolving the material on each slide with 200 µL water, followed by two further washings with 100 µL water. The material from all six slides was pooled. The components of the solution were then separated by preparative HPLC (monitoring by UV absorbance at 210 nm) using an Agilent Prep-C18 column 50 mm x 21.2 mm with a flow rate of 25 mL/min. The solution was injected onto the column in 300 µL portions and eluted with a gradient of 98% aqueous/2% acetonitrile for 5 minutes, followed by a ramp to 85% aqueous/15% acetonitrile over 12 minutes. The aqueous mobile phase contained 0.1% TFA. Four closely eluting peaks with the mass of $(Cy^{HA}\text{-}D)_2$ (two major peaks and two minor peaks) were collected. The two minor peaks were assumed to be epimerization products and were not used in this study. The two major peaks were lyophilized to yield 6.9 mg (13.6%) of α-(*S*,*S*,*S*,*S*)-$(Cy^{HA}\text{-}D)_2$ and 3.5 mg (6.9%) of β-(*S*,*S*,*S*,*S*)-$(Cy^{HA}\text{-}D)_2$ isomer. (See below for structural assignment.)

α-(*S*,*S*,*S*,*S*)-$(Cy^{HA}\text{-}D)_2$: $^1$H NMR (800 MHz, $D_2O$): δ 5.16 (dd, 7.0 Hz, 5.2 Hz, 1H, $\mathbf{H^{12}}$), 4.99 (t, 6.1 Hz, 1H, $\mathbf{H^{8}}$), 4.77 (dd, 7.0 Hz, 5.2 Hz, 1H, $\mathbf{H^{18}}$), 4.30 (dd, 8.6 Hz, 3.5 Hz, 1H, $\mathbf{H^{2}}$), 4.01-3.87 (m, 4H, $\mathbf{H^{4}}$, $\mathbf{H^{4'}}$, $\mathbf{H^{14}}$, $\mathbf{H^{14'}}$), 3.09 (dd, 17.2 Hz, 5.6 Hz, 1H, $\mathbf{H^{9}}$), 3.04 (dd, 17.2 Hz, 6.8 Hz, 1H, $\mathbf{H^{9'}}$), 3.00 (dd, 17.1 Hz, 5.2 Hz, 1H, $\mathbf{H^{19}}$), 2.96 (dd, 17.1 Hz, 7.0 Hz, 1H, $\mathbf{H^{19'}}$), 2.21 (m, 2H, $\mathbf{H^{13}}$, $\mathbf{H^{13'}}$), 2.13 (m, 1H, $\mathbf{H^{3}}$), 1.94 (m, 1H, $\mathbf{H^{3'}}$). $^{13}$C NMR (201 MHz, $D_2O$): 175.8 (**C1**), 174.2 (**C20**), 174.0 (**C10**), 173.5 (**C17**), 170.8 (**C11**), 170.7 (**C7**), 150.9 (**C5**), 150.8 (**C15**), 150.0 (**C6** or **C16**), 149.9 (**C6** or **C16**), 72.9 (**C12**), 69.3 (**C2**), 49.1 (**C18**), 48.7 (**C8**), 37.9 (**C4**), 37.3 (**C14**), 35.3 (**C19**), 35.1 (**C9**), 31.5 (**C3**), 28.9 (**C13**). HRMS (m/z): $[M+Na]^+$ $C_{22}H_{26}O_{17}N_8Na$, theoretical mass: 697.1308, actual mass: 697.1319.

β-($S$,$S$,$S$,$S$)-($Cy^{HA}$-D)$_2$: $^1$H NMR (800 MHz, $D_2O$): δ 5.09 (dd, 8.4 Hz, 3.9 Hz, 1H, $\mathbf{H^{12}}$), 4.87 (t, 6.0 Hz, 1H, $\mathbf{H^{8}}$), 4.76 (dd, 6.9 Hz, 5.4 Hz, 1H, $\mathbf{H^{18}}$), 4.29 (dd, 8.4 Hz, 3.6 Hz, 1H, $\mathbf{H^{2}}$), 4.03-3.85 (m, 4H, $\mathbf{H^{4}}$, $\mathbf{H^{4'}}$, $\mathbf{H^{14}}$, $\mathbf{H^{14'}}$), 3.14 (m, 2H, $\mathbf{H^{9}}$), 3.00 (dd, 17.2 Hz, 5.4 Hz, 2H, $\mathbf{H^{19}}$), 2.96 (dd, 17.2 Hz, 6.9 Hz, $\mathbf{H^{19'}}$), 2.18 (m, 2H, $\mathbf{H^{13}}$, $\mathbf{H^{13'}}$), 2.11 (m, 1H, $\mathbf{H^{3}}$), 1.94 (m, 1H, $\mathbf{H^{3'}}$). $^{13}$C NMR (201 MHz, $D_2O$): δ 175.7 (**C1**), 174.2 (**C20**), 173.6 (**C7**), 171.3 (**C17**), 171.2 (**C11**), 150.9 (**C5** or **C15**), 150.8 (**C5** or **C15**), 150.0 (**C6** or **C16**), 149.9 (**C6** or **C16**), 72.4 (**C12**), 69.4 (**C2**), 49.0 (**C18**), 48.6 (**C8**), 37.9 (**C4**), 37.4 (**C14**), 35.4 (**C9**), 35.3 (**C19**), 31.3 (**C3**), 29.0 (**C13**). HRMS (m/z): $[M+Na]^+$ $C_{22}H_{26}O_{17}N_8Na$, theoretical mass: 697.1308, actual mass: 697.1306.

*Structural Assignment of ($Cy^{HA}$-D)$_2$ Isomers*

Comparison of the $^1H$ NMR spectra of the two isolated isomers of ($Cy^{HA}$-D)$_2$ allowed assignment of the location of esterification. Upon dimerization of $Cy^{HA}$-D, the $^1H$ signals of the aliphatic protons adjacent to the ester carbonyl moiety shift downfield relative to their unesterified counterparts. Based on the chemical shifts of the α- and β-protons of the first aspartic acid residue of each dimer ($\alpha_1$ and $\beta_2$, respectively), it was found that the primary dimer (i.e., produced in higher yield) was α-($Cy^{HA}$-D)$_2$ and the secondary dimer (i.e., produced in lower yield) was β-($Cy^{HA}$-D)$_2$.

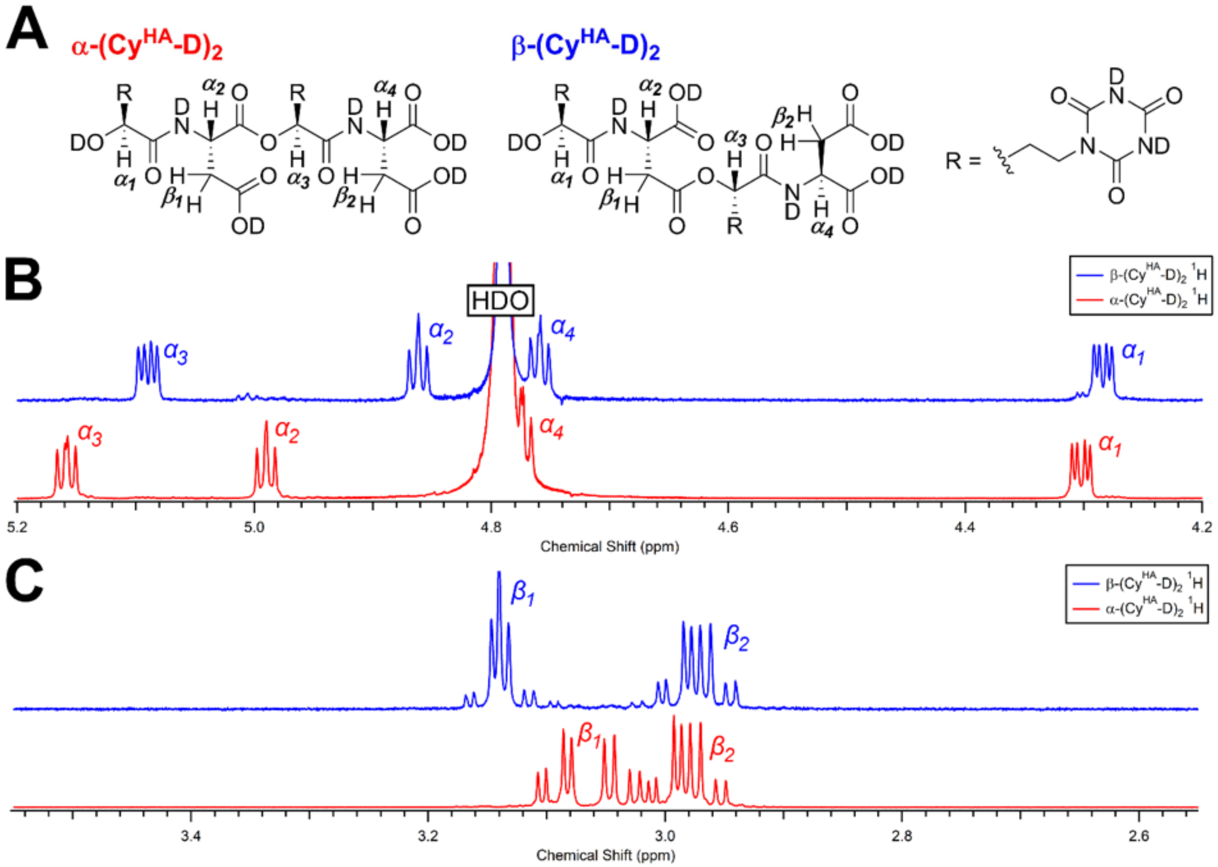

**Supplementary Figure 3.** Structural assignment of ($Cy^{HA}$-D)$_2$ isomers. **A.** Chemical structures of α-($Cy^{HA}$D)$_2$ and β-($Cy^{HA}$-D)$_2$ as they appear in $D_2O$, with relevant protons labeled. **B.** $^1H$ NMR spectra of α-($Cy^{HA}$-D)$_2$ (red) and β-($Cy^{HA}$-D)$_2$ (blue) in the α-proton region. The more downfield chemical shift of proton $\alpha_2$ in α-($Cy^{HA}$-D)$_2$ relative to $\alpha_2$ in β-($Cy^{HA}$-D)$_2$ indicates esterification at the α-carboxylic acid. **C.** $^1H$ NMR spectra of α-($Cy^{HA}$-D)$_2$ (red) and β-($Cy^{HA}$-D)$_2$ (blue) in the β-proton region (of aspartic acid residues). The more downfield chemical shift of proton $\beta_1$ in β-($Cy^{HA}$-D)$_2$ relative to $\beta_1$ in α-($Cy^{HA}$-D)$_2$ indicates esterification at the β-carboxylic acid.

*Preparative Synthesis of (S)-Mel^HA*

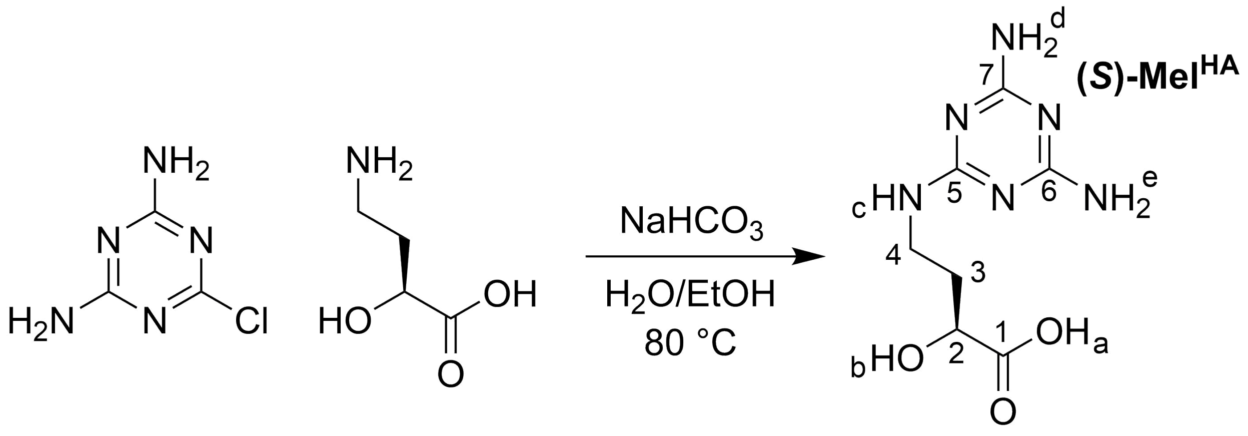

To 225 mL of 1:1 ethanol:water, 2-chloro-4,6-diamino-1,3,5-triazine (1.45 g, 10 mmol), (*S*)-4-amino-2-hydroxybutyric acid (1.79 g, 15 mmol), and sodium bicarbonate (1.68 g, 20 mmol) were added. The suspension was stirred at 80 °C for two days, after which time it appeared as a clear solution. The solution was then transferred to an ice bath. Once the internal temperature of the solution had reached 0 °C, concentrated HCl was added until the pH reacted 3.9. This produced a fine, white precipitate. The precipitate was filtered and washed with cold water and ethanol. The solid material was collected and dried under vacuum, giving compound (*S*)-Mel$^{HA}$ (1.8 g, 79%). $^{1}$H NMR (700 MHz, *d6*-DMSO): δ 6.48 (t, 5.6 Hz, 1H, **H$^{c}$**), 6.18 (s [br], 2H, **H$^{d}$** or **H$^{e}$**), 6.0 (s [br], 2H, **H$^{d}$** or **H$^{e}$**), 3.96 (dd, 8.8 Hz, 3.9 Hz, 1H, **H$^{2}$**), 3.28 (m, 2H, **H$^{4}$**), 1.87 (m, 1H, **H$^{3}$**), 1.63 (m, 1H, **H$^{3'}$**). $^{13}$C NMR (176 MHz, *d6*-DMSO): δ 176.3 (**C1**), 167.3 (br, **C6** or **C7**), 167.0 (br, **C6** or **C7**), 166.6 (**C5**), 68.4 (**C2**), 37.1 (**C4**), 34.6 (**C3**). HRMS (m/z): [M+H]$^{+}$ $C_7H_{13}O_3N_6$, theoretical mass: 229.1044, actual mass: 229.1045.

*Preparative Synthesis of (S)-Mel$^{HA}$ Isopropyl Ester*

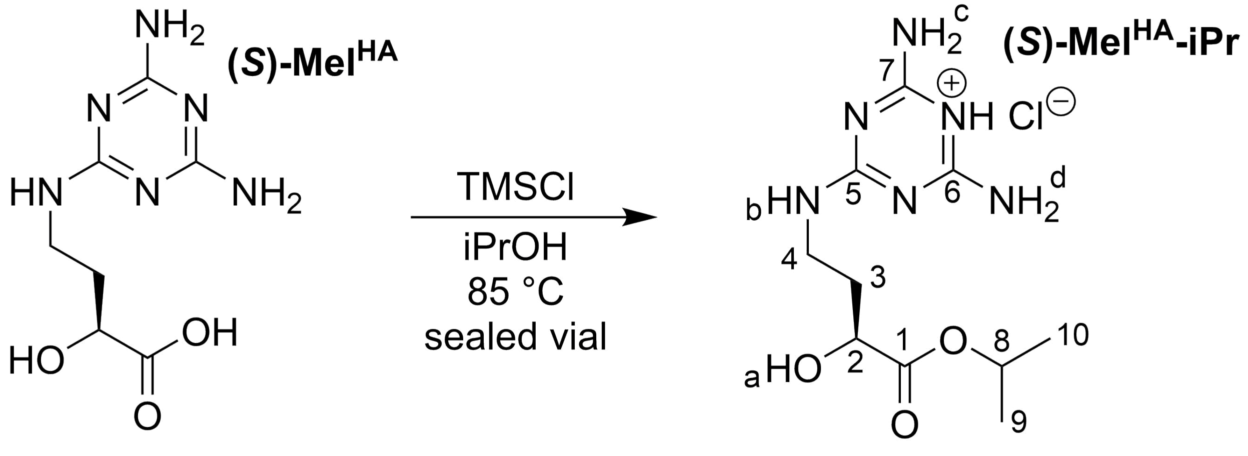

In a glass vial equipped with a stir bar, compound (*S*)-Mel$^{HA}$ (114 mg, 0.5 mmol) was suspended in 10 mL isopropanol, and trimethylsilyl chloride (635 µL, 5 mmol) was added. The vial was sealed and the mixture was stirred rapidly at 85 °C overnight, producing a solution. (WARNING: Heating solvents in sealed vessels above their boiling points creates a risk of explosion due to high pressure.)* The solvent was then evaporated first by blowing a stream of nitrogen onto the solution, and then by removing any residual solvent under vacuum, giving (*S*)-Mel$^{HA}$ isopropyl ester hydrochloride as a white solid. The yield, as determined by $^{1}$H NMR, was 84%. The solid was left in the glass vial and taken to the next synthetic step without any further purification. $^{1}$H NMR (800 MHz, *d6*-DMSO): δ 8.14 (s [br], 1H, $\mathbf{H^b}$), 8.04 (s [br], 2H, $\mathbf{H^c}$ or $\mathbf{H^d}$), 7.83 (s [br], 2H, $\mathbf{H^c}$ or $\mathbf{H^d}$), 4.91 (m, 6.3 Hz, 1H, $\mathbf{H^8}$), 4.05 (dd, 8.9 Hz, 3.9 Hz, 1H, $\mathbf{H^2}$), 3.38 (m, 2H, $\mathbf{H^4}$), 1.89 (m, 1H, $\mathbf{H^3}$), 1.72 (m, 1H, $\mathbf{H^{3'}}$), 1.20 (d, 6.3 Hz, 3H, $\mathbf{H^9}$ or $\mathbf{H^{10}}$), 1.19 (d, 6.3 Hz, 3H, $\mathbf{H^9}$ or $\mathbf{H^{10}}$). $^{13}$C NMR (201 MHz, *d6*-DMSO): δ 173.8 (**C1**), 158.8 (**C6** or **C7**), 158.6 (**C6** or **C7**), 157.0 (**C5**), 68.3 (**C2** or **C8**), 68.1 (**C2** or **C8**), 37.5 (**C4**), 33.6 (**C3**), 22.1 (**C9** or **C10**), 22.0 (**C9** or **C10**). HRMS (m/z): $[M+H]^+$ $C_{10}H_{19}O_3N_6$, theoretical mass: 271.1513, actual mass: 271.1517.

*This reaction is best performed in specialized high-pressure-tolerant glassware.

*Preparative Synthesis of (S,S)-Mel$^{HA}$-D*

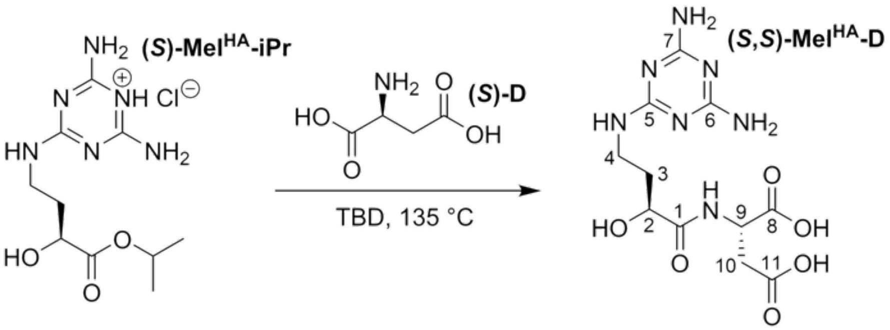

To a glass vial containing a recently prepared sample of (*S*)-Mel$^{HA}$ isopropyl ester hydrochloride (0.5 mmol), vacuum-dried L-aspartic acid (i.e., (*S*)-D, 80 mg, 0.6 mmol) and dry TBD (1,5,7-triazabicyclo[4.4.0]dec-5-ene, 237 mg, 1.7 mmol) were added. The glass vial, equipped with a stir bar, was sealed and stirred at 135 °C for 10 hours, producing a clear, brown, viscous solution. This mixture was dissolved in 450 µL water with 50 µL formic acid, and the pH of the resulting solution was adjusted to 5 by the further addition of formic acid. The solution was loaded onto a column containing 75 mL of QAE Sephadex A-25 resin suspended in ammonium formate buffer, 50 mM, pH 5, and separated using a Teledyne Isco CombiFlash Rf+ purification system (monitoring by UV absorbance at 235 nm) with a gradient of ammonium formate pH 5, 50 mM, to ammonium formate, pH 5, 500 mM. The fractions containing product were pooled and reduced to about 5 mL with a rotary evaporator. This solution was transferred to a conical tube and brought to a total volume of 12 mL by the addition of water. 12 mL of ethyl acetate were then added, and the solution was shaken vigorously for several minutes. After standing for about an hour, a bed of acetamide formed at the bottom of the aqueous layer. The organic phase was removed and the aqueous phase was heated to bring all of the acetamide into solution. This solution was separated using a Teledyne Isco CombiFlash Rf+ purification system (monitoring by UV absorbance at 235 nm) with a RediSep C18Aq 150 g Gold column with a 100% water eluent. The solution was injected onto the column in 4 mL portions. A small amount of another diastereomer (either (*R*,*S*)-Mel$^{HA}$-D, (*S*,*R*)-Mel$^{HA}$-D, or both, produced by epimerization during the synthesis) could be separated (eluting first) from the main product using this chromatographic method. Fractions containing the main product were pooled and reduced to about 5 mL with a rotary evaporator. This solution was then lyophilized to give the product, (*S*,*S*)-Mel$^{HA}$-D, free of any counter-ions or buffer salts. The reaction yield of Mel$^{HA}$-D was estimated by UV-LC/MS to be 56% based on UV peak integration of all melamine species at 235 nm. $^{1}$H NMR (800 MHz, $D_2O$): δ 4.51 (dd, 6.8 Hz, 5.4 Hz, 1H, **H$^{9}$**), 4.21 (dd, 7.4 Hz, 4.3 Hz, 1H, **H$^{2}$**), 3.44 (m, 2H, **H$^{4}$**), 2.81 (dd, 16.4 Hz, 5.1 Hz, 1H, **H$^{10}$**), 2.78 (dd, 16.4 Hz, 9.4 Hz, 1H, **H$^{10\prime}$**), 2.01 (m, 1H, **H$^{3}$**), 1.89 (m, 1H, **H$^{3\prime}$**). $^{13}$C NMR (201 MHz, $D_2O$): δ 176.3 (**C1**, **C8**, or **C11**), 175.8 (**C1**, **C8**, or **C11**), 175.5 (**C1**, **C8**, or **C11**), 160.7 (br, **C6** or **C7**), 159.5 (br, **C6** or **C7**), 157.3 (**C5**), 69.1 (**C2**), 50.6 (**C9**), 37.2 (**C4**), 36.6 (**C10**), 32.5 (**C3**). HRMS (m/z): $[M+H]^+$ $C_{11}H_{18}O_6N_7$, theoretical mass: 344.1313, actual mass: 344.1316.

*Preparative Synthesis of* rac-*Ad^HA*

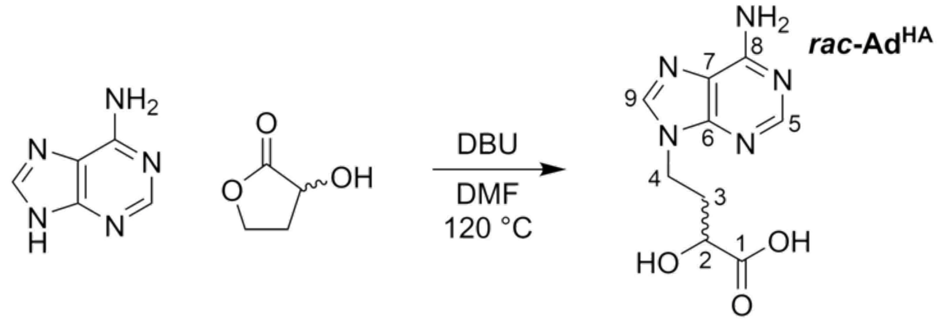

To 100 mL of DMF, 6.76 g of adenine (50 mmol) and 7.48 mL of DBU (50 mmol) were added. The mixture was heated to 120 °C with rapid stirring. Once the adenine had fully dissolved, 3.9 mL of *rac*-α-hydroxy-γ-butyrolactone (50 mmol) were added dropwise. The reaction was stirred at 120 °C for 24 hours. The solution was then poured onto a watch glass and allowed to evaporate over several days, eventually producing a thick, brown oil. To this oil, 50 mL of water were added, and the components were gently mixed, forming a suspension. The suspension was then centrifuged. The supernatant was decanted, and the pellet was resuspended in 10 mL water, vortexed, and centrifuged, and the supernatant decanted. This was repeated once more. The pooled supernatants were reduced in volume to about 20 mL. To this solution, 2.5 mL of formic acid were added to bring the pH to 4-4.5. 60 mL of ethanol were then added, producing a precipitate. The suspension placed at -20 °C overnight, and was then centrifuged and the supernatant discarded. The pellet was then washed with 20 mL hot water followed by centrifugation and removal of the supernatant. This water-washing process was performed four times in total. The residual solid was then dried by lyophilization to yield *rac*-$Ad^{HA}$, which was determined to be 90 mole % pure by $^1H$ NMR, with some adenine and 2,4-dihydroxybutyric acid remaining. This sample was used directly in the next step of the synthesis. Further purification of this sample was attempted using a Teledyne Isco CombiFlash Rf+ purification system (monitoring by UV absorbance at 260 nm) with a RediSep C18Aq 150 g Gold column with a 100% water eluent. A 5 mL sample of a hot, saturated solution of this solid was immediately injected and the fractions containing product were pooled and lyophilized. This sample was free of 2,4-dihydroxybutyric acid, but contained 14 mole % adenine. The increase in free adenine content from the previous sample was attributed to the low solubility of $Ad^{HA}$ in water and the co-elution of adenine with $Ad^{HA}$. The reaction yield of $Ad^{HA}$ was estimated by UV-LC/MS to be 19% based on UV peak integration of all adenine species at 260 nm. $^1H$ NMR (700 MHz, *d6*-DMSO): δ 12.51 (s [br], 1H, $CO_2$**H**), 8.11 (s, 1H, $\mathbf{H^9}$), 8.07 (s, 1H, $\mathbf{H^5}$), 7.16 (s, 2H, N$\mathbf{H_2}$), 5.46 (s [br], 1H, O**H**), 4.22 (t, 7.2 Hz, 2H, $\mathbf{H^4}$), 3.87 (dd, 9.0 Hz, 3.9 Hz, 1H, $\mathbf{H^2}$), 2.22 (m, 1H, $\mathbf{H^3}$), 1.97 (m, 1H, $\mathbf{H^{3\prime}}$). $^{13}C$ NMR (176 MHz, *d6*-DMSO): δ 175.6 (**C1**), 156.4 (**C8**), 152.8 (**C5**), 150.0 (**C6**), 141.4 (**C9**), 119.3 (**C7**), 67.7 (**C2**), 40.5 (**C4**), 34.2 (**C3**). HRMS (m/z): $[M+H]^+$ $C_9H_{12}N_5O_3$, theoretical mass: 238.0935, actual mass: 238.0933.

*Preparative Synthesis of* rac-*Ad^HA Isopropyl Ester*

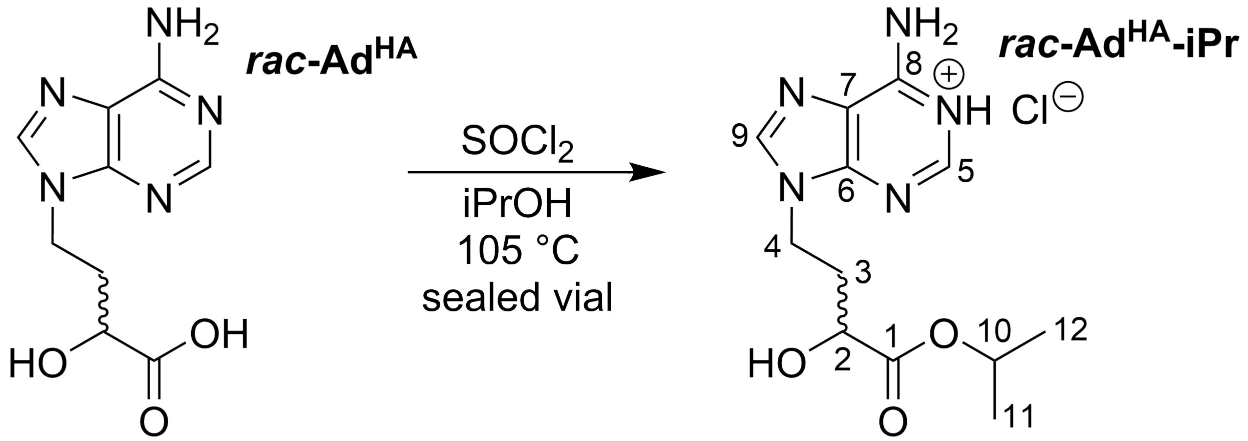

In a glass vial equipped with a stir bar, crude *rac*-Ad$^{HA}$ (250 mg, ~1 mmol) was suspended in 10 mL isopropanol with rapid stirring. The vial is then submersed in an ice bath and allowed to cool. 400 µL of thionyl chloride are then carefully added dropwise to the mixture with rapid stirring. (WARNING: The direct addition of thionyl chloride to protic solvents is violently exothermic.) After all of the thionyl chloride had been added, the vial is removed from the ice bath and allowed to reach room temperature.* The vial was then sealed and transferred to an oil bath at 100-110 °C. (WARNING: Heating solvents in sealed vessels above their boiling points creates a risk of explosion due to high pressure.)** The mixture was stirred overnight, producing a clear, yellow-brown solution. The solvent was then evaporated first by blowing a stream of nitrogen onto the solution, and then by removing any residual solvent under vacuum, giving *rac*-Ad$^{HA}$ isopropyl ester hydrochloride as an off-white solid. The yield, as determined by $^1$H NMR, was 74%. The solid was left in the glass vial and taken to the next synthetic step without any further purification. $^1$H NMR (800 MHz, *d6*-DMSO): δ 8.14 (s, 1H, **H$^9$**), 8.08 (s, 1H, **H$^5$**), 7.20 (s, 2H, N**H$_2$**), 5.66 (d, 5.7 Hz, 1H, O**H**), 4.84 (sept, 6.3 Hz, 1H, **H$^{10}$**), 4.24 (m, 2H, **H$^4$**), 3.97 (m, 1H, **H$^2$**), 2.20 (m, 1H, **H$^3$**), 2.04 (m, 1H, **H$^{3'}$**), 1.16 (d, 6.3, Hz 3H, **H$^{11}$** or **H$^{12}$**), 1.12 (d, 6.3 Hz, 3H, **H$^{11}$** or **H$^{12}$**). $^{13}$C NMR (201 MHz, d6-DMSO): δ 173.4 (**C1**), 156.4 (**C8**), 152.8 (**C5**), 150.0 (**C6**), 141.4 (**C9**), 119.3 (**C7**), 68.2 (**C10**) 67.9 (**C2**), 40.5 (**C4**), 34.1 (**C3**), 22.0 (**C11** or **C12**), 21.8 (**C11** or **C12**). HRMS (m/z): [M+H]$^+$ $C_{12}H_{18}N_5O_3$, theoretical mass: 280.1404, actual mass: 280.1404.

*In the original synthesis of this compound, the thionyl chloride was added to the stirred mixture at room temperature. This modification of the synthesis is recommended as an additional safety measure.

**This reaction is best performed in specialized high-pressure-tolerant glassware.

*Preparative Synthesis of (S,S)-Ad^HA^-D*

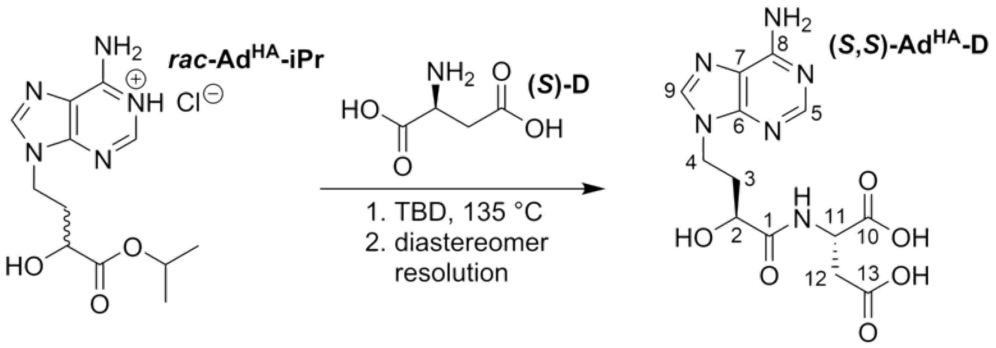

To a glass vial containing a recently prepared sample of *rac*-Ad$^{HA}$ isopropyl ester hydrochloride (1 mmol), vacuum-dried L-aspartic acid (i.e., (*S*)-D, 166.4 mg, 1.25 mmol) and dry TBD (1,5,7-triazabicyclo[4.4.0]dec-5-ene, 487.2 mg, 3.5 mmol) were added. The glass vial, equipped with a stir bar, was sealed and stirred at 135 °C for 4 hours, producing a clear, brown, viscous solution. This mixture was dissolved in 900 μL water with 100 μL formic acid, and the pH of the resulting solution was adjusted to 5 by the further addition of formic acid. The solution was loaded onto a column containing 75 mL of QAE Sephadex A-25 resin suspended in ammonium formate buffer, 50 mM, pH 5, and separated using a Teledyne Isco CombiFlash Rf+ purification system (monitoring by UV absorbance at 260 nm) with a gradient of ammonium formate pH 5, 50 mM, to ammonium formate, pH 5, 500 mM. The fractions containing product were pooled and reduced to about 10 mL with a rotary evaporator. This solution was then transferred to a separatory funnel. 50 mL of ethyl acetate were then added, and the solution was shaken vigorously for several minutes. After standing for about an hour, a bed of acetamide formed at the bottom of the aqueous layer. The aqueous and organic phases were separated, and the remaining acetamide was dissolved in 10 mL water and added to the aqueous phase. This solution was separated using a Teledyne Isco CombiFlash Rf+ purification system (monitoring by UV absorbance at 260 nm) with a RediSep C18Aq 150 g Gold column with a 100% water eluent. The solution was injected onto the column in 5 mL portions. Two well-separated peaks eluted which, when analyzed by LC/MS, both showed the mass of Ad$^{HA}$-D. Anticipating that the (*S*,*S*)-diastereomer of Ad$^{HA}$-D, like (*S*,*S*)-Mel$^{HA}$-D and (*S*,*S*)-Cy$^{HA}$-D, would elute second (see above), fractions corresponding to the second peak were pooled and reduced to about 5 mL with a rotary evaporator. This solution was then lyophilized to give the product, (*S*,*S*)-Ad$^{HA}$-D, free of any counter-ions or buffer salts. (See below for stereochemical determination.) The reaction yield of Ad$^{HA}$-D (both diastereomers) was estimated by UV-LC/MS to be 48% based on UV peak integration of all adenine species at 260 nm (except parent adenine). $^{1}$H NMR (800 MHz, $D_2O$): δ 8.28 (s, 1H, **H$^{9}$**), 8.21 (s, 1H, **H$^{5}$**), 4.40 – 4.13 (m, 3H, **H$^{4}$**, **H$^{11}$**), 4.10 (dd, 7.8 Hz, 4.3 Hz, 1H, **H$^{2}$**), 2.71 (m, 2H, **H$^{12}$**), 2.30 (m, 1H, **H$^{3}$**), 2.17 (m, 1H, **H$^{3'}$**). $^{13}$C NMR (201 MHz, $D_2O$): δ 175.8 (**C1**, **C10**, or **C13**), 175.3 (**C1**, **C10**, or **C13**), 175.0 (**C1**, **C10**, or **C13**), 150.9 (**C8**), 148.7 (**C5**), 145.9 (**C6**), 144.5 (**C9**), 118.1 (**C7**), 68.5 (**C2**), 50.2 (**C11**), 40.5 (**C4**), 36.7 (**C12**), 32.7 (**C3**). HRMS (m/z): $[M+H]^+$ $C_{13}H_{17}N_6O_6$, theoretical mass: 353.1204, actual mass: 353.1199.

*Stereochemical Determination of Ad$^{HA}$-D*

*1. Synthesis of Morpholine-2,5-Dione of Ad$^{HA}$-D (Cyclic Ad$^{HA}$-D)*

A crude NMR sample of cyclic Ad$^{HA}$-D, chromatographic diastereomer 2, was prepared exactly as was described for cyclic Cy$^{HA}$-D (see above).

*2. NMR Analysis of Cyclic Cy$^{HA}$-D*

NMR analysis of the sample was performed on a Bruker 800 MHz instrument. 1D ROE analysis of the α-protons of cyclic Ad$^{HA}$-D, which adopts a boat-like structure, determined the relative orientation of the two stereocenters (as with cyclic Cy$^{HA}$-D; see above). A through-space correlation will be observed between the α-protons of cyclic (*S*,*S*)-Ad$^{HA}$-D, but no such correlation will be present for cyclic (*R*,*S*)-Ad$^{HA}$-D. It was found that the isolated diastereomer was the desired (*S*,*S*)-diastereomer.

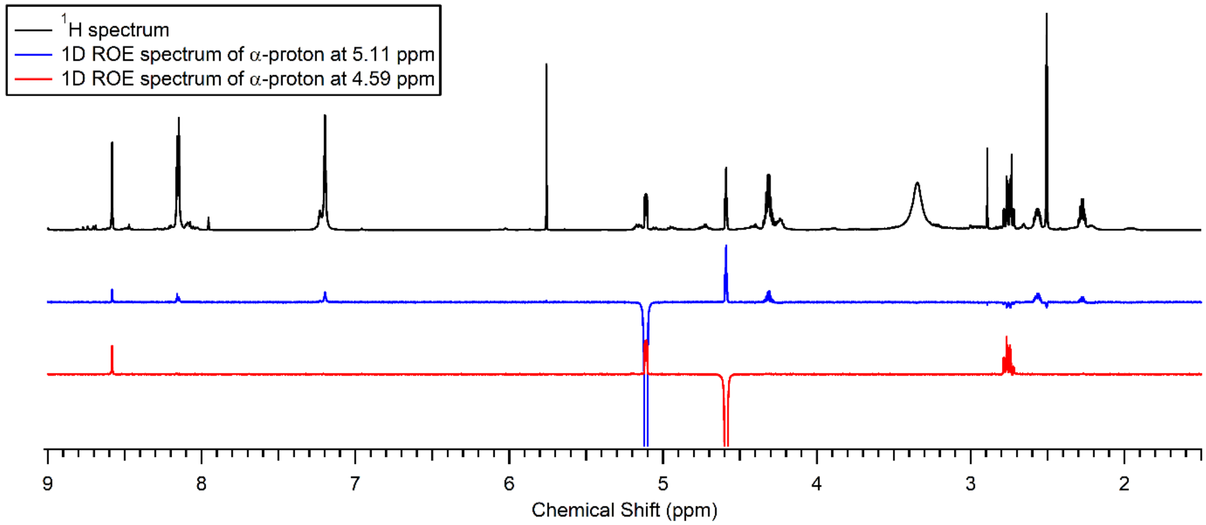

**Supplementary Figure 4.** NMR analysis of cyclic Ad$^{HA}$-D diastereomer 2 ((*S*,*S*)-diastereomer). Note that a through-space correlation exists between the two α-protons, indicating that the two α-carbons are of the same stereochemical configuration.

### III. Prebiotic Syntheses of Model Proto-Nucleic Acid Monomers[1]

*Prebiotic Synthesis of Cy$^{HA}$*

*1. Synthetic Protocol*

To an 8 dram vial equipped with a stir bar, sodium formate buffer (1 M, pH 4, 20 mL), cyanuric acid (129 mg, 1 mmol), and potassium ferrocyanide trihydrate (422.4 mg, 1 mmol) were added. The suspension was stirred rapidly with mild heating until all of the components dissolved. Acrolein (67 µL, 1 mmol) was then added, and the vial was sealed and stirred in an oil bath at 45 °C for 22 hours, over which time the reaction took on a green color. Samples were taken for UV-LC/MS analysis at 0, 2, 4, 6, 20, and 22 hours. The reaction was then stopped and extracted with 5 x 100 mL $CH_2Cl_2$. The combined extracts were dried over magnesium sulfate and the solvent was removed under vacuum. The resulting yellow residue was dissolved in 5 mL of hydrochloric acid (6 M) and refluxed for 18 hours.[2] The reaction solution was then lyophilized and redissolved in 600 µL $D_2O$ for NMR analysis.

*2. UV-LC/MS Analysis*

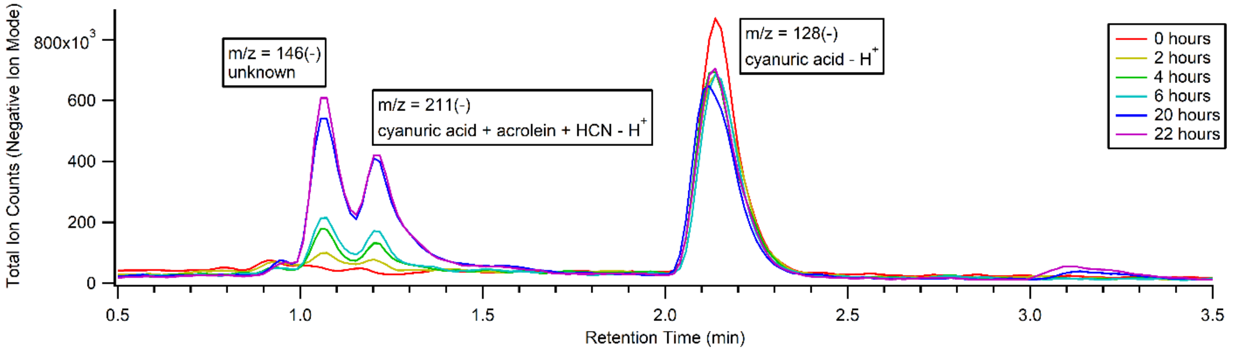

**Supplementary Figure 5.** LC/MS monitoring of the reaction of cyanuric acid with acrolein and potassium ferrocyanide. Due to the poor absorbance of cyanuric acid species, the total ion traces (negative mode) are shown instead. The mass of the Cy$^{HA}$ cyanohydrin (211(-)) is detected.

*3. NMR Analysis*

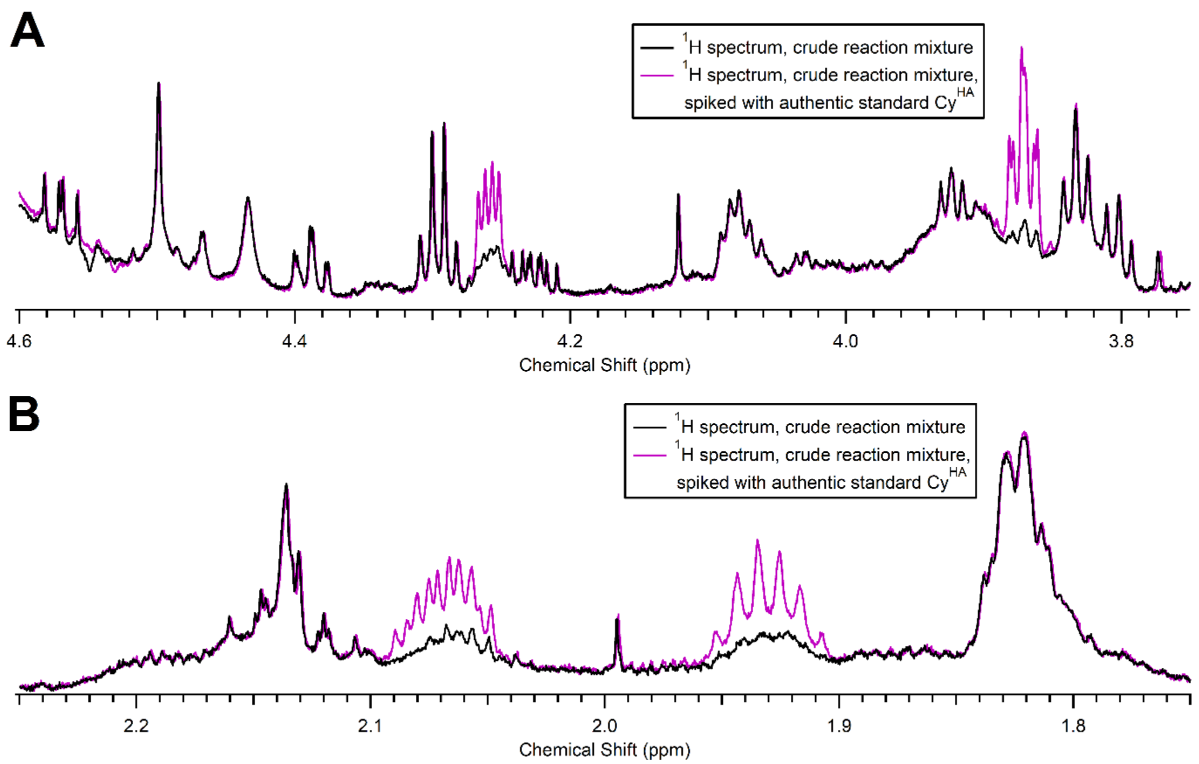

**Supplementary Figure 6.** $^1$H NMR analysis of the hydrolyzed extract of the reaction of cyanuric acid with acrolein and potassium ferrocyanide. The successful formation of Cy$^{HA}$ is indicated by the increase in signal intensity, without the formation of new signals, of the aliphatic protons of Cy$^{HA}$ upon spiking with an authentic standard. **A.** $^1$H NMR spectra from 3.75 to 4.6 ppm; the region where the α- and γ-protons of Cy$^{HA}$ appear. **B.** $^1$H NMR spectra from 1.75 to 2.25 ppm; the region where the β-protons of Cy$^{HA}$ appear.

*Prebiotic Synthesis of Ad$^{HA}$*

*1. Synthetic Protocol*

To a 2 dram vial equipped with a stir bar, sodium formate buffer (1 M, pH 4, 5 mL), adenine (67.6 mg, 0.5 mmol), and potassium ferrocyanide trihydrate (211.2 mg, 0.5 mmol) were added. The suspension was stirred rapidly with mild heating until all of the components dissolved. Acrolein (67 µL, 1 mmol) was then added, and the vial was sealed and stirred in an oil bath at 45 °C for 22 hours, over which time the reaction took on a green color. Samples were taken for UV-LC/MS analysis at 0, 2, 4, 6, 20, and 22 hours. The reaction was then stopped and extracted with 5 x 50 mL $CH_2Cl_2$. The combined extracts were dried over magnesium sulfate and the solvent was removed under vacuum. The resulting yellow residue was dissolved in 5 mL of hydrochloric acid (6 M) and refluxed for 18 hours.[2] The reaction solution was then lyophilized and redissolved in 600 µL $D_2O$ for NMR analysis.

### 2. UV-LC/MS Analysis

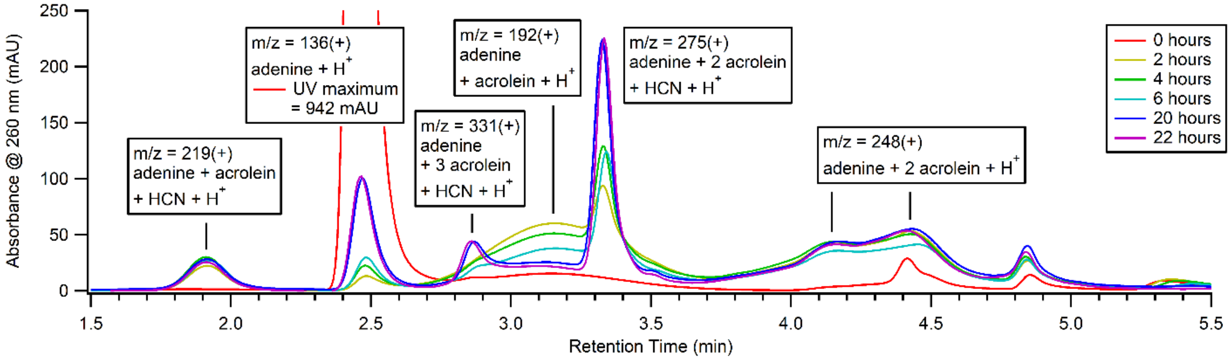

**Supplementary Figure 7.** UV-LC/MS monitoring of the reaction of adenine with acrolein and potassium ferrocyanide. The mass of the $Ad^{HA}$ cyanohydrin (219(+)) is detected.

### 3. NMR Analysis

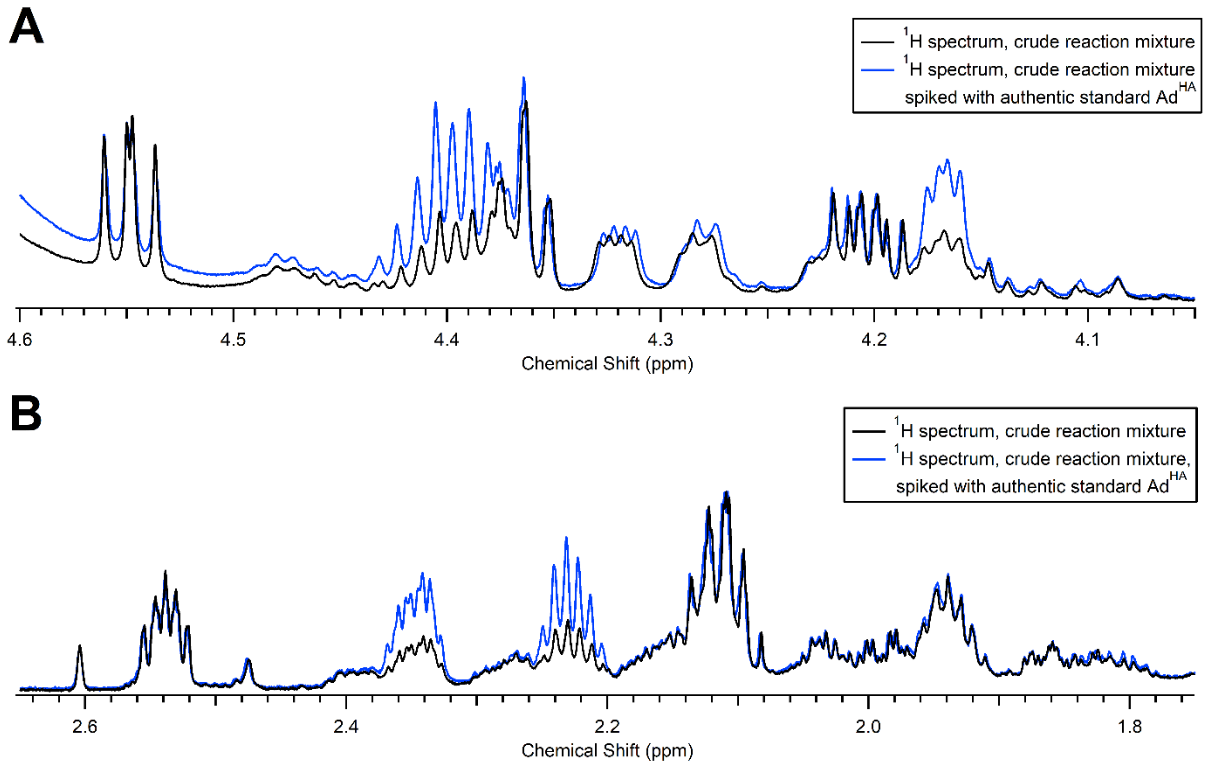

**Supplementary Figure 8.** $^{1}H$ NMR analysis of the hydrolyzed extract of the reaction of adenine with acrolein and potassium ferrocyanide. The successful formation of $Ad^{HA}$ is indicated by the increase in signal intensity, without the formation of new signals, of the aliphatic protons of $Ad^{HA}$ upon spiking with an authentic standard. **A.** $^{1}H$ NMR spectra from 4.05 to 4.6 ppm; the region where the α- and γ-protons of $Ad^{HA}$ appear. **B.** $^{1}H$ NMR spectra from 1.75 to 2.65 ppm; the region where the β-protons of $Ad^{HA}$ appear.

*Attempted Prebiotic Synthesis of Mel$^{HA}$ with Acrolein and Potassium Ferrocyanide*

*1. Synthetic Protocol*

To a 2 dram vial equipped with a stir bar, sodium formate buffer (1 M, pH 4, 5 mL) and melamine (63 mg, 0.5 mmol) were added. The suspension was stirred rapidly with mild heating until all of the melamine dissolved. Potassium ferrocyanide trihydrate (211.2 mg, 0.5 mmol) was then added, causing a cloudy, white precipitate to immediately form. This precipitate could not be redissolved with stirring at heating up to 90 °C. To the suspension, acrolein (67 µL, 1 mmol) was then added, and the vial was sealed and stirred in an oil bath at 45 °C for 22 hours, over which time the reaction took on a green color. The cloudy precipitate persisted for the entire reaction. Samples were taken for UV-LC/MS analysis at 0, 2, 4, 6, 20, and 22 hours. The reaction was then stopped and extracted with 5 x 50 mL $CH_2Cl_2$. The combined extracts were dried over magnesium sulfate and the solvent was removed under vacuum. The resulting yellow residue was dissolved in 5 mL of hydrochloric acid (6 M) and refluxed for 18 hours.[2] The reaction solution was then lyophilized and redissolved in 600 µL $D_2O$ for NMR analysis.

*2. UV-LC/MS Analysis*

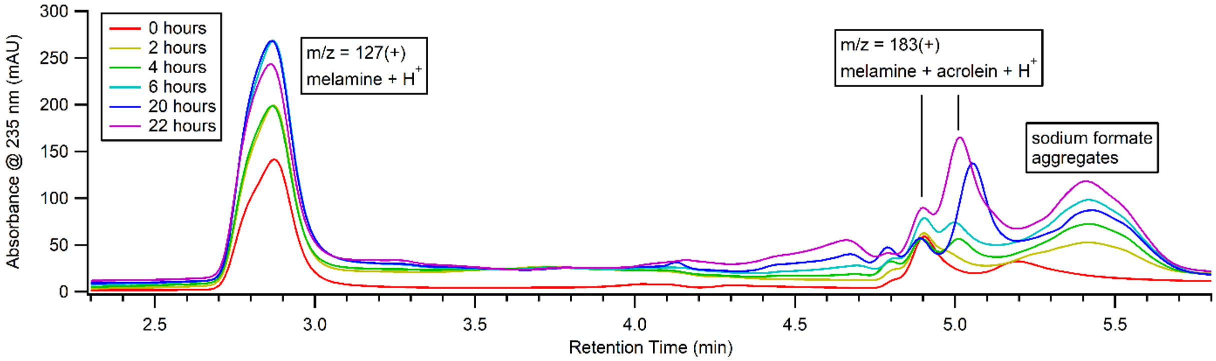

**Supplementary Figure 9.** UV-LC/MS monitoring of the reaction of melamine with acrolein and potassium ferrocyanide. The mass of the Mel$^{HA}$ cyanohydrin (210(+)) is not detected. Melamine initially co-precipitates with ferrocyanide. The recovery of melamine over time may be due to the conversion of ferrocyanide into different iron complexes that do not form insoluble salts with melamine.

*3. NMR Analysis*

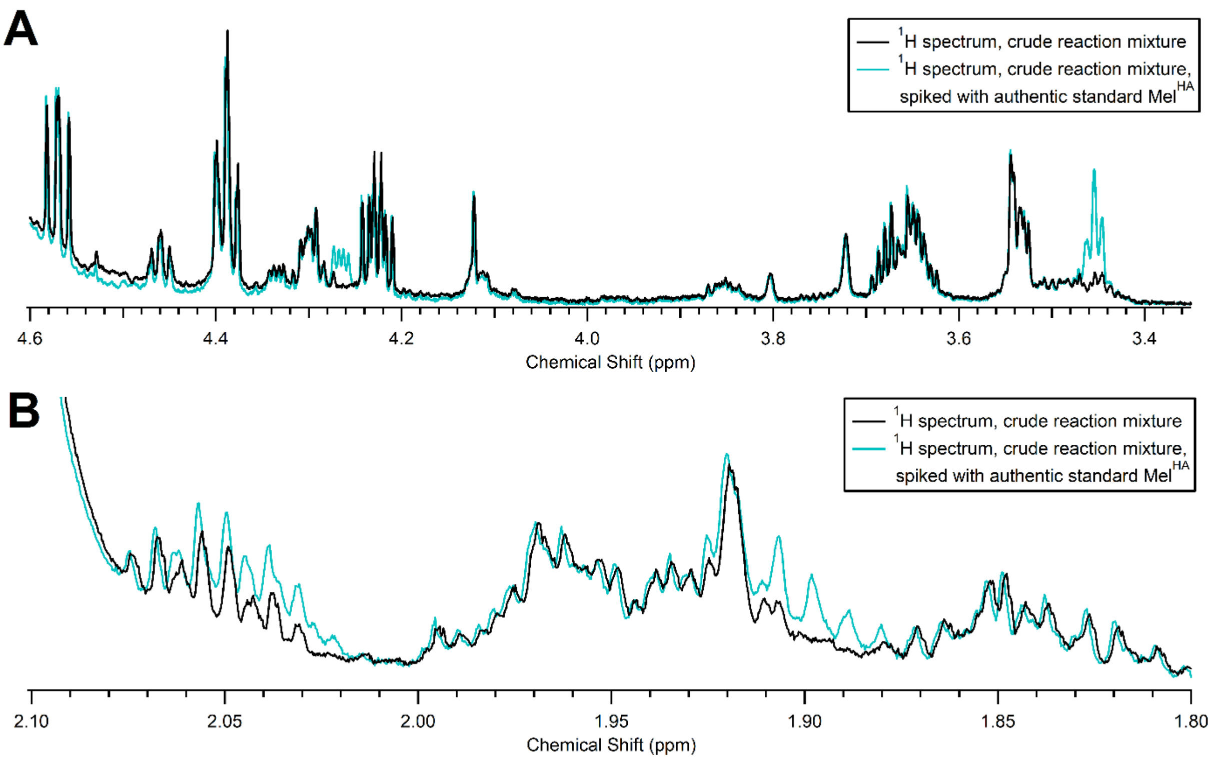

**Supplementary Figure 10.** $^1$H NMR analysis of the hydrolyzed extract of the reaction of melamine with acrolein and potassium ferrocyanide. The unsuccessful formation of Mel$^{HA}$ is indicated by the formation of new signals upon spiking with an authentic standard. **A.** $^1$H NMR spectra from 3.35 to 4.6 ppm; the region where the α- and γ-protons of Mel$^{HA}$ appear. **B.** $^1$H NMR spectra from 1.8 to 2.1 ppm; the region where the β-protons of Mel$^{HA}$ appear.

*Attempted Prebiotic Synthesis of Mel$^{HA}$ with Acrolein and Acetone Cyanohydrin*

*1. Synthetic Protocol*

Because melamine co-precipitates with ferrocyanide, an alternative synthesis of Mel$^{HA}$ (via the corresponding cyanohydrin) was attempted with the transhydrocyanation reagent acetone cyanohydrin. To a 1 dram vial equipped with a stir bar, sodium formate buffer (1 M, pH 4, 1 mL) and melamine (12.6 mg, 0.1 mmol) were added. The suspension was stirred rapidly with mild heating until all of the melamine dissolved. Acetone cyanohydrin (36.6 µL, 0.4 mmol) was then added, followed by acrolein (13 µL, 0.2 mmol) was then added, and the vial was sealed and stirred in an oil bath at 45 °C for 24 hours. The reaction remained a clear solution. Samples were taken for UV-LC/MS analysis at 0, 2, 4, 6, and 24 hours. The reaction was then stopped and 1 mL of concentrated HCl was added. The solution was then refluxed for 24 hours,[2] after which it was lyophilized and redissolved in 600 µL $D_2O$ for NMR analysis.

### *2. UV-LC/MS Analysis*

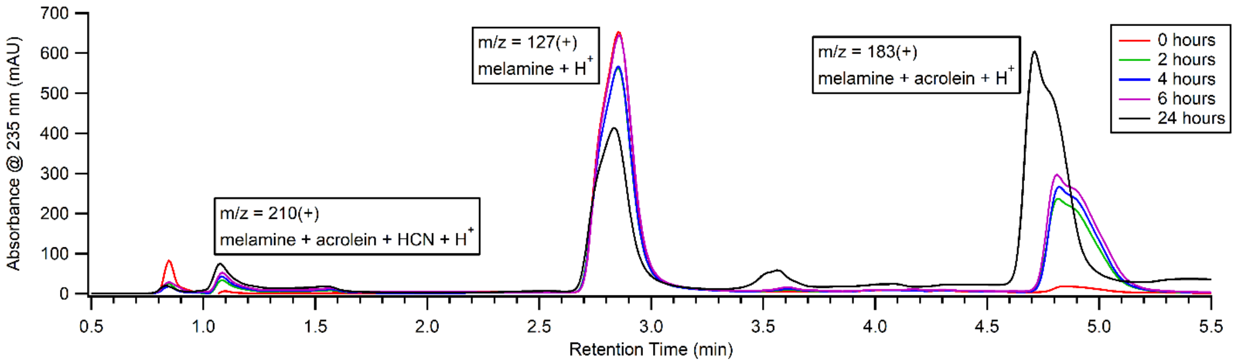

**Supplementary Figure 11.** UV-LC/MS monitoring of the reaction of melamine with acrolein and acetone cyanohydrin. The mass of the Mel$^{HA}$ cyanohydrin (210(+)) is detected.

### *3. NMR Analysis*

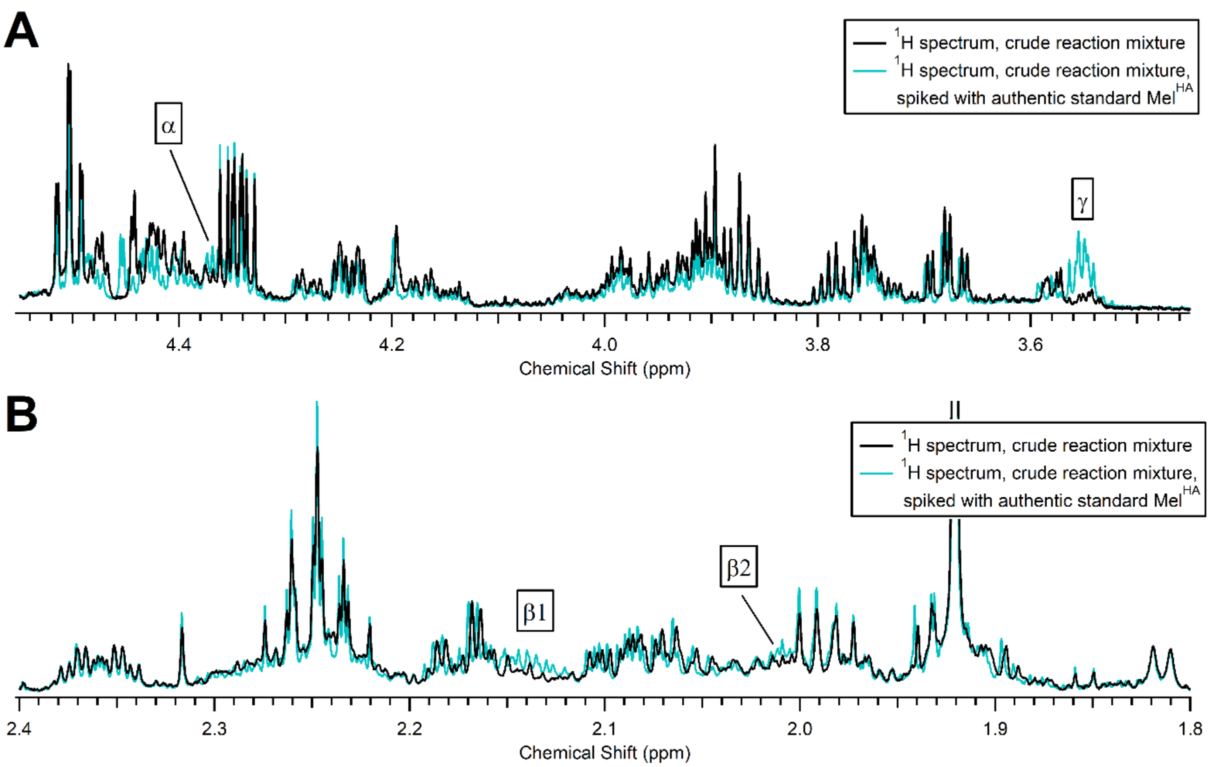

**Supplementary Figure 12.** $^{1}$H NMR analysis of the hydrolyzed reaction of melamine with acrolein and acetone cyanohydrin. The determination of the formation of Mel$^{HA}$ by spiking with an authentic standard is inconclusive, as the candidate product signals of Mel$^{HA}$ are weak and obscured by other signals. **A.** $^{1}$H NMR spectra from 3.45 to 4.55 ppm; the region where the α- and γ-protons of Mel$^{HA}$ appear. **B.** $^{1}$H NMR spectra from 1.8 to 2.4 ppm; the region where the β-protons of Mel$^{HA}$ appear.

*Reaction of Melamine with Acrolein*

*1. Synthetic Protocol*

Because the reaction of melamine with acrolein and acetone cyanohydrin only produced a small amount of cyanohydrin, melamine was reacted with acrolein in the absence of a cyanide source in order to determine if some property of these adducts (e.g., cyclization) inherently limit cyanohydrin formation. To a 1 dram vial equipped with a stir bar, water (300 μL), melamine (3.8 mg, 30 μmol), and formic acid (1.13 μL, 30 μmol) were added. The suspension was stirred rapidly with mild heating until all of the melamine dissolved. Acrolein (4 μL, 60 μmol) was then added, and the vial was sealed and stirred in an oil bath at 45 °C for 3 hours. Samples were taken for UV-LC/MS analysis at 0, 0.5, 1, 1.5, 2, 2.5, and 3 hours. The reaction was then stopped and 150 μL were removed, lyophilized, and redissolved in $D_2O$ for NMR analysis.

*2. UV-LC/MS Analysis*

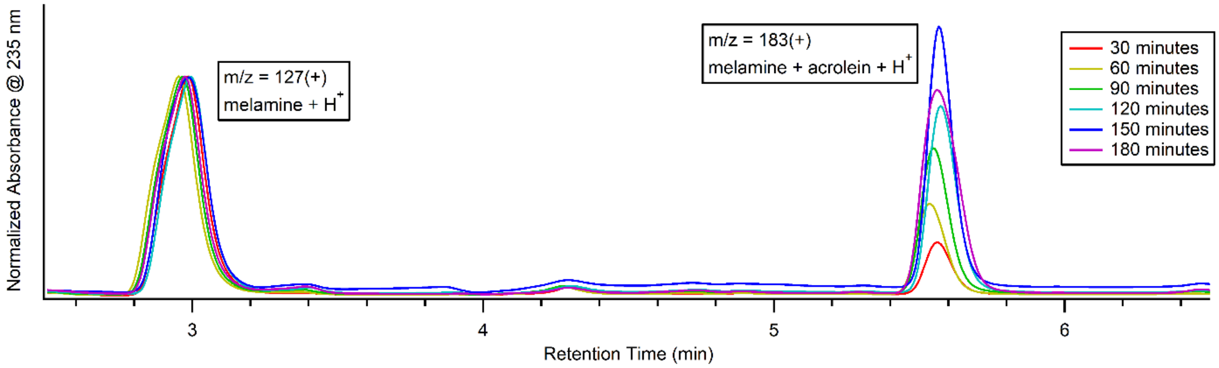

**Supplementary Figure 13.** UV-LC/MS monitoring of the reaction of melamine with acrolein. The traces have been normalized according to the height of the melamine peak. The mass of a melamine-acrolein adduct (183(+)) is detected.

*3. NMR Analysis*

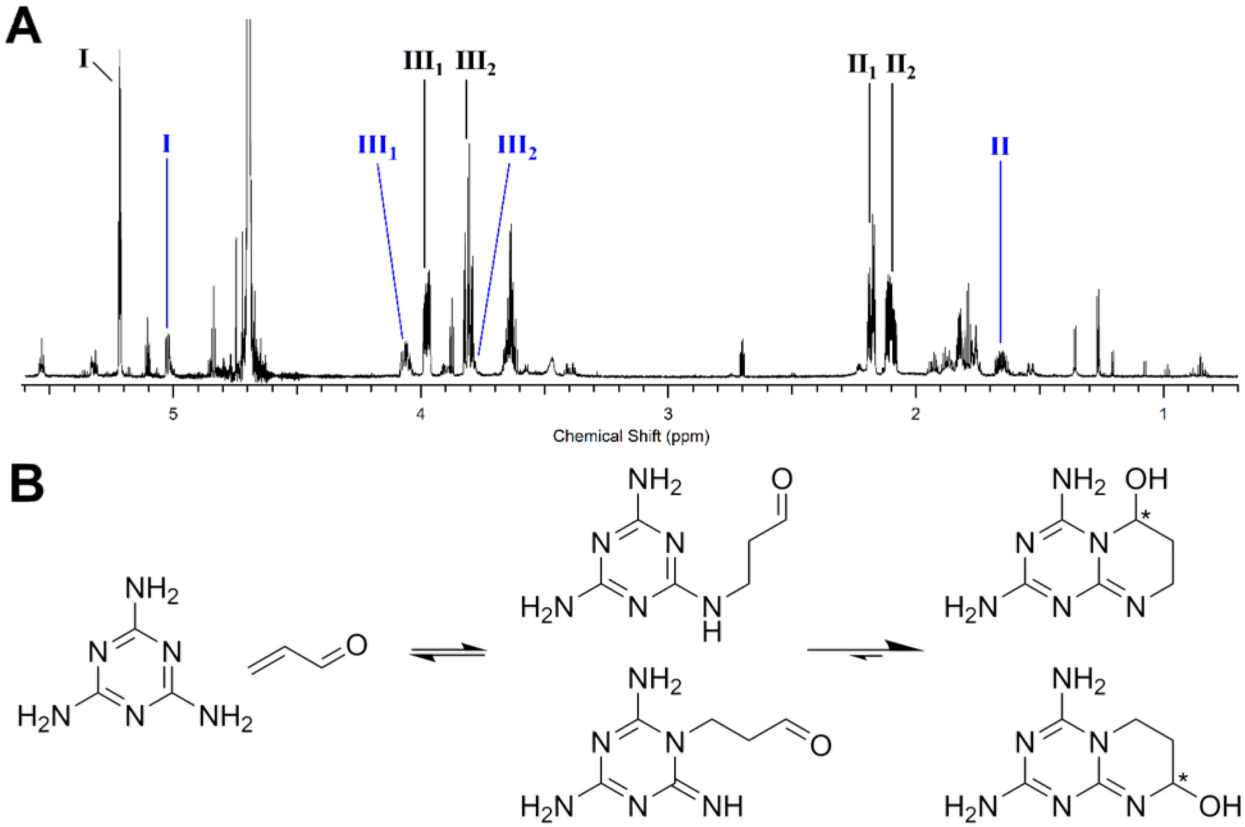

**Supplementary Figure 14.** [1]H NMR analysis of the reaction of melamine with acrolein. **A.** [1]H NMR spectrum of the reaction of melamine with acrolein from 0.7 to 5.6 ppm. Sets of coupled signals (determined by COSY) are indicated by label color. Roman numerals indicate the coupling pattern [I ↔ ($II_1$ ↔ $II_2$) ↔ ($III_1$ ↔ $III_2$)]. Arabic number subscripts indicate that the signal is a component of an AB spin system. **B.** Possible products formed by the reaction of melamine with acrolein. Cyclization without dehydration gives chiral hemiaminal moieties. Stereogenic centers are indicated with an asterick.

The appearance of AB signals in the [1]H NMR spectrum indicates the presence of a chiral center in the compound. This is possible by the formation of cyclic hemiaminal adducts of melamine and acrolein. Unlike cyanuric acid and adenine, all of the nucleophilic sites of melamine are part of a 1,3-binucleophilic motif; therefore, all conjugate adducts of melamine with acrolein are prone to cyclization. Cyclization sequesters the aldehyde moiety, inhibiting cyanohydrin formation.

*Prebiotic Synthesis of Mel$^{HA}$ from Thioammeline and α-Hydroxy-γ-aminobutyric acid*

*1. Synthetic Protocol*

To a 1 dram vial equipped with a stir bar, sodium carbonate buffer (1 M, pH 10, 1 mL), thioammeline (14.3 mg, 0.1 mmol), and (*S*)-α-hydroxy-γ-aminobutyric acid (14.9 mg, 0.125 mmol) were added. The vial was sealed and stirred in an oil bath at 100 °C for 7 days. The reaction was then lyophilized and redissolved in 1 mL $D_2O$ with DSS (10 mM proton signal) as an internal standard for $^1$H NMR analysis.

*2. NMR Analysis*

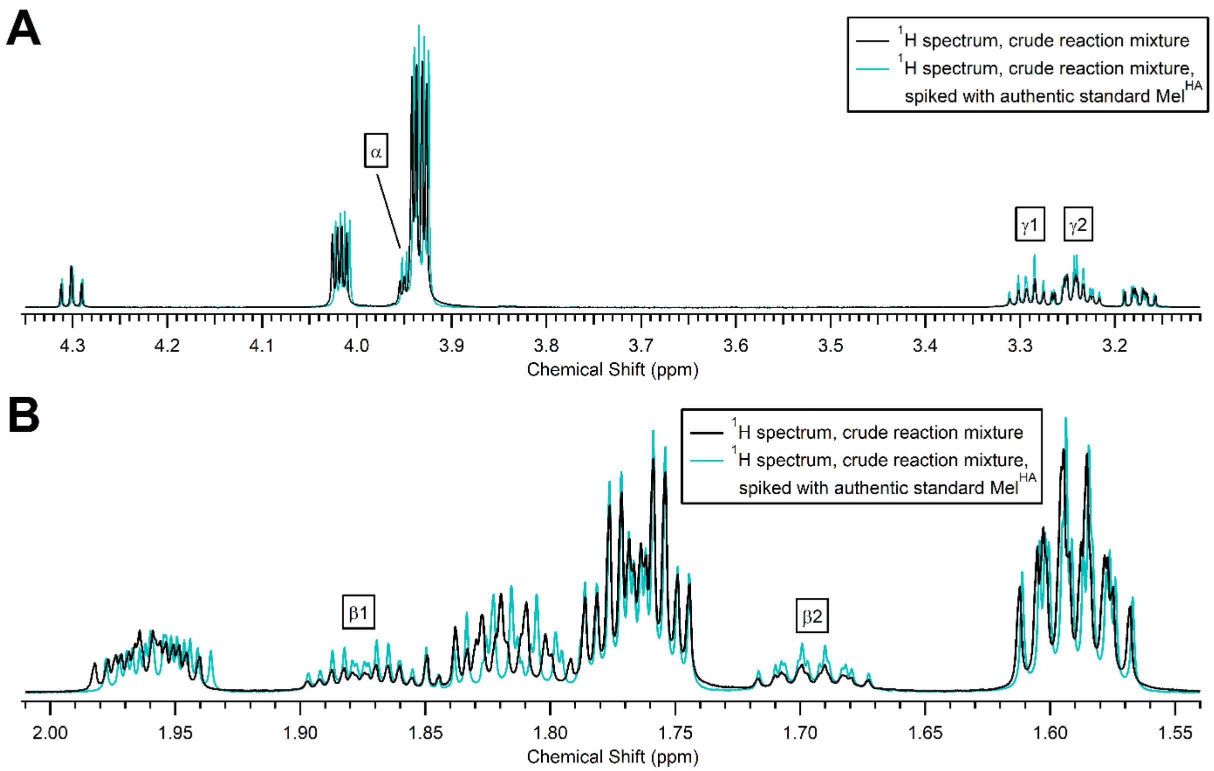

**Supplementary Figure 15.** $^1$H NMR analysis of the reaction of thioammeline with α-hydroxy-γ-aminobutyric acid. The successful formation of Mel$^{HA}$ is indicated by the increase in signal intensity, without the formation of new signals, of the aliphatic protons of Mel$^{HA}$ upon spiking with an authentic standard. **A.** $^1$H NMR spectra from 3.11 to 4.35 ppm; the region where the α- and γ-protons of Mel$^{HA}$ appear. The apparent increases in intensity of the signals at 4.02 ppm and 3.93 ppm are due to peak sharpening from better shimming in the analysis of the spiked sample. **B.** $^1$H NMR spectra from 1.54 to 2.01 ppm; the region where the β-protons of Mel$^{HA}$ appear. The signals at ca. 1.97 ppm and 1.82 ppm are shifted upfield slightly due to a slight change in pD upon the addition of the Mel$^{HA}$ authentic standard.

*Prebiotic Syntheses of Cy^HA^-D, Ad^HA^-D, and Mel^HA^-D*

*1. Synthetic Protocol*

Stock solutions of *rac*-$Cy^{HA}$ and L-aspartic acid were prepared at 62.5 mM in deionized water. Stock solutions of *rac*-$Ad^{HA}$ and (*S*)-$Mel^{HA}$ were prepared at 25 mM in deionized water and were kept hot to maintain solubility. A stock solution of pyridine was prepared at 500 mM in deionized water.

Reaction samples were prepared in PCR tubes so that each contained 0.5 µmol of $Nu^{HA}$ (Nu = Cy, Mel, Ad), 0.5 µmol of L-aspartic acid, and 0 or 4 µmol pyridine. For reactions containing two $Nu^{HA}$ species, 0.25 µmol of each were used. The reaction solutions were left uncapped and placed in an oven at 85 °C. The samples were rehydrated once daily with 20 µL of an aqueous pyridine solution (0.2 M). The samples were briefly heated and vortexed to bring the components into solution. For some samples containing $Mel^{HA}$ and $Ad^{HA}$, full re-dissolution could not be achieved, and the samples were left as fine suspensions. The samples were then placed back in the 85 °C oven uncapped, initiating the next dry phase. The reactions were dried 7 times total. After the final dry phase, the samples were redissolved in 50 µL deionized water and analyzed by UV-LC/MS.

*2. UV-LC/MS Analysis*

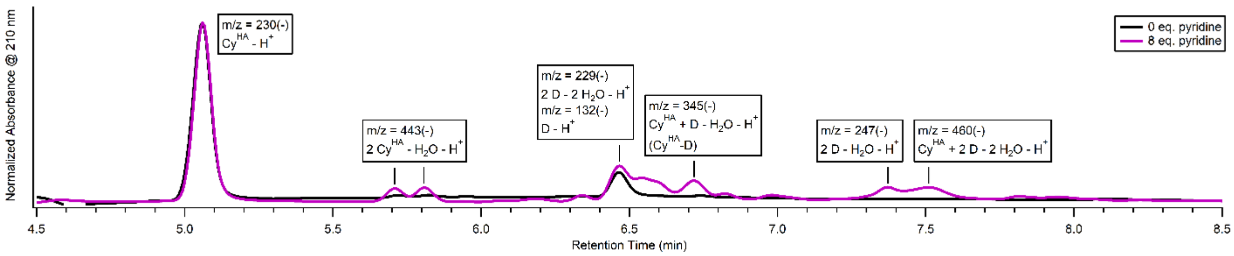

**Supplementary Figure 16.** UV-LC/MS monitoring (210 nm) of the wet-dry cycling reaction of $Cy^{HA}$ with aspartic acid (D) and 0 equivalents of pyridine (black trace) or 8 equivalents of pyridine (purple trace) after 7 cycles. The spectra have been scaled according to the height of the peak corresponding to $Cy^{HA}$.

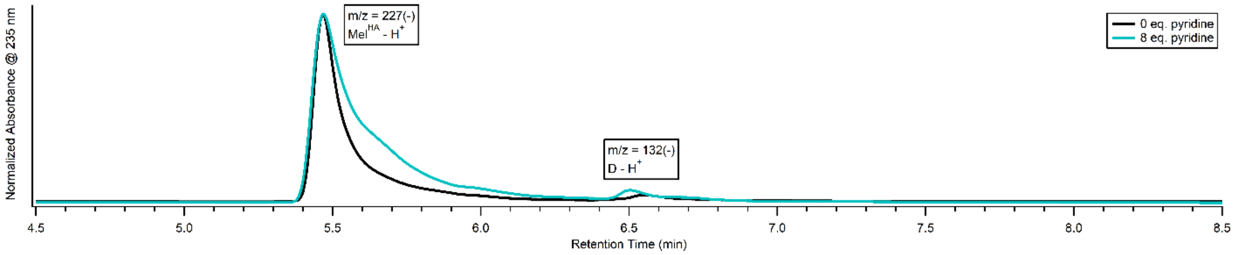

**Supplementary Figure 17.** UV-LC/MS monitoring (235 nm) of the wet-dry cycling reaction of $Mel^{HA}$ with aspartic acid (D) and 0 equivalents of pyridine (black trace) or 8 equivalents of pyridine (teal trace) after 7 cycles. The spectra have been scaled according to the height of the peak corresponding to $Mel^{HA}$.

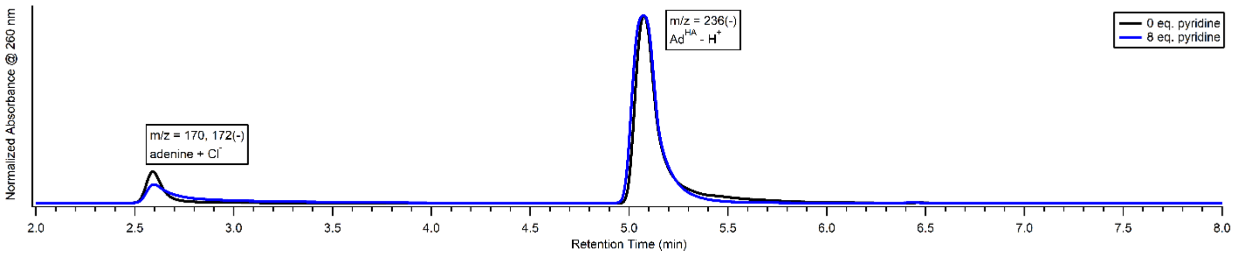

**Supplementary Figure 18.** UV-LC/MS monitoring (260 nm) of the wet-dry cycling reaction of Ad$^{HA}$ with aspartic acid (D) and 0 equivalents of pyridine (black trace) or 8 equivalents of pyridine (blue trace) after 7 cycles. The spectra have been scaled according to the height of the peak corresponding to Ad$^{HA}$.

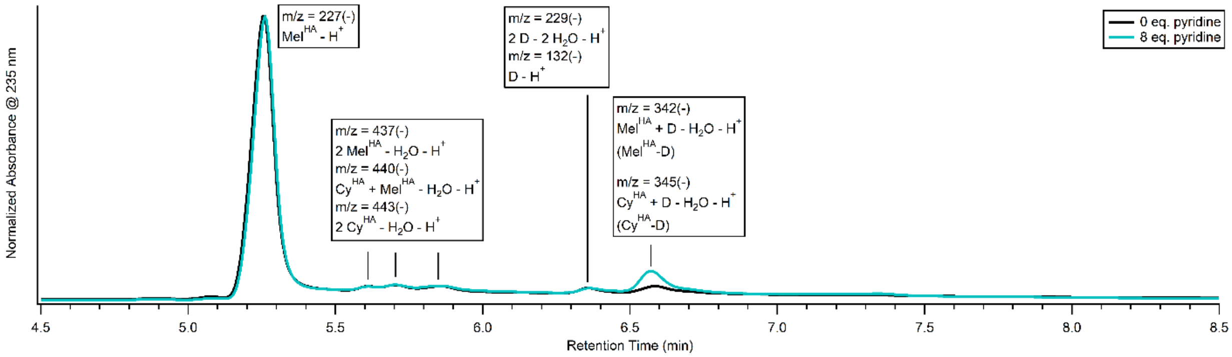

**Supplementary Figure 19.** UV-LC/MS monitoring (235 nm, melamine absorbance shoulder) of the wet-dry cycling reaction of Cy$^{HA}$ and Mel$^{HA}$ with aspartic acid (D) and 0 equivalents of pyridine (black trace) or 8 equivalents of pyridine (teal trace) after 7 cycles. The spectra have been scaled according to the height of the peak corresponding to Mel$^{HA}$. Note that, at 235 nm, the absorbance of species containing only Cy$^{HA}$ is lower than those containing Mel$^{HA}$.

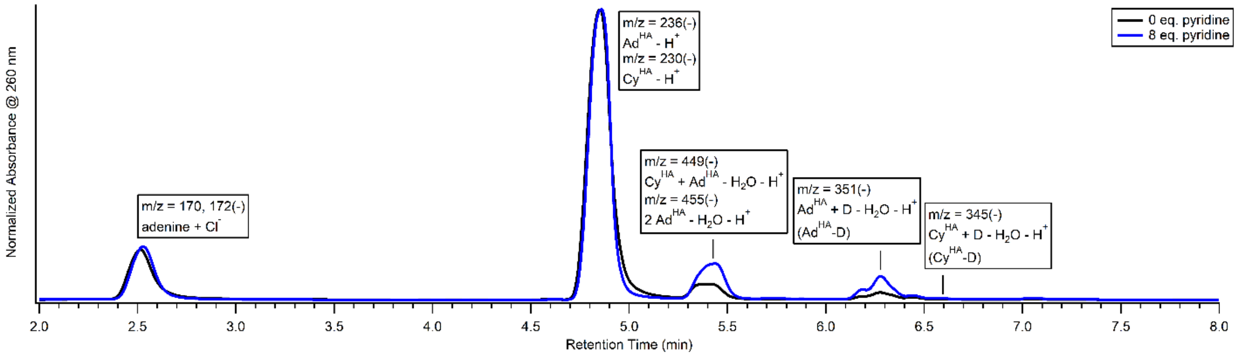

**Supplementary Figure 20.** UV-LC/MS monitoring (260 nm, adenine absorbance maximum) of the wet-dry cycling reaction of Cy$^{HA}$ and Ad$^{HA}$ with aspartic acid (D) and 0 equivalents of pyridine (black trace) or 8 equivalents of pyridine (blue trace) after 7 cycles. Note that, at 260 nm, the absorbance of species containing only Cy$^{HA}$ is lower than those containing Ad$^{HA}$.

### IV. Nu$^{HA}$-D Prebiotic Oligomerization Experiments[1]

*Heterocyclic Additive Survey in Dry-Down Oligomerization Reactions*

*1. Reaction Protocol*

Stock solutions of (*S*,*S*)-Cy$^{HA}$-D, (*S*,*S*)-Mel$^{HA}$-D, and (*S*,*S*)-Ad$^{HA}$-D were prepared at 312.5 mM in deionized water, and stock solutions of heterocyclic compounds (pyridine, 2-hydroxypyridine, or 2-mercaptopyridine) were prepared at 2.5 M, 1.25 M, 625 mM, and 312.5 mM in deionized water. To a PCR tube, 1.6 μL of Nu$^{HA}$-D (Nu = Cy, Mel, Ad) solution (or 0.8 μL of each solution for reactions containining two Nu$^{HA}$-D species) and 0.4 μL of the appropriate stock solution of heterocyclic compound were added to give 2, 1, 0.5, or 0.25 molar equivalents of heterocyclic additive. In the case of 4 eq. of heterocyclic additive, 0.8 μL of the 2.5 M solution were added. In the case of 0 eq. heterocyclic additive, 0.4 μL water were added. This gave a final Nu$^{HA}$-D concentration of 250 mM (or 208.3 mM for the reactions containing 4 eq. of heterocyclic additive). Reactions typically had starting pH values around 2-3 due to the carboxylic acid moieties of Nu$^{HA}$-D. With the caps open, the PCR tubes were transferred to an oven at 85 °C and left to dry and react for 1 week. No intermittent rehydration was performed. After 1 week, the residue was redissolved in 20 μL deionized water and analyzed by UV-LC/MS.

*2. UV-LC/MS Analysis*

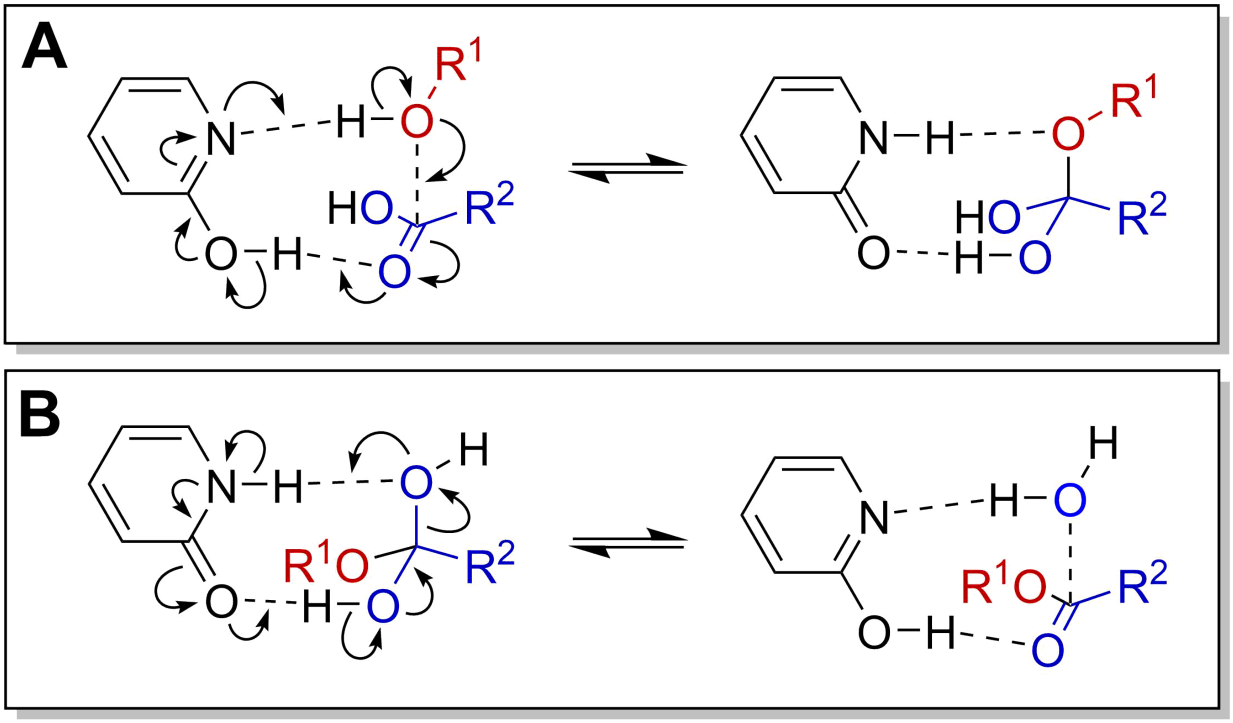

**Supplementary Figure 21.** The ability of 2-hydroxypyridine to enhance the extent of oligomerization with the greatest efficiency (at a 1 molar equivalent loading; see figures below) may be due to its ability to act as a bifunctional catalyst. The elementary reactions steps corresponding to **A.** attack of the alcohol substrate (heavy atoms shown in red) on the carboxylic acid substrate (heavy atoms shown in blue) and **B.** subsequent expulsion of water from the rearranged tetrahedral intermediate may be catalyzed by 2-hydroxypyridine.

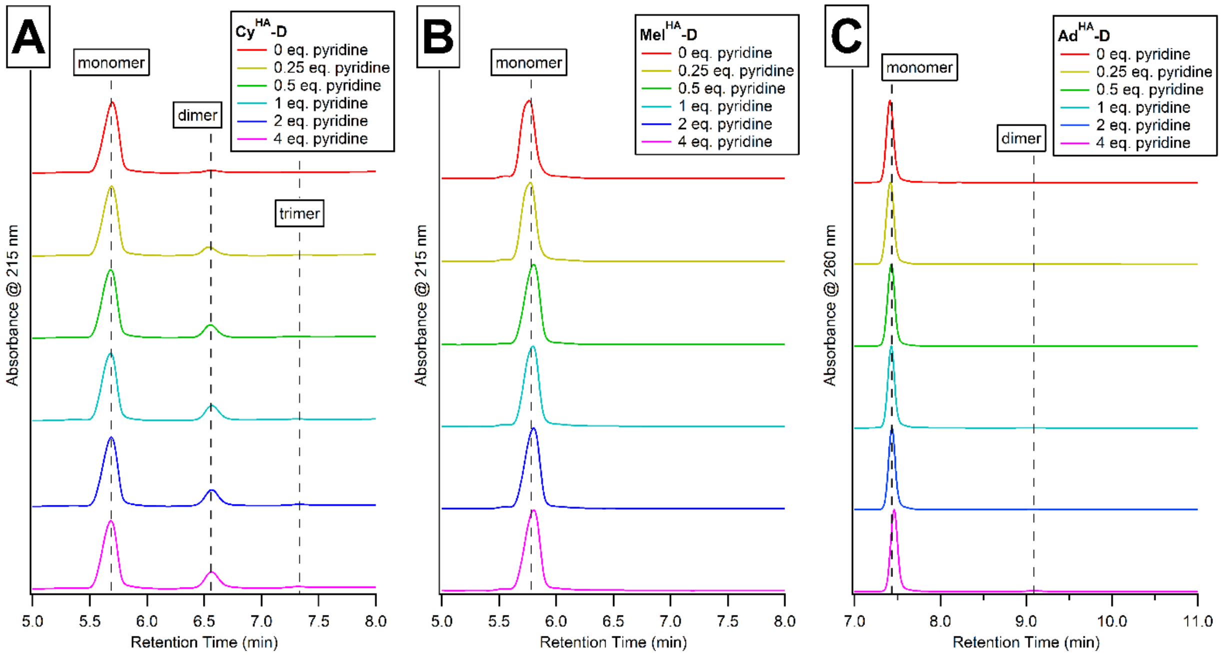

**Supplementary Figure 22.** UV-LC/MS analysis of Nu$^{HA}$-D oligomerization reactions at 85 °C after one week with variable loadings of pyridine. **A.** Cy$^{HA}$-D oligomerization chromatograms. **B.** Mel$^{HA}$-D oligomerization chromatograms. **C.** Ad$^{HA}$-D oligomerization chromatograms.

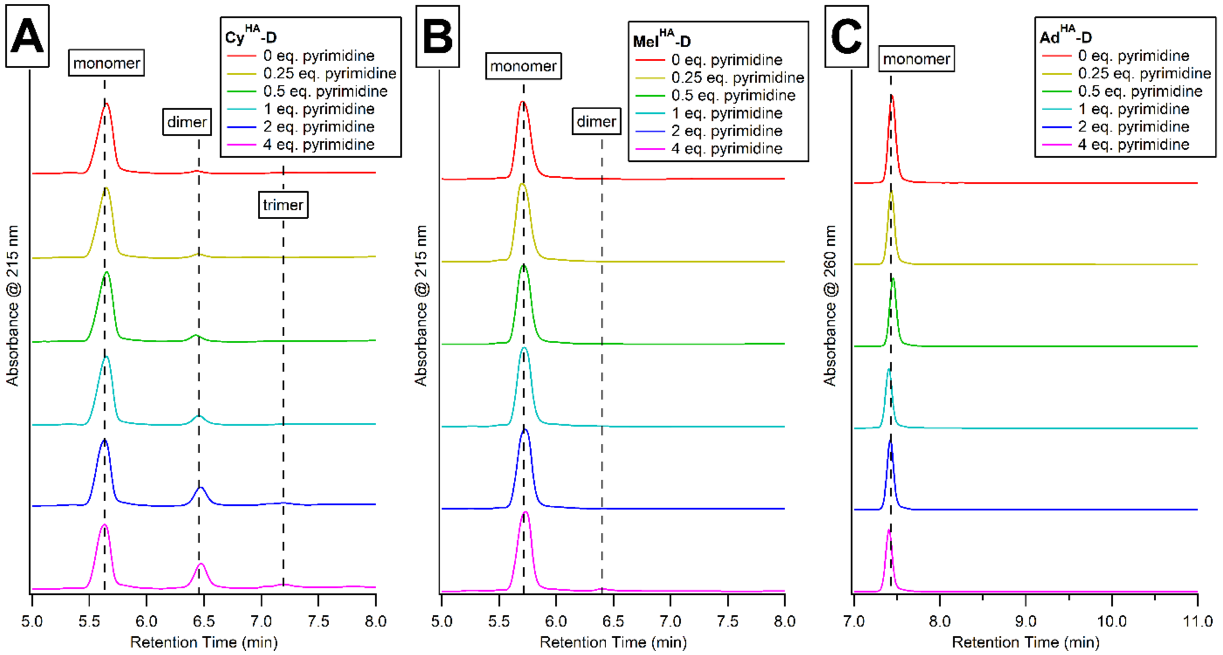

**Supplementary Figure 23.** UV-LC/MS analysis of Nu$^{HA}$-D oligomerization reactions at 85 °C after one week with variable loadings of pyrimidine. **A.** Cy$^{HA}$-D oligomerization chromatograms. **B.** Mel$^{HA}$-D oligomerization chromatograms. **C.** Ad$^{HA}$-D oligomerization chromatograms.

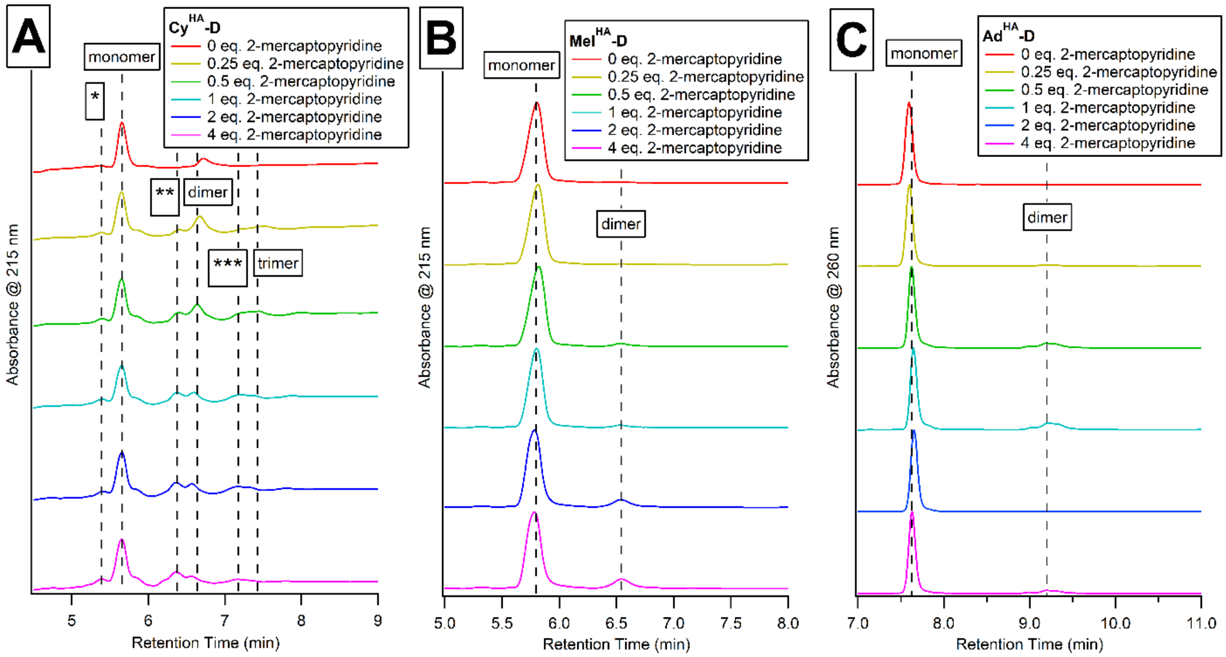

**Supplementary Figure 24.** UV-LC/MS analysis of Nu$^{HA}$-D oligomerization reactions at 85 °C after one week with variable loadings of 2-mercaptopyridine. **A.** Cy$^{HA}$-D oligomerization chromatograms. *Product with mass (Cy$^{HA}$-D) + D – 2 $H_2O$. **Product with mass $(Cy^{HA}\text{-}D)_2$ + D – 2 $H_2O$. ***Product with mass $(Cy^{HA}\text{-}D)_3$ + D – 2 $H_2O$. **B.** Mel$^{HA}$-D oligomerization chromatograms. **C.** Ad$^{HA}$-D oligomerization chromatograms.

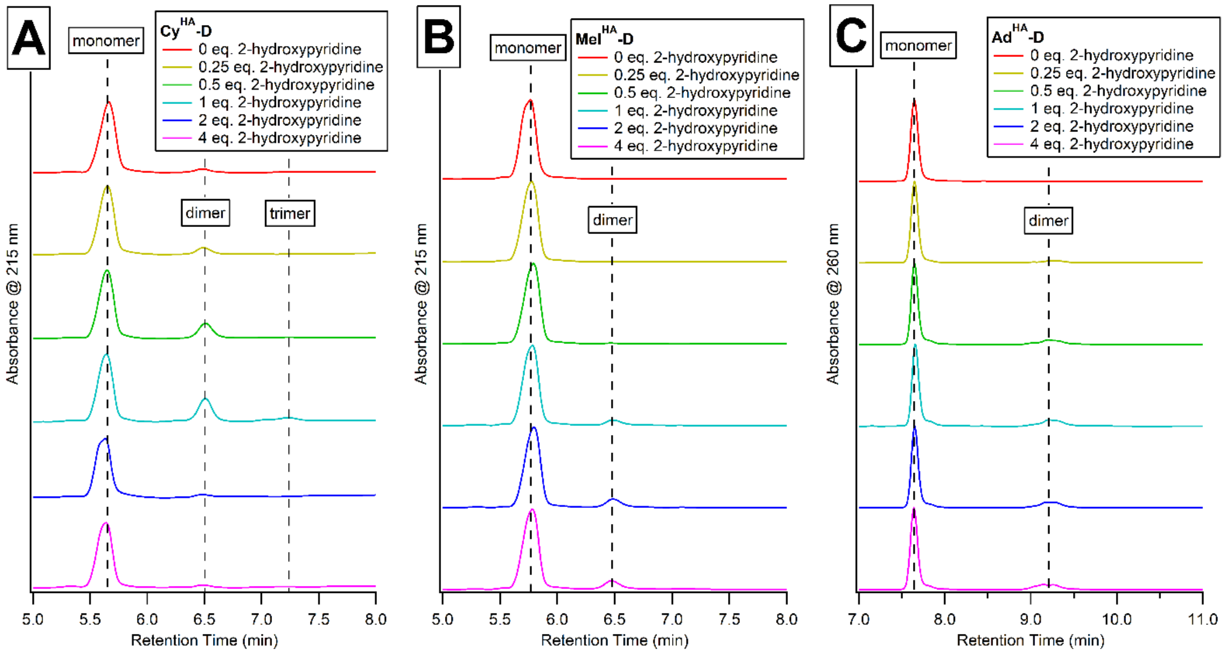

**Supplementary Figure 25.** UV-LC/MS analysis of Nu$^{HA}$-D oligomerization reactions at 85 °C after one week with variable loadings of 2-hydroxypyridine. **A.** Cy$^{HA}$-D oligomerization chromatograms. **B.** Mel$^{HA}$-D oligomerization chromatograms. **C.** Ad$^{HA}$-D oligomerization chromatograms.

*Long-Term Dry-Down Oligomerization Reactions*

*1. Reaction Protocol*

Stock solutions of (*S*,*S*)-Cy$^{HA}$-D, (*S*,*S*)-Mel$^{HA}$-D, and (*S*,*S*)-Ad$^{HA}$-D were prepared at 312.5 mM in deionized water, and a stock solution of 2-hydroxypyridine was prepared at 1.25 M in deionized water. To a PCR tube, 1.6 µL of Nu$^{HA}$-D (Nu = Cy, Mel, Ad) solution (or 0.8 µL of each solution for reactions containining two Nu$^{HA}$-D species) and 0.4 µL of 2-hydroxypyridine solution were added to give final Nu$^{HA}$-D and 2-hydroxypyridine concentrations of 250 mM. Reactions typically had starting pH values around 2-3 due to the carboxylic acid moieties of Nu$^{HA}$-D. With the caps open, the PCR tubes were transferred to an oven at 85 °C and left to dry and react for 4 or 8 weeks. No intermittent rehydration was performed. After the specified time, the residue was redissolved in 20 µL deionized water and analyzed by UV-LC/MS.

*2. UV-LC/MS Analysis*

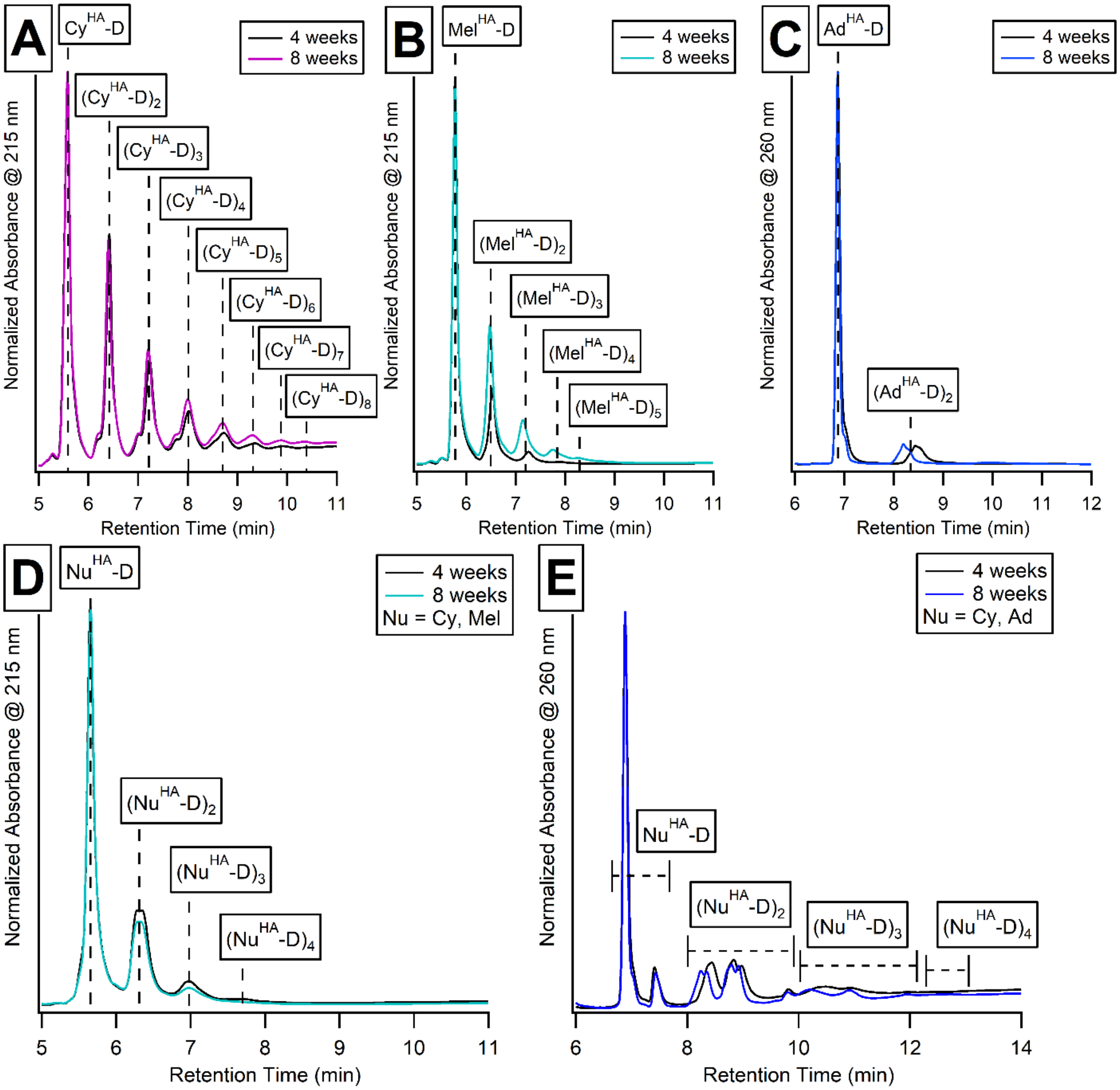

**Supplementary Figure 26.** UV-LC/MS analysis of Nu$^{HA}$-D oligomerization reactions at 85 °C after 4 or 8 weeks with 1 equivalent of 2-hydroxypyridine. **A.** Cy$^{HA}$-D oligomerization chromatograms. **B.** Mel$^{HA}$-D oligomerization chromatograms. **C.** Ad$^{HA}$-D oligomerization chromatograms. **D.** Cy$^{HA}$-D/Mel$^{HA}$-D co-oligomerization chromatograms. **E.** Cy$^{HA}$-D/Ad$^{HA}$-D co-oligomerization chromatograms. See main text for specific peak assignments.

### V. Self-Assembly Experiments

*Minimum Assembly Concentration Experiments*

As described previously, a $^{1}$H NMR-based method was used to determine how efficiently the compounds studied self-assemble in water. Briefly, below the minimum assembly concentration (MAC), no assembly occurs, and the concentration of assembling molecules, as measured by NMR, matches the true concentration. However, above the MAC, self-assembly occurs, forming large aggregates that have baseline-broadened NMR signals that cannot be integrated. Therefore, the NMR-measured concentration of assembling molecules above the MAC will always report the MAC value, representing the concentration of molecules still free in solution. All experiments were performed with equal concentrations of cyanuric acid-containing compound and melamine-containing compound, except for $(Cy^{HA}\text{-}D)_2$, in which the concentration of melamine was double the concentration of $(Cy^{HA}\text{-}D)_2$, but equal to the concentration of cyanuric acid moieties. All experiments were performed on a Bruker 800 MHz instrument with a 200 µL sample in a 3 mm tube with a DSS (2,2-dimethyl-2-silapentane-5-sulfonate-$d_6$ sodium salt, CAS 284664-85-3) internal standard at 10 mM. Unless stated otherwise, all spectra were collected at 25 °C as the sum of 64 scans.

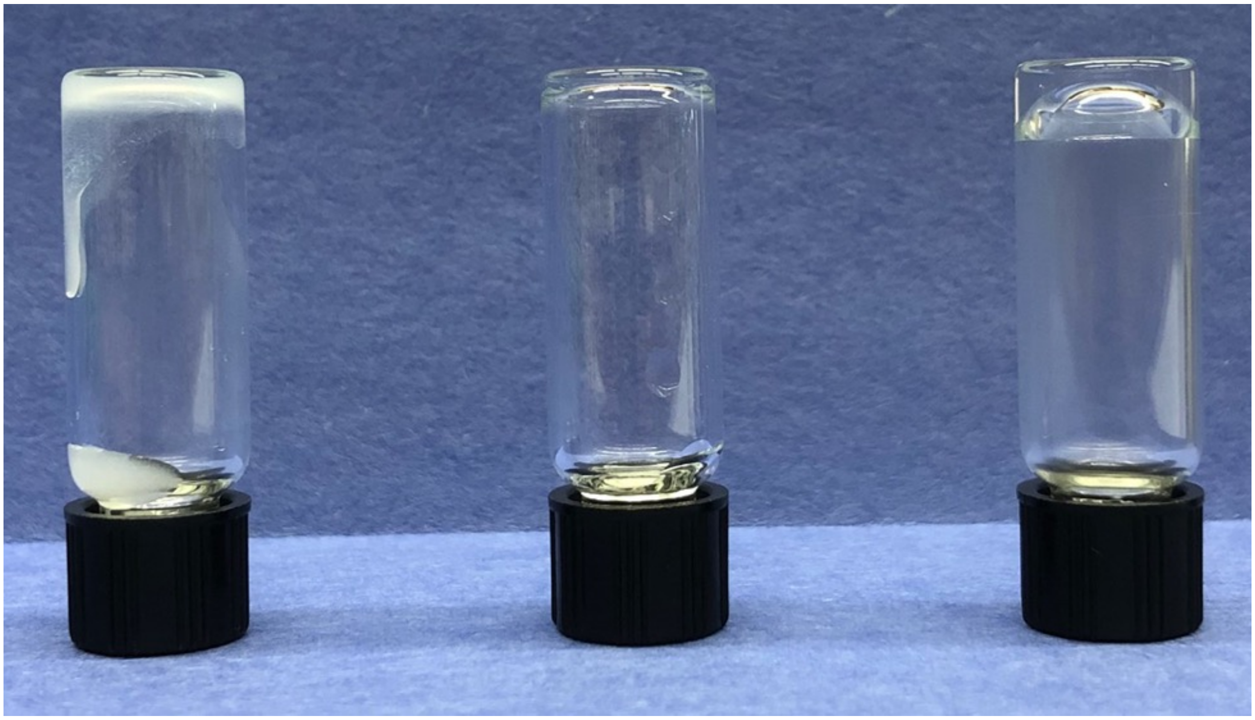

**Supplementary Figure 27.** Vial inversion test for samples containing $Cy^{HA}$-D or $Mel^{HA}$-D with a pairing partner in 100 mM N-methylmorpholine DCl buffer pD 7.5 with magnesium chloride 25 mM. **Left:** $Cy^{HA}$-D with melamine, 50 mM. Precipitation occurs without gel formation. Because this mixture does not form soluble assemblies, the MAC was not determined. **Middle:** $Cy^{HA}$-D with $Mel^{HA}$-D, 50 mM. Neither precipitation nor gelation occurs. **Right:** $Mel^{HA}$-D with cyanuric acid, 50 mM. Gelation occurs without precipitation.

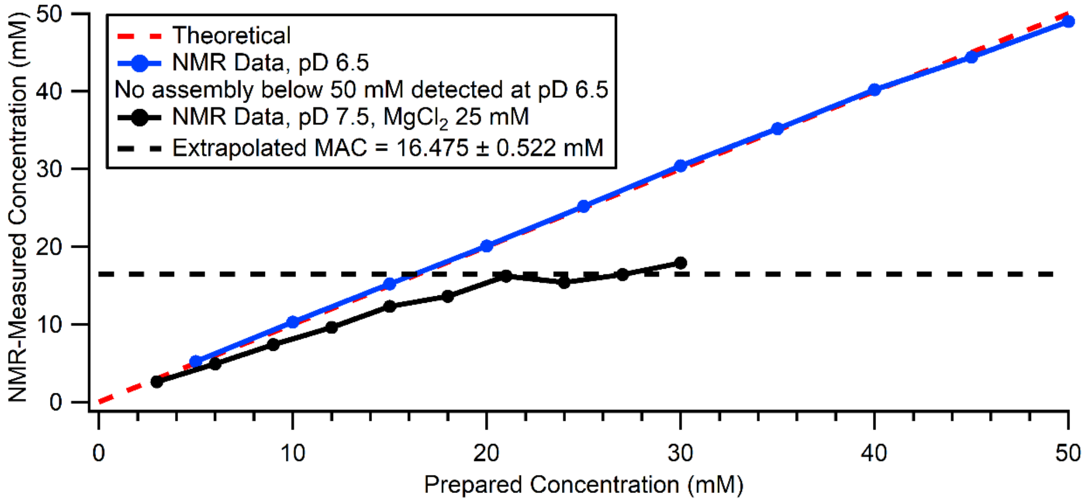

**Supplementary Figure 28.** MAC determination of Mel$^{HA}$-D with cyanuric acid. **Blue trace:** 2,6-lutidine DCl buffer, 100 mM, pD 6.5. No assembly below 50 mM was detected under these conditions. **Black trace:** N-methylmorpholine DCl buffer, 100 mM, pD 7.5, with $MgCl_2$ 25 mM. The extrapolated MAC is 16.5 mM.

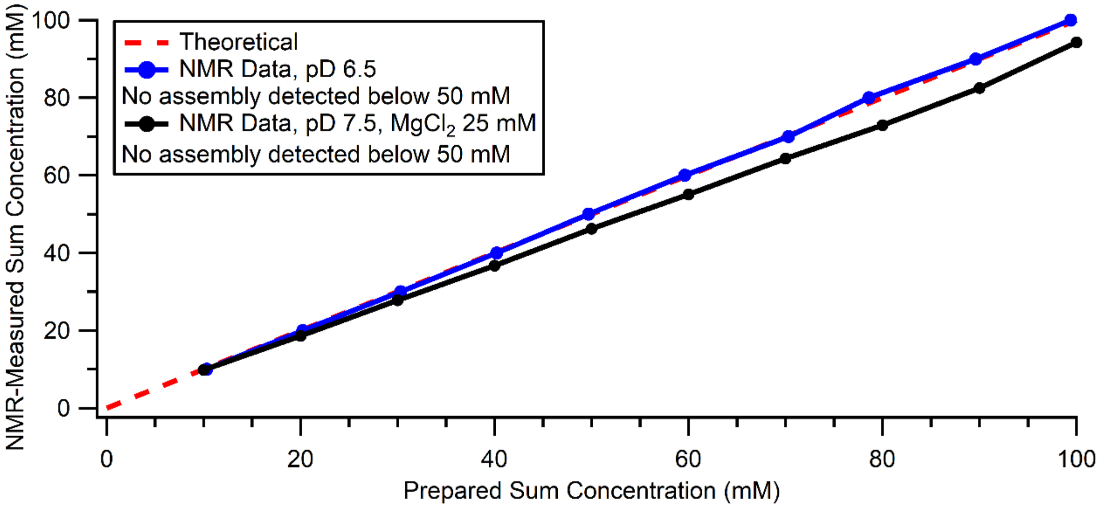

**Supplementary Figure 29.** MAC determination of Cy$^{HA}$-D with Mel$^{HA}$-D. The reported concentrations represent the sums of the concentrations of Cy$^{HA}$-D and Mel$^{HA}$-D. **Blue trace:** 2,6-lutidine DCl buffer, 100 mM, pD 6.5. No assembly below 50 mM was detected under these conditions. **Black trace:** N-methylmorpholine DCl buffer, 100 mM, pD 7.5, with $MgCl_2$ 25 mM. No assembly below 50 mM was detected under these conditions.

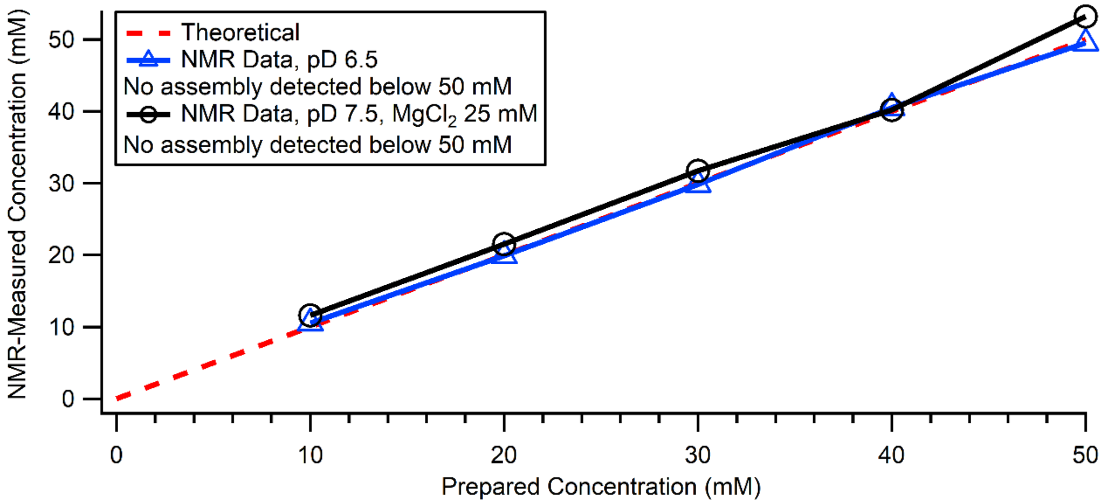

**Supplementary Figure 30.** MAC determination of Cy[HA]-D with adenine. **Blue trace:** 2,6-lutidine DCl buffer, 100 mM, pD 6.5. No assembly below 50 mM was detected under these conditions. **Black trace:** N-methylmorpholine DCl buffer, 100 mM, pD 7.5, with $MgCl_2$ 25 mM. No assembly below 50 mM was detected under these conditions.

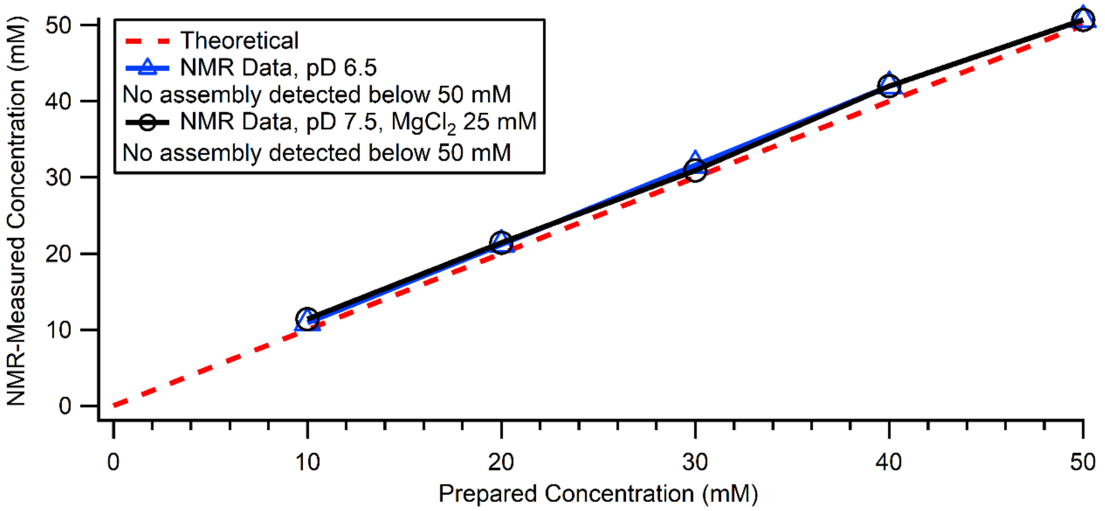

**Supplementary Figure 31.** MAC determination of Ad[HA]-D with cyanuric acid. **Blue trace:** 2,6-lutidine DCl buffer, 100 mM, pD 6.5. No assembly below 50 mM was detected under these conditions. **Black trace:** N-methylmorpholine DCl buffer, 100 mM, pD 7.5, with $MgCl_2$ 25 mM. No assembly below 50 mM was detected under these conditions.

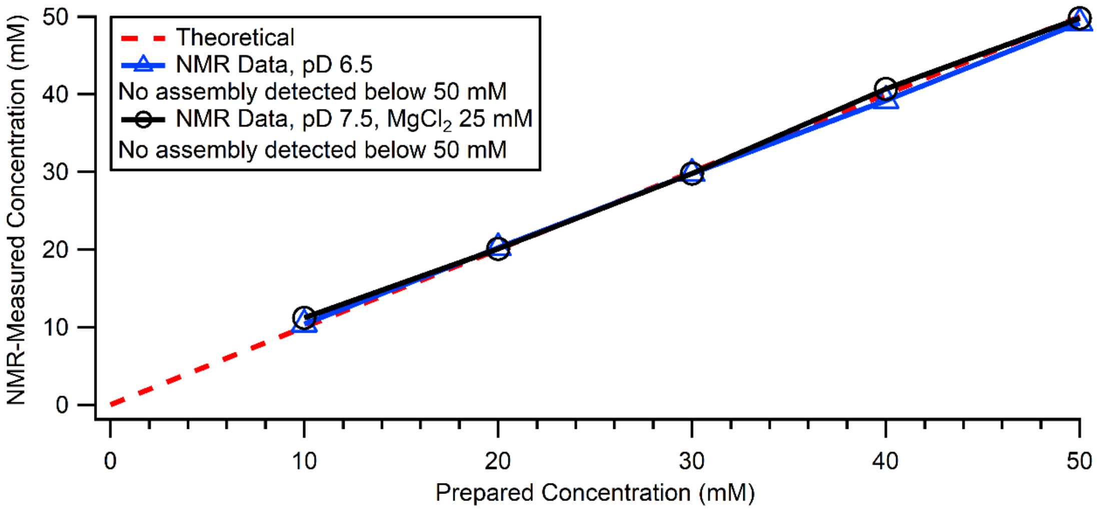

**Supplementary Figure 32.** MAC determination of $Cy^{HA}$-D with $Ad^{HA}$-D. The reported concentrations represent those measured for $Cy^{HA}$-D. **Blue trace:** 2,6-lutidine DCl buffer, 100 mM, pD 6.5. No assembly below 50 mM was detected under these conditions. **Black trace:** N-methylmorpholine DCl buffer, 100 mM, pD 7.5, with $MgCl_2$ 25 mM. No assembly below 50 mM was detected under these conditions.

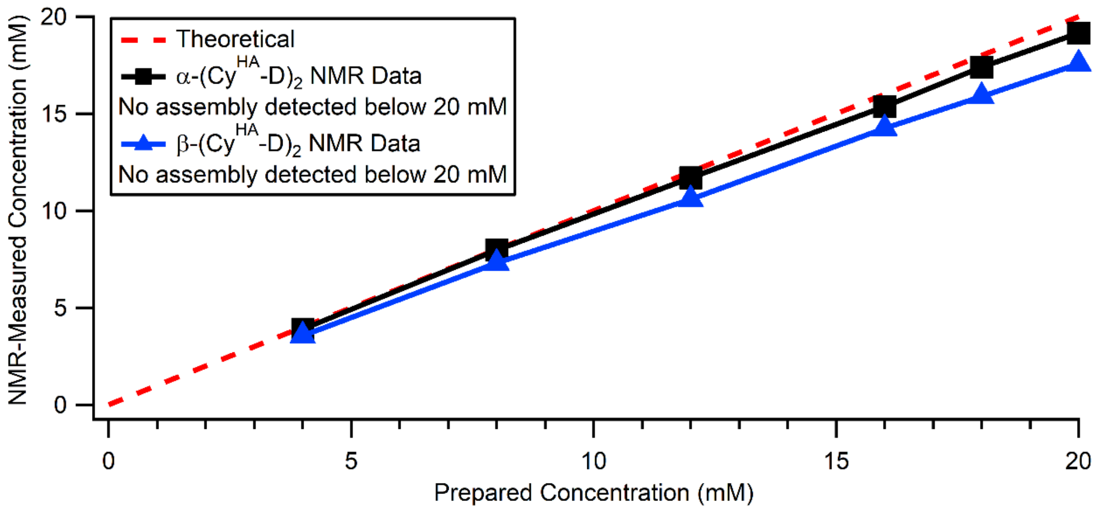

**Supplementary Figure 33.** MAC determination of $(Cy^{HA}\text{-D})_2$ with adenine. Reported concentrations represent those measured for $(Cy^{HA}\text{-D})_2$; adenine was prepared at a 2-fold greater concentration. Experiment performed at 5 °C. **Black trace:** α-$(Cy^{HA}\text{-D})_2$, 2,6-lutidine DCl buffer, 100 mM, pD 6.5. No assembly below 20 mM was detected under these conditions. **Blue trace:** β-$(Cy^{HA}\text{-D})_2$, 2,6-lutidine DCl buffer, 100 mM, pD 6.5. No assembly below 20 mM was detected under these conditions.

*Atomic Force Microscopy Imaging*

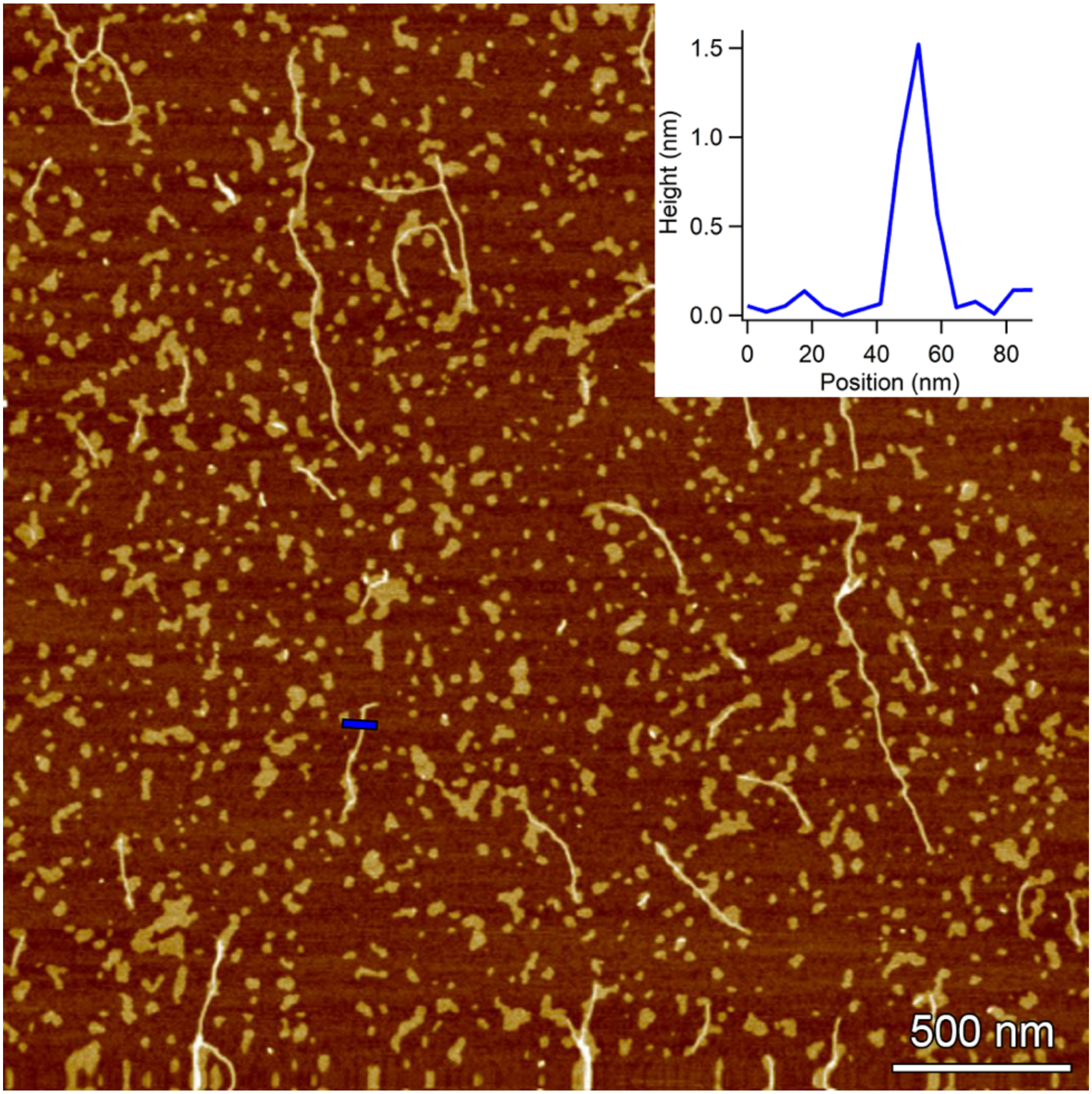

**Supplementary Figure 34.** AFM topographical image of assembly between Cy$^{HA}$-D and melamine, 50 mM, in lutidine DCl buffer, 100 mM, pD 6.5 ($D_2O$), pD-adjusted with triethylamine and DCl. Inset shows height profile delineated by blue line in main panel.

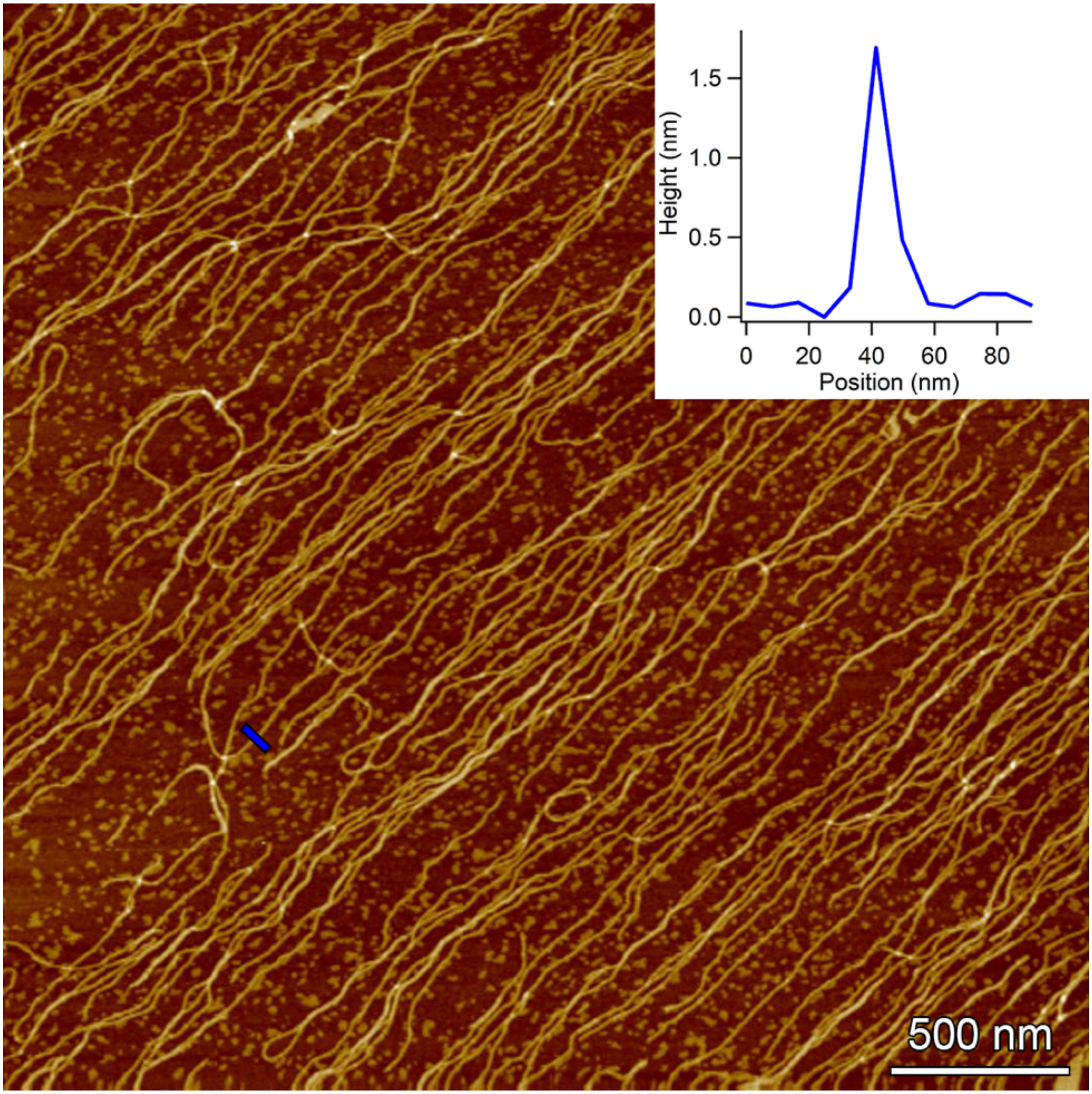

**Supplementary Figure 35.** AFM topographical image of assembly between Cy$^{HA}$-D and melamine, 50 mM, in *N*-methylmorpholine DCl buffer, 100 mM, pD 7.5 ($D_2O$), with $MgCl_2$ 25 mM, and pD-adjusted with triethylamine and DCl. Inset shows height profile delineated by blue line in main panel.

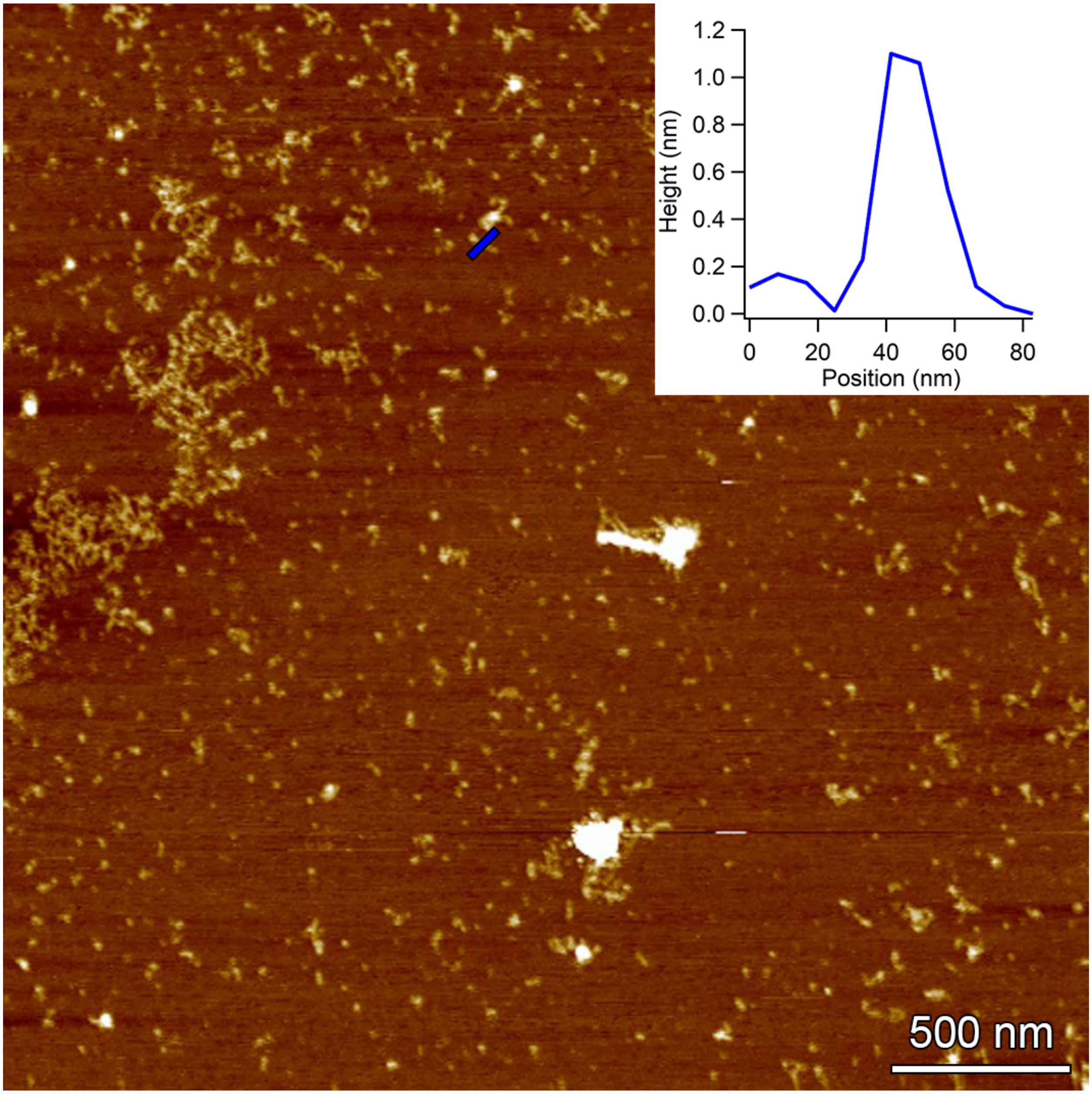

**Supplementary Figure 36.** AFM topographical image of assembly between Mel$^{HA}$-D and cyanuric acid, 50 mM, in lutidine DCl buffer, 100 mM, pD 6.5 ($D_2O$), pD-adjusted with triethylamine and DCl. Inset shows height profile delineated by blue line in main panel.

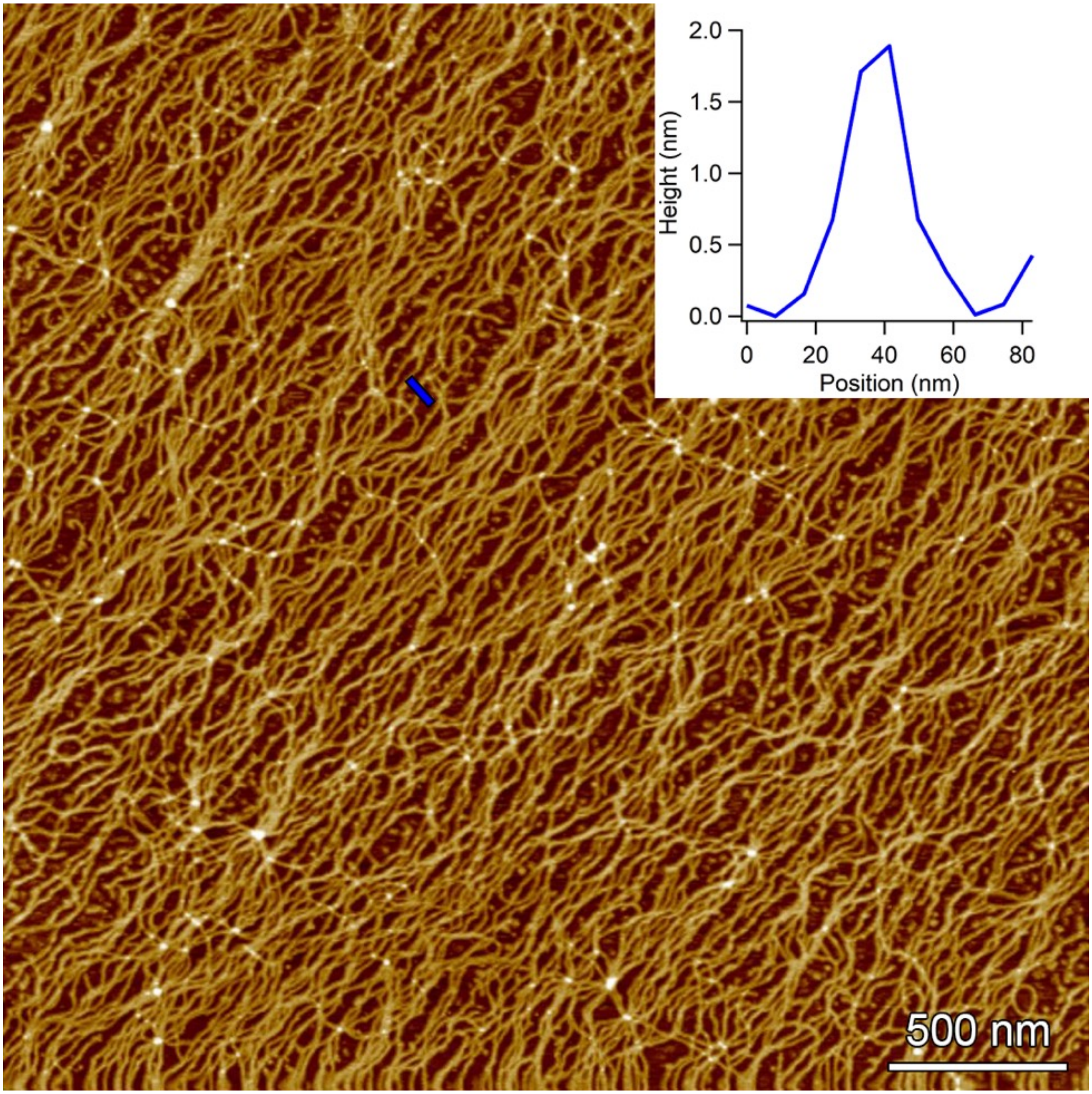

**Supplementary Figure 37.** AFM topographical image of assembly between Mel$^{HA}$-D and cyanuric, 50 mM, in *N*-methylmorpholine DCl buffer, 100 mM, pD 7.5 ($D_2O$), with $MgCl_2$ 25 mM, and pD-adjusted with triethylamine and DCl. Inset shows height profile delineated by blue line in main panel.

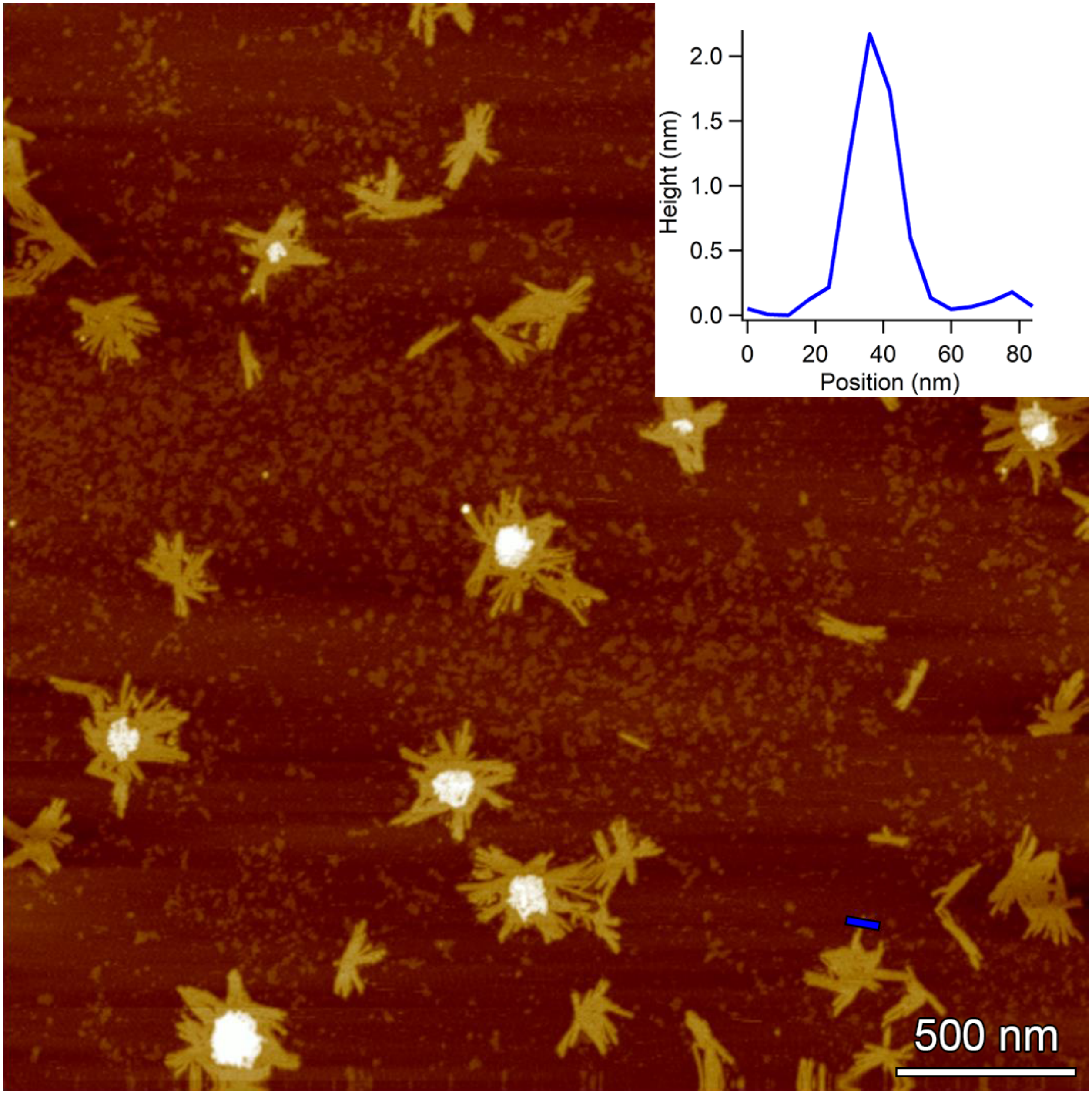

**Supplementary Figure 38.** AFM topographical image of assembly between Cy$^{HA}$-D and Mel$^{HA}$-D, 50 mM, in lutidine DCl buffer, 100 mM, pD 6.5 ($D_2O$), pD-adjusted with triethylamine and DCl. Inset shows height profile delineated by blue line in main panel.

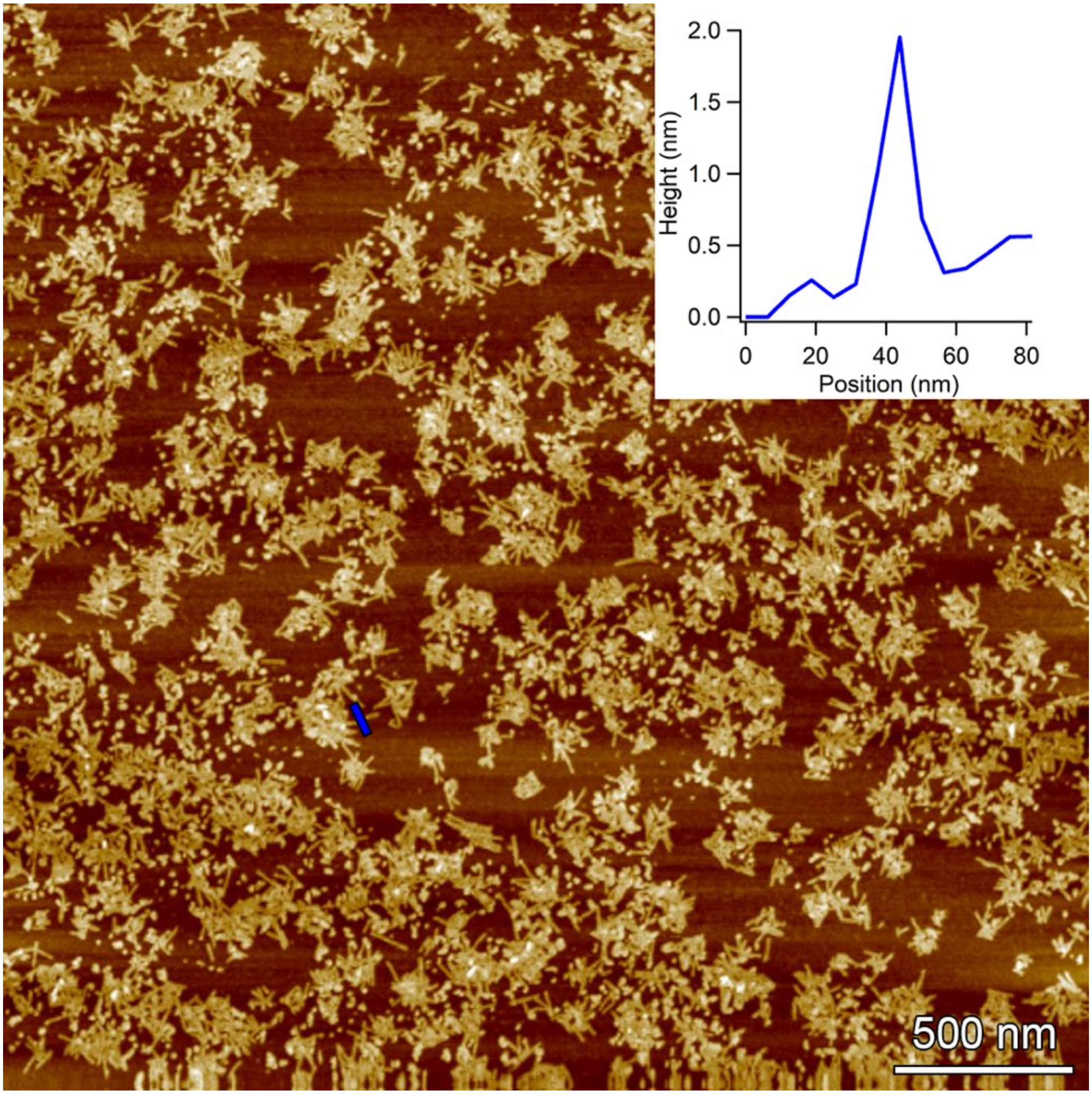

**Supplementary Figure 39.** AFM topographical image of assembly between Cy$^{HA}$-D and Mel$^{HA}$-D, 50 mM, in *N*-methylmorpholine DCl buffer, 100 mM, pD 7.5 ($D_2O$), with $MgCl_2$ 25 mM, and pD-adjusted with triethylamine and DCl. Inset shows height profile delineated by blue line in main panel.

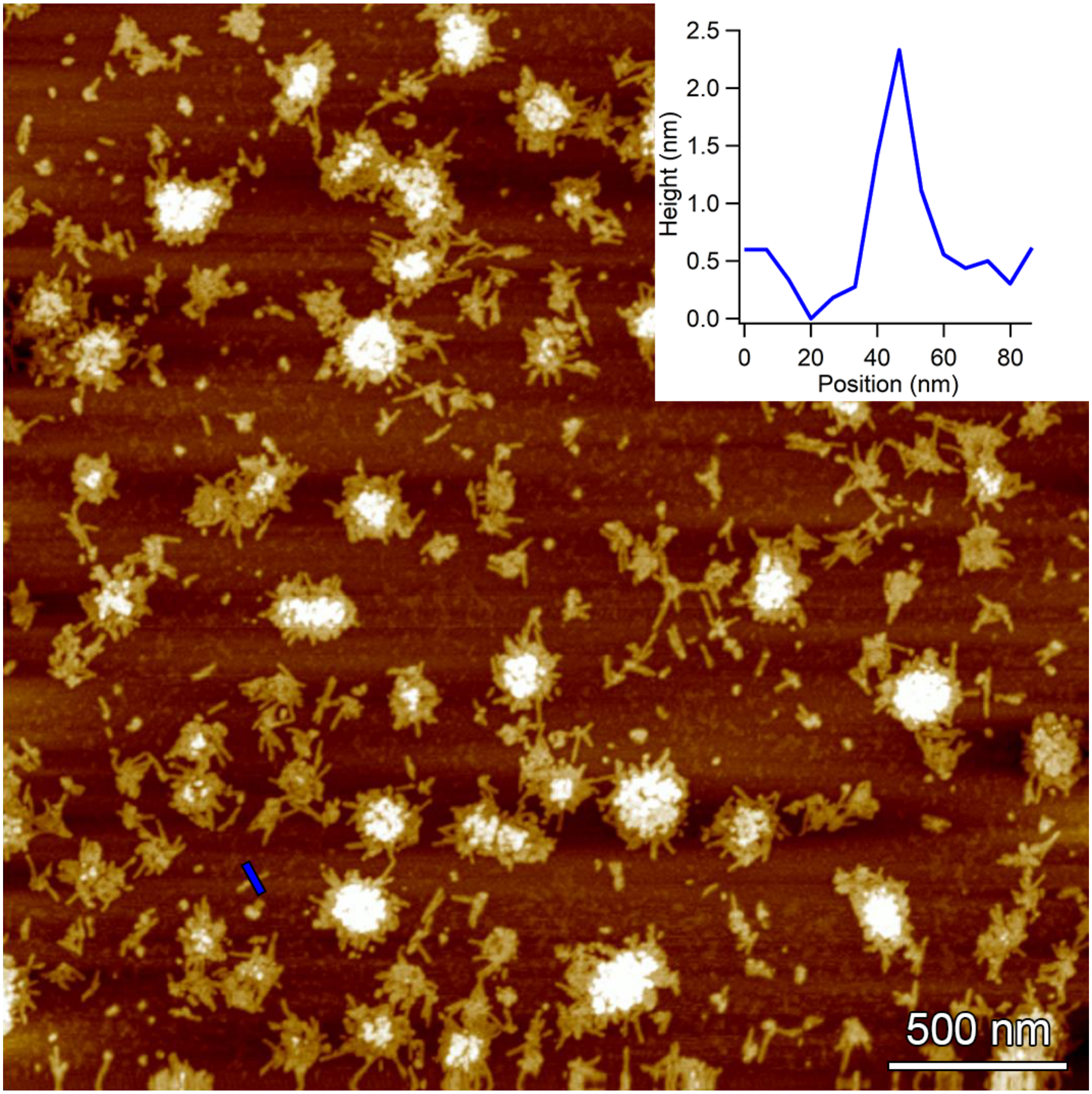

**Supplementary Figure 40.** AFM topographical image of assembly between Cy$^{HA}$-D and Mel$^{HA}$-D, 50 mM, in *N*-methylmorpholine DCl buffer, 100 mM, pD 7.5 ($D_2O$), with $MgCl_2$ 25 mM, 2-hydroxypyridine 50 mM, and pD-adjusted with triethylamine and DCl. Inset shows height profile delineated by blue line in main panel.

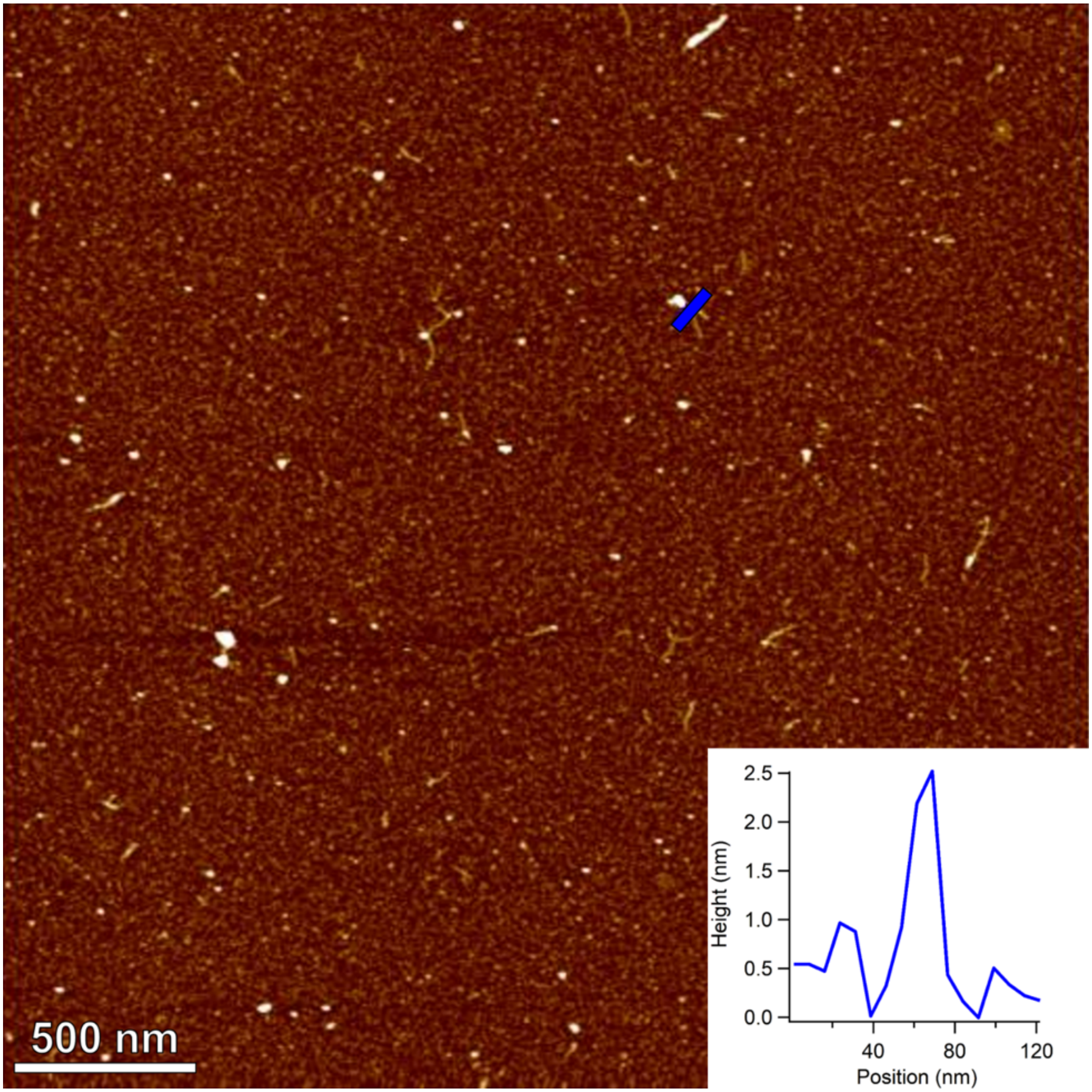

**Supplementary Figure 41.** AFM topographical image of assembly between oligo-(Cy$^{HA}$-D) and adenine, 50 mM, in lutidine DCl buffer, 200 mM, pD 6.5 ($D_2O$), with 2-hydroxypyridine 50 mM, and pD-adjusted with triethylamine and DCl. Inset shows height profile delineated by blue line in main panel.

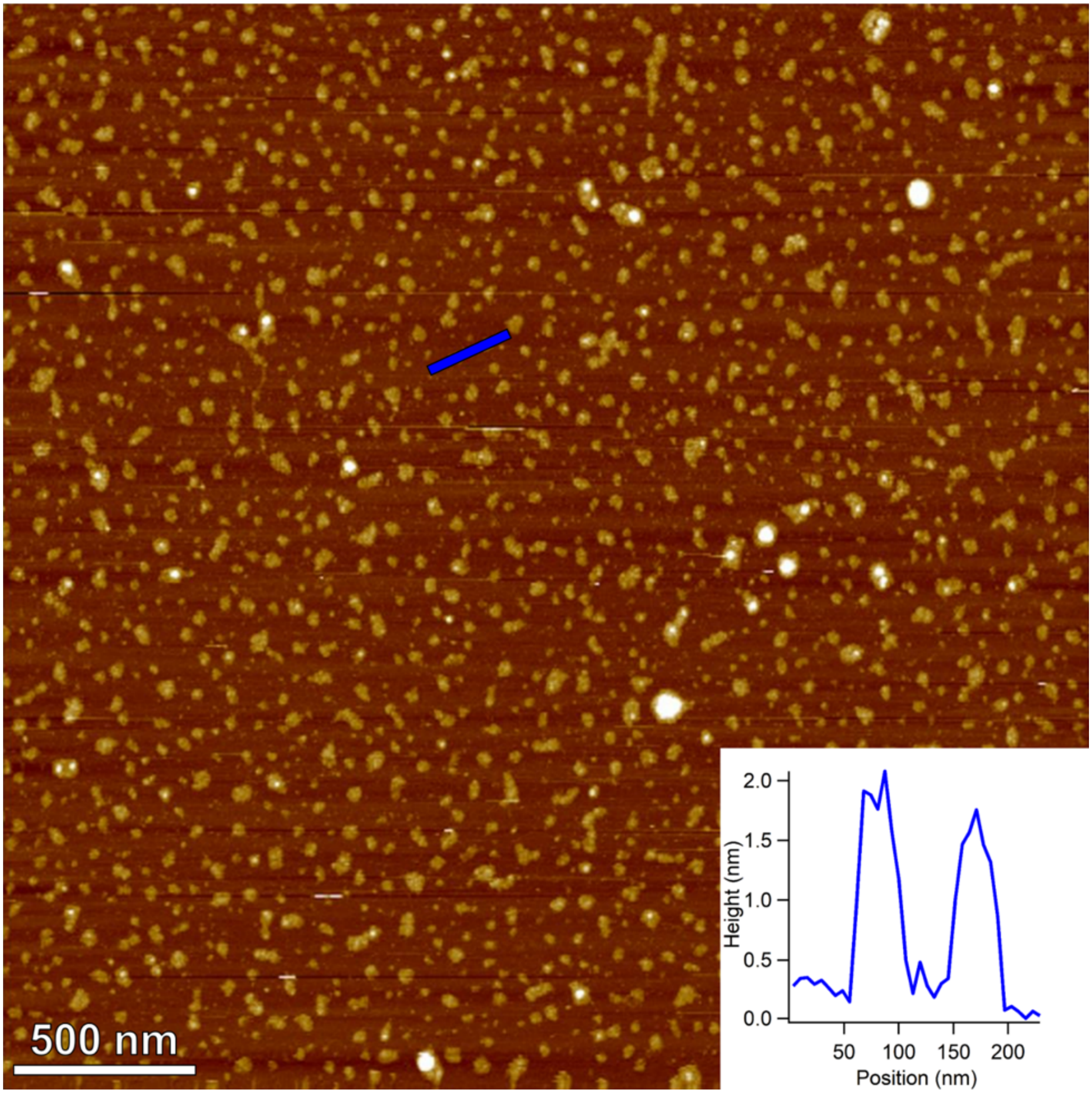

**Supplementary Figure 42.** AFM topographical image of assembly between oligo-(Ad$^{HA}$-D) and cyanuric acid, 50 mM, in lutidine DCl buffer, 200 mM, pD 6.5 ($D_2O$), with 2-hydroxypyridine 50 mM, and pD-adjusted with triethylamine and DCl. Inset shows height profile delineated by blue line in main panel.

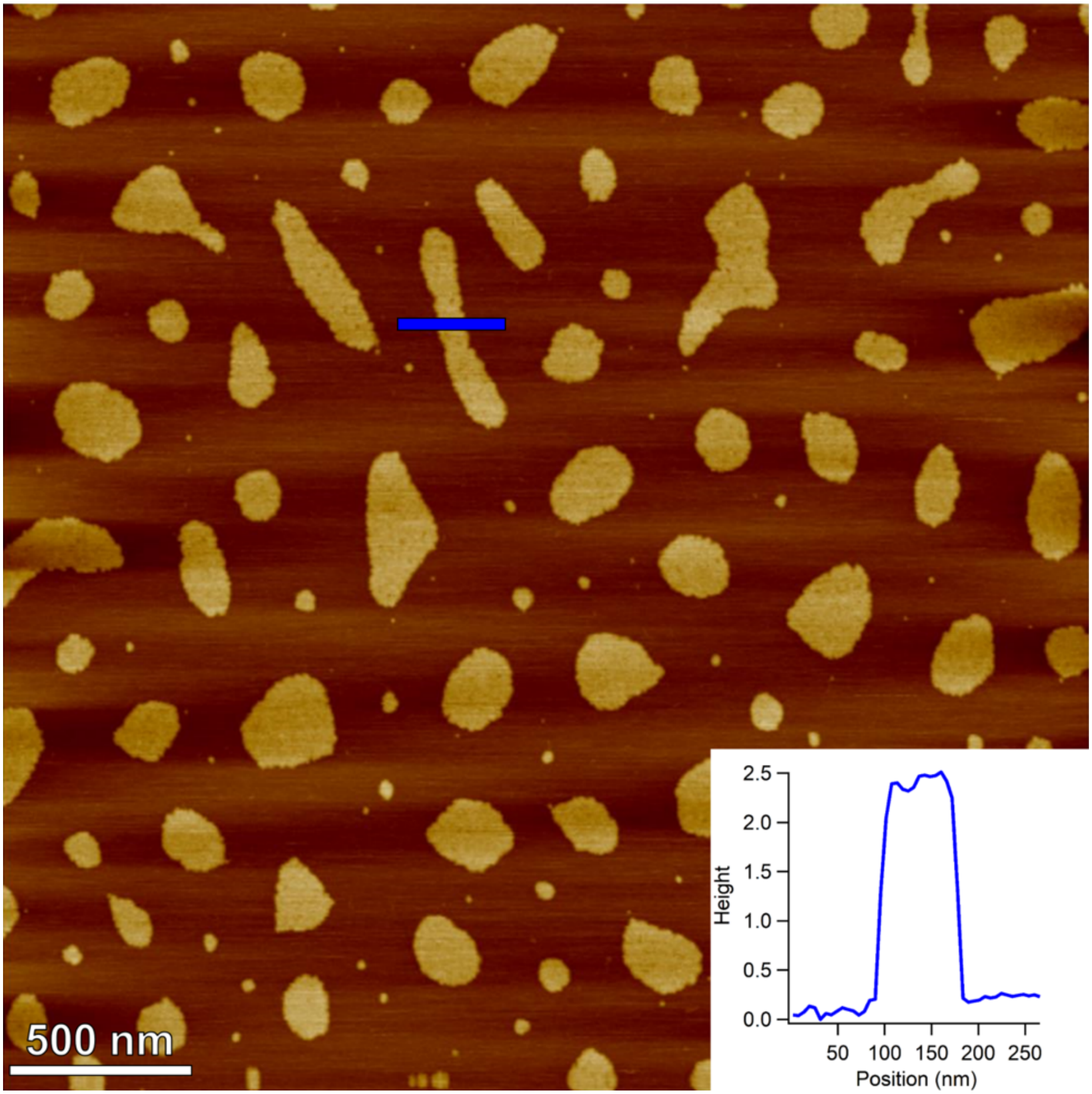

**Supplementary Figure 43.** AFM topographical image of assembly of Ad$^{HA}$-D/ Cy$^{HA}$-D co-oligomers, 50 mM, in lutidine DCl buffer, 200 mM, pD 6.5 ($D_2O$), with 2-hydroxypyridine 50 mM, and pD-adjusted with triethylamine and DCl. Inset shows height profile delineated by blue line in main panel.

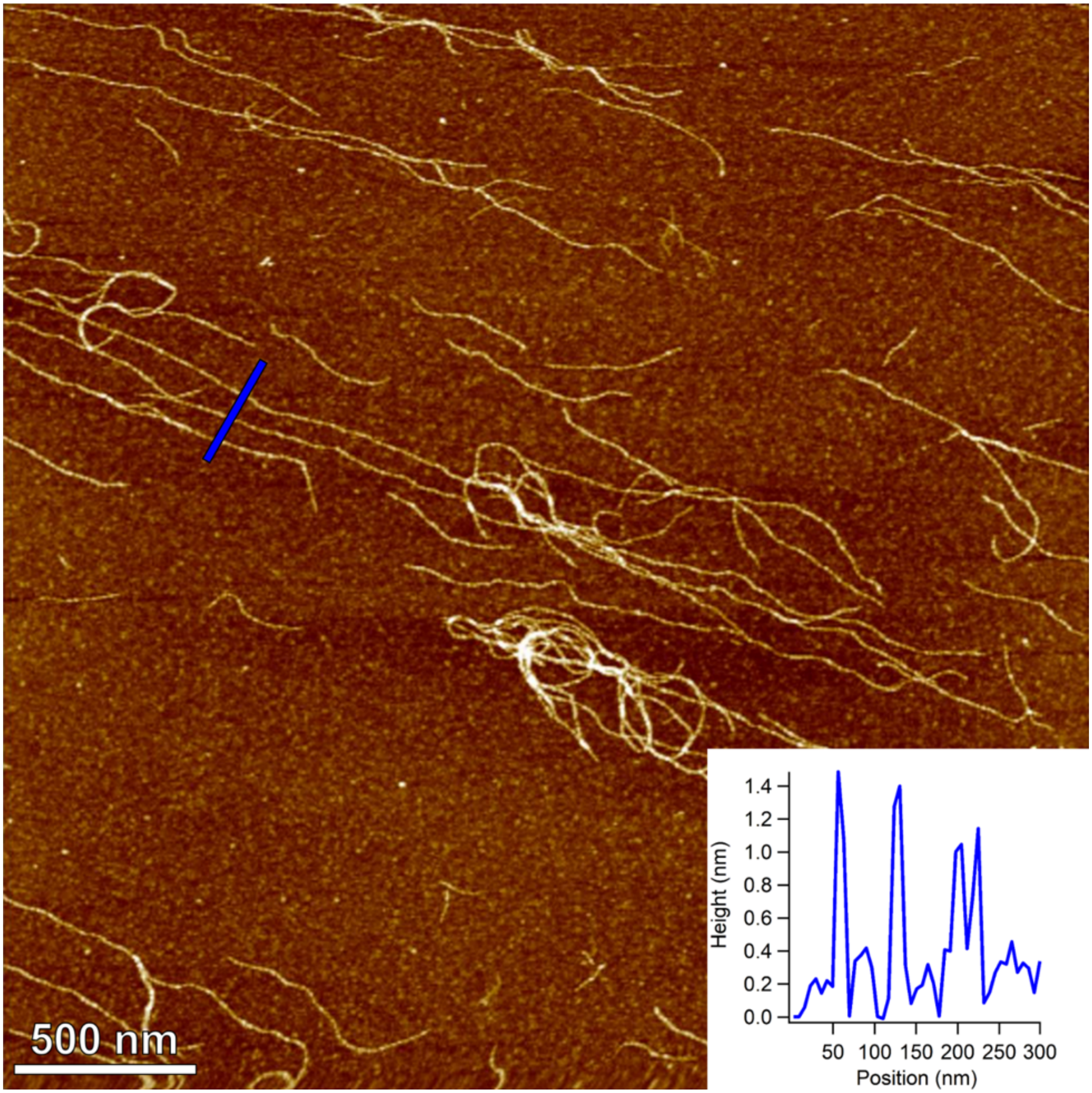

**Supplementary Figure 44.** AFM topographical image of assembly between α-(Cy$^{HA}$-D)$_2$ 25 mM and adenine 50 mM, in lutidine DCl buffer, 100 mM, pD 6.5 ($D_2O$), pD-adjusted with triethylamine and DCl. Inset shows height profile delineated by blue line in main panel.

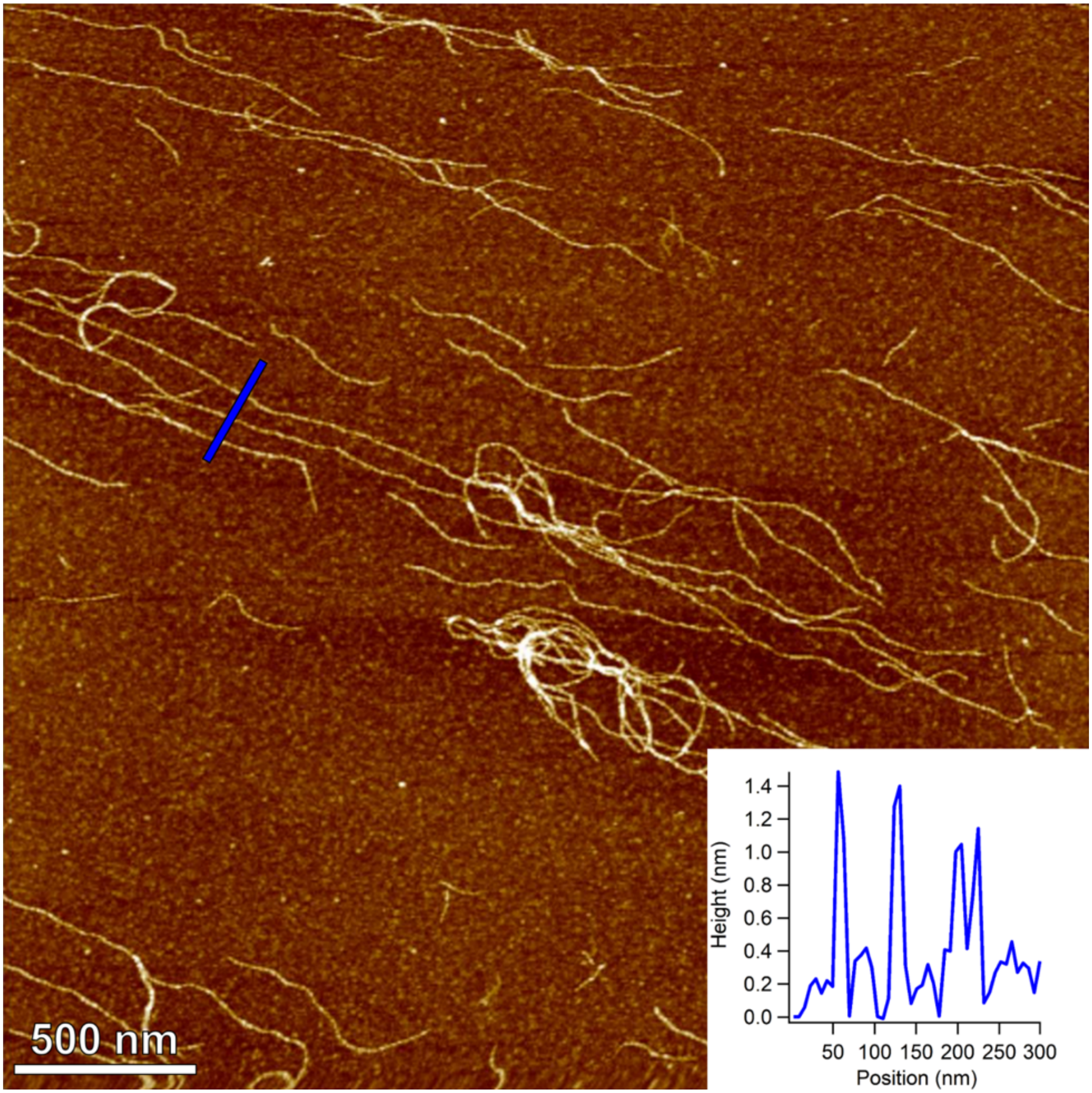

**Supplementary Figure 45.** AFM topographical image of assembly between β-(Cy$^{HA}$-D)$_2$ 25 mM and adenine 50 mM, in lutidine DCl buffer, 100 mM, pD 6.5 ($D_2O$), pD-adjusted with triethylamine and DCl. Inset shows height profile delineated by blue line in main panel.

### VI. Hydrolysis Experiments

*Sample Preparation*

Stock solutions of α-$(Cy^{HA}\text{-}D)_2$ and β-$(Cy^{HA}\text{-}D)_2$ were prepared at 20 mM in 2,6-lutidine DCl buffer pD 6.5 (prepared in $D_2O$ for comparability with NMR data). Stock solutions of melamine were prepared at 40 mM, 80 mM, and 120 mM in 2,6-lutidine DCl buffer pD 6.5. Experimental samples were prepared by combining the appropriate stock solutions in a 1:1 volume ratio in HPLC vials, and the resulting mixtures were immediately vortexed. All samples that contained melamine formed clear, viscous hydrogels within several seconds. The samples were incubated at 25 °C.

*UV-LC/MS Analysis*

Standard curves for α-$(Cy^{HA}\text{-}D)_2$, β-$(Cy^{HA}\text{-}D)_2$, and $Cy^{HA}$-D were prepared by integration of the chromatographic peaks for each species at 215 nm. The experimental samples were aliquoted directly for UV-LC/MS analysis (1 µL injection volume) without prior dilution. Concentrations were then calculated using the standard curves for each species. Because of the irreproducibility in injection volume for the experimental hydrogel samples, mass balance corrections to the calculated concentrations were made so that the reported concentrations obeyed the relation $[(Cy^{HA}\text{-}D)_2] + ½[Cy^{HA}\text{-}D] = 10$ mM. Data analysis was performed with Igor Pro 6.37. Exponential decay curves of the form $C(t) = C_0e^{-kt}$ (where C(t) equals the measured concentration at a given time, $C_0$ equals the initial concentration, and k equals the extrapolated decay constant) were fitted using the default software regression algorithm. Half-life values were calculated from the decay constants k by the equation $t_{½} = (\ln 2)/k$.

| Sample | $C_0$ (mM) | k ($d^{-1}$) | $t_{½}$ (d) |
|---|---|---|---|
| α-$(Cy^{HA}\text{-}D)_2$ 20 mM melamine 0 mM | 9.800 ± 0.144 | 0.1527 ± 0.0037 | 4.541 ± 0.109 |
| α-$(Cy^{HA}\text{-}D)_2$ 20 mM melamine 20 mM | 9.436 ± 0.217 | 0.07373 ± 0.00390 | 9.401 ± 0.472 |
| α-$(Cy^{HA}\text{-}D)_2$ 20 mM melamine 40 mM | 9.196 ± 0.182 | 0.01865 ± 0.00274 | 37.17 ± 4.76 |
| α-$(Cy^{HA}\text{-}D)_2$ 20 mM melamine 60 mM | 9.554 ± 0.098 | 0.007044 ± 0.001260 | 98.40 ± 14.9 |
| β-$(Cy^{HA}\text{-}D)_2$ 20 mM melamine 0 mM | 9.925 ± 0.021 | 0.002799 ± 0.000258 | 247.6 ± 20.9 |
| β-$(Cy^{HA}\text{-}D)_2$ 20 mM melamine 20 mM | 9.992 ± 0.028 | 0.004029 ± 0.000348 | 172.0 ± 13.7 |

**Supplementary Table 1.** Calculated initial concentration parameters (millimolar units), rate constants (inverse day units), and half-life values (day units) for the hydrolysis of α-$(Cy^{HA}\text{-}D)_2$ and β-$(Cy^{HA}\text{-}D)_2$ with varying amounts of melamine. Errors are expressed as one standard deviation.

### VII. Notes

1. Because the UV or ESI response factors for the many compounds produced in these experiments are unknown in general, yields cannot be quantified reliably.

2. In the prebiotic syntheses of nucleobase-functionalized α-hydroxy acids, extraction of the reaction mixtures with dichloromethane and subsequent hydrolysis with 6 M HCl at reflux were performed for experimental convenience and emphatically for any presumed prebiotic plausibility of these conditions. The acid-catalyzed hydrolysis of cyanohydrins under prebiotically realistic conditions to α-hydroxy carboxamides, and eventually to α-hydroxy acids, would proceed on a time scale that is too long for practical experimentation. Because strong, refluxing acid was used to hydrolyze the cyanohydrins, and because analysis of these hydrolysis reactions would be performed by $^{1}$H NMR, a dichloromethane extraction was performed in most cases as a safety measure to prevent the production of HCN from iron cyanides, and to exclude any paramagnetic iron species that would interfere with NMR analysis. Yields could not be reliably calculated from this method since the nucleobase-functionalized cyanohydrins are not efficiently extracted from water.
